# *regionalpcs*: improved discovery of DNA methylation associations with complex traits

**DOI:** 10.1101/2024.05.01.590171

**Authors:** Tiffany Eulalio, Min Woo Sun, Olivier Gevaert, Michael D. Greicius, Thomas J. Montine, Daniel Nachun, Stephen B. Montgomery

**Affiliations:** Department of Biomedical Data Science, Stanford University, Stanford, CA, 94305, USA; Department of Neurology & Neurological Sciences, Stanford University, Stanford, CA, 94305, USA; Department of Pathology, Stanford University, Stanford, CA, 94305, USA

**Keywords:** epigenetics, methylation, statistical method, quantitative trait loci

## Abstract

We have developed the regional principal components (rPCs) method, a novel approach for summarizing gene-level methylation. rPCs address the challenge of deciphering complex epigenetic mechanisms in diseases like Alzheimer’s disease (AD). In contrast to traditional averaging, rPCs leverage principal components analysis to capture complex methylation patterns across gene regions. Our method demonstrated a 54% improvement in sensitivity over averaging in simulations, offering a robust framework for identifying subtle epigenetic variations. Applying rPCs to the AD brain methylation data in ROSMAP, combined with cell type deconvolution, we uncovered 838 differentially methylated genes associated with neuritic plaque burden—significantly outperforming conventional methods. Integrating methylation quantitative trait loci (meQTL) with genome-wide association studies (GWAS) identified 17 genes with potential causal roles in AD, including *MS4A4A* and *PI-CALM*. Our approach is available in the Bioconductor package *regionalpcs*, opening avenues for research and facilitating a deeper understanding of the epigenetic landscape in complex diseases.

## Introduction

DNA methylation is a key component of the epigenome responsible for regulating gene expression, holding considerable promise as a disease biomarker and a therapeutic target. Predominantly occurring at cytosine-phosphate-guanine (CpG) sites, this epigenetic modification of DNA is influenced by both inherited genetic variation and environmental factors (1–5). Specific methylation changes have been associated with neurological disorders, notably Alzheimer’s disease (AD) (6–9), suggesting that DNA methylation may be helpful in identifying novel disease treatments. The applications of demethylating agents to treat acute myeloid leukemia and myelodysplastic syndrome are among a growing number of methylation-based therapeutics (10–13). DNA methylation changes may identify genes and pathways relevant to disease biology that are more difficult to detect with gene or protein expression because of often noisier data (14, 15).

Although many associations have been identified between DNA methylation and disease pathogenesis, interpretation of these findings remains challenging. Complex, sometimes contradictory, relationships exist between DNA methylation and gene expression. Methylation of promoters typically induces transcriptional silencing, while methylation within gene bodies can increase gene expression (16, 17). Like other functional genomics modalities, DNA methylation is also cell type-specific (15, 18). There is considerable variability in the magnitude and direction of associations reported between methylation and disease phenotypes (19).

Compounding these challenges in interpreting DNA methylation is the granularity at which it is assessed. Existing approaches typically analyze methylation at the level of single CpG sites, ranging from methylation arrays that quantify several hundred thousand CpG sites to whole genome methylation sequencing consisting of nearly 30 million CpG sites (20–22). Although many studies have investigated CpG-level changes in methylation across disease states (23–27), the functional implications of changes in methylation of individual CpGs often remain ambiguous. Most studies map CpG sites of interest to the nearest genes to interpret their biological impact (23, 28, 29). However, this approach does not account for the cumulative and regional effects of changes in multiple CpG sites on gene function and disease pathology. Some studies try to model regional changes in methylation by systematically segmenting the genome into regions spanning 100-1000 base pairs to identify differentially methylated regions (DMRs) (30, 31). One intrinsic limitation of DMR-based approaches is that they depend on arbitrary parameters such as region size, whose values can substantially affect the results in ways that may be difficult to interpret biologically (32).

An alternative solution to improve interpretability is to summarize methylation at the level of specific genomic regions, such as promoters or CpG islands. Research by Cai *et al*. demonstrated that methylation aggregated at the gene level more effectively discriminated between disease groups than individual CpG-level measurements (33). While averaging across CpG sites within these regions is the most common aggregation strategy (34–38), this approach oversimplifies the often more complex correlation structures between CpG sites across a region. Kapourani and Sanguinetti underscored this by demonstrating that large genomic regions (+/- 7kb around the transcription start site) exhibit complex methylation patterns that influence gene expression and that such phenomena were not adequately captured by averaging CpG-level methylation across the region (39). Zheng *et al*. further demonstrated that gene-region methylation profiles can delineate disease subgroups, emphasizing the nuanced complexity of methylation at the gene level (40). Thus, there is a need for an effective method of capturing methylation signals within gene regions to improve the detection of biologically relevant methylation changes.

We have developed the regional principal components (rPCs) method for aggregating methylation data to improve gene region methylation summaries. This method leverages principal components analysis (PCA) within genomic regions to capture methylation changes pertinent to disease with increased accuracy. By capturing multiple orthogonal axes of variance across CpG sites, rPCs offer a robust representation of methylation data. To facilitate its broad application in the research community, we have made our software available as an R package, *regionalpcs*, available on Bioconductor (41) and GitHub at https://github.com/tyeulalio/regionalpcs.

Our study used *regionalpcs* to identify cell type-specific methylation changes associated with AD and genetic variation within the ROSMAP cohort (7). AD is a neurodegenerative disorder that affects approximately 50 million people globally and is marked by progressive memory loss and cognitive decline (42–44). This condition severely diminishes patient quality of life, overburdens caregivers, and exhausts healthcare infrastructures. These issues highlight an urgent need for effective interventions. Presently, the absence of definitive prevention or treatment strategies underscores a gap in our comprehension of the underlying mechanisms driving AD. Research implicates DNA methylation changes as a contributor to AD pathology, with hundreds of regions identified as differentially methylated in the context of the disease (45). The ROSMAP cohort comprising clinical, lifestyle, genetic, and methylation data from over 3,200 older adult participants offers an invaluable resource for investigating the epigenetic landscape of AD, which can facilitate insights into the disease’s complex etiology.

We demonstrate a comprehensive AD methylation analysis by applying *regionalpcs* in the ROSMAP cohort. Differential methylation analysis was used to discern complex links between DNA methylation and AD phenotypes. Next, we mapped quantitative trait loci (QTL) to identify genomic regions where genetic variants affect DNA methylation. We then associated those variants with the genetic risk of AD by integrating AD GWAS using colocalization (46), instrumental variables analysis (PTWAS) (47), and the probabilistic integration of these analyses (INTACT) (48). Across these analyses, *regionalpcs* revealed cell type-specific epigenetic changes associated with AD pathology and disease risk that would not be captured by CpGs or averages alone. Regional PCs balance the reduced multiple testing burden and greater interpretability of regional aggregation of methylation with the granularity of CpG-level analysis. Our method broadly applies to any analysis associating methylation with phenotypes or mapping methylation QTLs. We anticipate that this innovative approach will improve the identification of novel treatment targets and disease risk prediction.

## Results

### Refining methylation analysis through gene-level summaries

We developed the *regionalpcs* package to summarize methylation over genomic regions using PCA (Figure 1a). We chose to use PCA because it is a simple, computationally efficient approach with a well-developed theoretical framework. The rPCs we estimate summarize the highly correlated CpG features into a new set of orthogonal features that capture more information about methylation in a region than averaging but still provide a low dimensional representation of the data for downstream analyses.

**Fig. 1.**
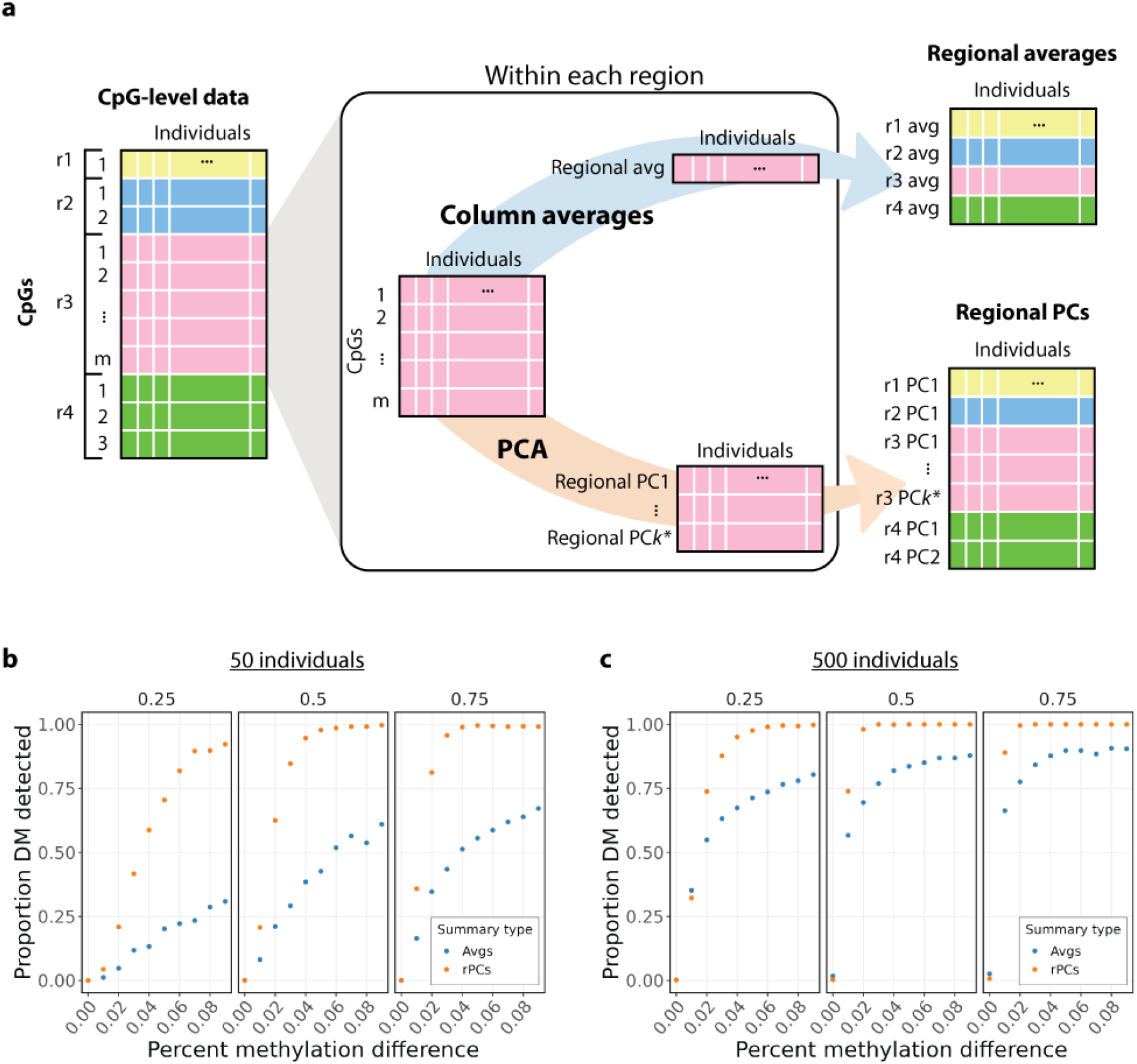
Overview of the *regionalpcs* method. a) Summarizing regional methylation using averages and regional principal components (rPCs). The CpG-level data contains normalized methylation for CpGs (rows) and individuals (columns) grouped into regions based on genomic annotations (represented by colors). For regional averages, mean methylation across CpGs in each region is calculated for each individual. rPCs are using PCA with the number of PCs (*k*^*∗*^) determined by the Gavish-Donoho method, selecting the top PCs for regional representation. b) and c) show comparative performance of summarizing regional methylation using rPCs (orange) and averages (blue) in simulations for b) 50 individuals and c) 500 individuals, highlighting the impact of larger sample sizes on methylation detection. Facets show the proportion of differentially methylated CpG sites. Each point indicates the proportion of regions (out of 1,000 simulated) identified with significant differential methylation (y-axis) at varying percent methylation differences between cases and controls (x-axis). Simulations involved 50 CpG sites per region and samples evenly split between cases and controls. Facets show the proportion of differentially methylated CpG sites.

A key advantage of using PCA for summarization over other dimensionality reduction methods is that there are well-established methods in the field of random matrix theory for identifying how many eigenvectors capture a distinguishable signal from random noise. Using a principled approach to determine the number of components to summarize a region makes this approach feasible when performing this summary over many thousands of regions. We used the Gavish-Donoho method (49), designed to identify the optimal eigen-value threshold for minimizing the asymptotic mean squared reconstruction error of the original data. Including too few eigenvectors will lead to suboptimal reconstruction of the data because some axes of variation representing genuine signal have been ignored. However, including too many eigenvectors with very small eigenvalues will introduce random noise that increases the reconstruction error. We also allow users to use the less conservative Marchenko-Pastur method as an alternative approach to choosing the number of components (50).

We chose to summarize methylation at the level of gene regions. This gene-level methylation allows for a more straightforward interpretation of biological pathways—context often lost when focusing on individual CpG sites. Our framework can accommodate a variety of regional annotations, such as promoters, CpG islands, intergenic enhancers, or any custom regions specified by the user.

### rPCs detect more subtle methylation differences in simulated data

We evaluated the effectiveness of rPCs for identifying differential methylation using the simulation framework *RRBS-sim* (51) to generate artificial reduced representation bisulfite sequencing (RRBS) data. This type of data can be seen as an intermediary between the sparse coverage of methylation microarrays and the denser coverage of whole genome bisulfite sequencing.

Our analysis aimed to compare the summarization of methylation using *regionalpcs* against traditional averaging to identify methylation changes. Averages were computed as the mean methylation across all CpGs within a region, while rPCs were derived through PCA of CpG-level methylation (Figure 1a).

We found that rPCs outperform averaging in identifying differentially methylated (DM) regions. We simulated 1,000 regions, each with 50 CpG sites and 50 individuals (split evenly between cases and controls) and varied the extent of differential methylation and the proportion of DM sites. The rPCs detected more DM regions, even when the proportion of DM sites was as low as 25%, detecting a median of 73.1% of DM regions, compared to just 19.1% with averages. This performance gap lessened slightly when the proportion of DM sites increased to 75%, with rPCs identifying 99% of the cases, compared to a 57.4% detection rate with averages (Figure 1b).

The magnitude of methylation differences between cases and controls was a critical factor in DM detection. For a modest 1% difference in methylation, rPCs successfully detected DM in 18.8% of regions, doubling the performance of averaging, which detected 8.4%. When the methylation difference was increased to 9%, the performance of rPCs identified 99.7% of DM regions, in contrast to the 50.1% detection rate with averages.

The number of samples in the simulation also influenced DM detection, particularly for the averaging method. With a sample size of 50, rPCs detected a median of 94.4% of DM regions across simulation parameters, compared to 32.6% with averaging. Increasing the sample size to 500 improved the detection capability of both methods, with rPCs detecting a median of 99.9% of DM regions, compared to 80.4% with averages (Figure 1c). This finding demonstrates the improved sensitivity of rPCs, especially in studies with smaller sample sizes.

Our analysis showed that the number of CpG sites per re-gion had a more subtle effect on DM detection. In regions with 20 CpG sites, representative of RRBS data in promoter regions, rPCs detected a median of 78.2% of DM regions compared to only 45.4% with averaging. In regions with 50 CpG sites, representing full gene regions in RRBS data, the detection rate was 99.0% for rPCs compared to 59.1% for averages.

In addition to increased sensitivity, rPCs have comparable specificity to averaging. In a matrix comprising 700 non-DM regions and 300 DM regions with varying methylation differences, rPCs accurately identified all non-DM regions and a median of 100% of DM regions, compared to averaging, which only identified 86.7% of DM regions (Supplementary Figure 1, Supplementary Table 1). These findings highlight that rPCs improve the detection rate of DM regions without incurring additional false positives.

### Applications to Alzheimer’s Disease

We used the *regionalpcs* package to summarize methylation in an AD cohort to demonstrate the value of our tool in a real dataset. We analyzed paired DNA methylation and whole genome sequencing data from a subset of 563 older individuals from the ROSMAP cohort (7, 52, 53). Data were obtained post-mortem from the dorsolateral prefrontal cortex (DLPFC). The cohort’s median age at death was 88.5 years, with females comprising 63.2%, and all participants reporting European ancestry.

Individuals were classified through annual and post-mortem clinical assessments into Alzheimer’s Disease Dementia (ADD, 42.8%), Mild Cognitive Impairment (MCI, 25.4%), No Cognitive Impairment (NCI, 30.2%), or Other Dementia (1.6%) categories (Supplementary Table 2). Neuropathological indices for neurofibrillary tangles (54) and neuropathological diagnoses based on CERAD scores for neuritic plaques were also recorded (55). Braak staging, quantifies neurofibrillary tangle extent throughout the brain. Approximately 80% of the cohort had tangles extending into limbic regions (Braak stage III to VI), while only 24% had tangles in neocortical regions, which include the DLPFC (Braak stages V and VI). The CERAD score, assessing neuritic plaque density in the neocortex, was used to diagnose individuals as Definite, Probable, Possible, or No AD. Neuropathologic diagnosis from CERAD score showed 24.7% of the cohort had no AD, contrasting with 30.2% diagnosed with definite AD.

In analyzing the ROSMAP data, we applied *regionalpcs* to identify methylation changes relevant to AD (Figure 2). Starting with DNA methylation data from bulk tissue of the DLPFC, we utilized cell type deconvolution to isolate cell type-specific signals (Figure 2a-b). Subsequently, we summarized this methylation data across various gene regions (Figure 2c-d). Our goal was to assess how methylation summarization at the gene level using *regionalpcs* compares against traditional averaging and single CpG analysis in detecting changes linked to neuritic plaque burden (Figure 2e). Moreover, we explored the effectiveness of *regionalpcs* versus averages for mapping meQTLs and integrating these findings with AD GWAS data (Figure 2f-g).

**Fig. 2.**
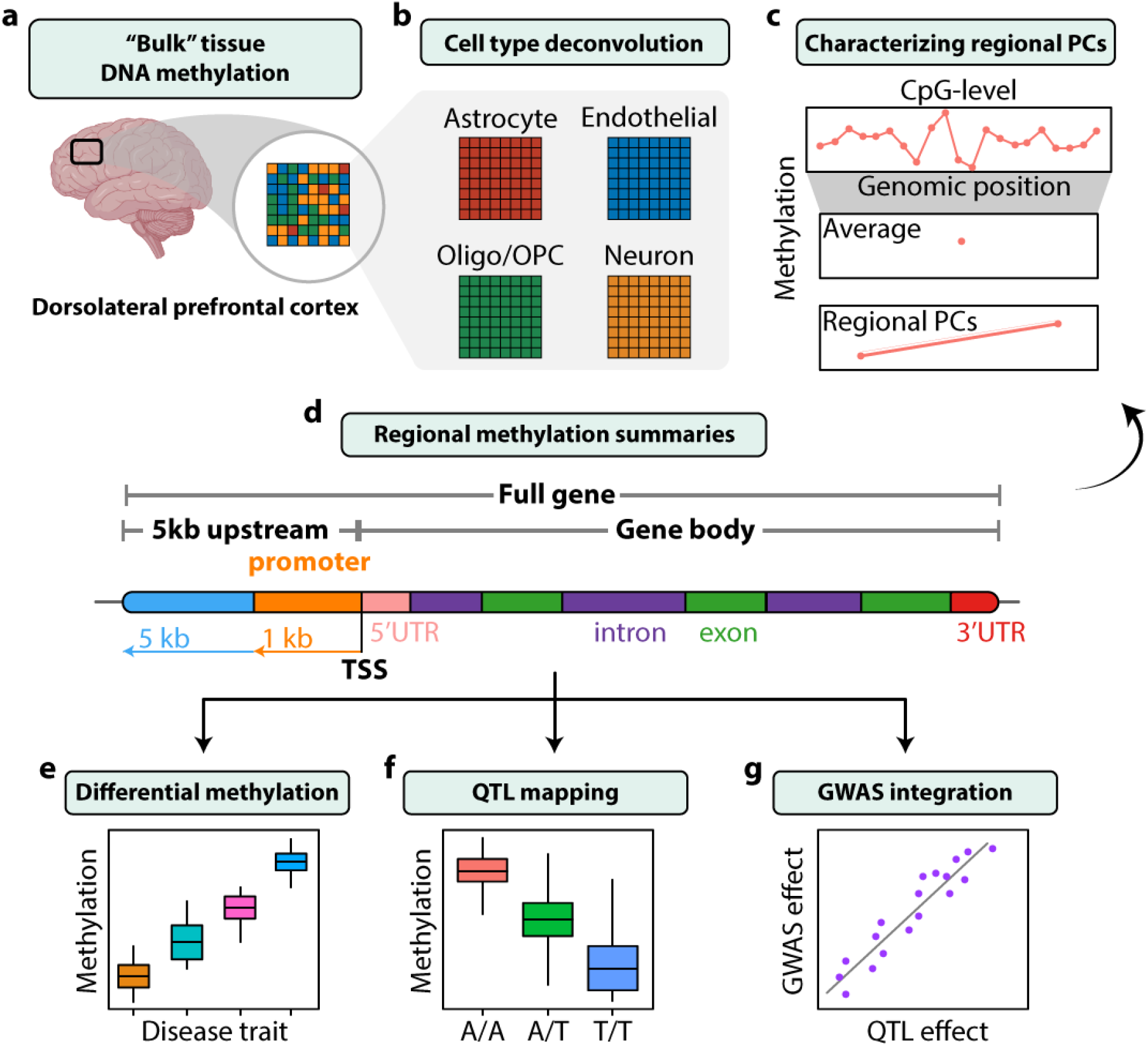
Methodological overview of DNA methylation analysis in AD using the ROSMAP cohort. a) Data consisted of DNA methylation from the dorsolateral prefrontal cortex obtained from the ROSMAP study. The bulk methylation data, indicative of a heterogeneous mixture of cell types, is visually portrayed. b) Cell type deconvolution isolated cell-type specific signals from the bulk methylation data, resulting in four datasets to study cell type-specific methylation changes in Alzheimer’s disease. c) Regional CpG-level methylation data was summarized using averaging and *regionalpcs*. Each summary type is depicted here for an example gene, with methylation levels and features plotted on the y-axis and x-axis, respectively. d) Analysis focused on four gene regions: promoters, gene body, 5kb upstream, and the full gene, using comprehensively annotated regions to study methylation on multiple scales e) Performed differential methylation analysis to detect changes in gene-level methylation across disease states and various phenotype traits within our cohort. A toy example is shown in the boxplot, with methylation levels (y-axis) increasing across the disease trait groups (x-axis). f) Mapped quantitative trait loci (QTL) using the regional methylation summaries to identify genes with methylation levels associated with genomic variants. A toy example of a QTL is shown in the boxplot, where methylation levels (y-axis) change based on the genotype (x-axis) across individuals. g) Integrated genome-wide association study (GWAS) results with methylation QTLs to identify genetic variants with shared signals between methylation changes and Alzheimer’s disease susceptibility. A toy example is shown in the scatter plot with a positive correlation between QTL and GWAS effects.

### Cell type deconvolution to impute cell type-specific DNA methylation from bulk tissue

Before applying the *regionalpcs* method on the AD cohort, we addressed the issue of mixed cell type signals in the methylation data. Previous investigations into differential methylation and methylation QTLs have primarily used bulk tissue samples consisting of a heterogeneous mixture of cell types (7, 56–58). Although these studies have yielded valuable insights, attributing the changes in methylation to cell type proportions or cell type specificity is challenging (59). Cell type heterogeneity often acts as a confounding factor in functional genomics data collected from bulk tissue samples (60, 61).

To better understand the complexities of DNA methylation and its association with disease phenotypes, we refined our approach by implementing cell type deconvolution. This approach facilitated the imputation of cell type-specific methylation profiles from the ROSMAP cohort’s bulk tissue samples. We estimated methylation levels for the predominant brain cell types: neurons, astrocytes, endothelial cells, and a combined group of oligodendrocytes and oligodendrocyte progenitor cells (oligo/OPCs). These estimates improved upon the common practice of adjusting for cell type composition by either removing its effects or including it as a covariate in downstream analyses.

We leveraged *EpiSCORE* (62) to estimate the proportions of each cell type. Aligning with existing literature (63–65), neurons were the most abundant cell type, followed by astrocytes, oligodendrocyte/OPCs, and endothelial cells. Microglia were not detected, likely due to their relatively low abundance in the DLPFC brain region (64). Our subsequent analyses focused on the four cell types with reliably estimated proportions (Figure 3a). Validation with an alternative method (66) for estimating neuron proportions demonstrated high consistency (Pearson *r* = 0.8, *p*-value = 1.2 *×* 10^*−*128^; Supplementary Figure 2).

**Fig. 3.**
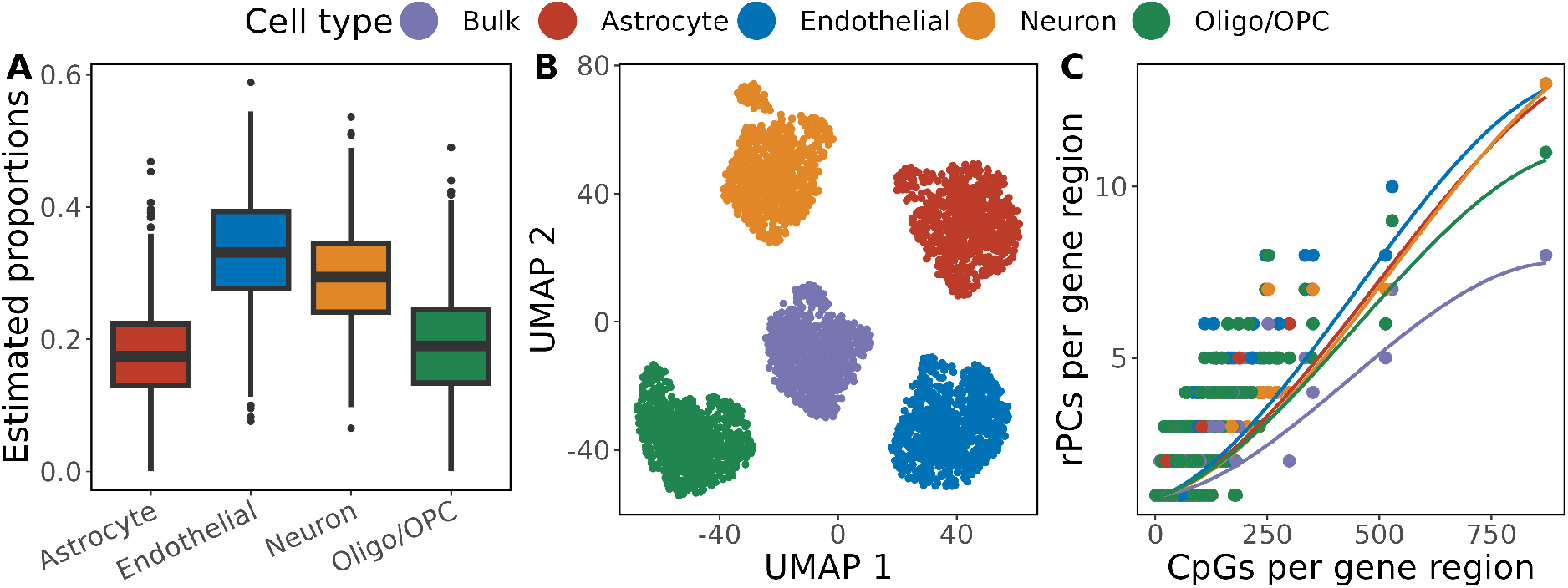
Cell type proportions and rPCs in brain cells. a) Boxplots represent the estimated cell type proportion distributions across four primary brain cell groups: astrocytes, endothelial cells, neurons, and oligodendrocytes/oligodendrocyte progenitor cells (Oligo/OPC). Horizontal lines within the interquartile range (IQR) boxes indicate the median values for each group, whiskers represent a distance of 1.5*IQR from the IQR box. b) UMAP visualization of deconvolved cell type-specific data across all samples. Bulk tissue samples occupy a central position, with samples clustering by cell type. c) The scatter plot depicts the positive association between the number of CpGs and the number of regional PCs (rPCs) representing the gene regions, colored by cell type. LOESS lines depict data trends for each cell type, showing that the number of rPCs starts to level off as the number of CpGs exceeds 750.

Applying *Tensor Composition Analysis (TCA)* (59), we decomposed the bulk methylation data into cell type-specific profiles. This step mirrors the resolution achievable through cell sorting techniques like fluorescence-activated cell sorting (FACS), which isolates individual cell populations before assaying methylation. Uniform Manifold Approximation and Projection (UMAP) analysis (67) showed distinct clustering of bulk and cell type-specific methylation profiles (Figure 3b). K-means clustering analysis (68) showed higher mean Silhouette scores for cell type-specific profile clusters compared to the bulk, indicating more homogenous cell type-specific clusters (astrocytes = 0.85, endothelial cells = 0.85, neurons = 0.79, oligo/OPC = 0.78, bulk cells = 0.67).

Clustering analysis validated that our deconvolved methylation profiles clustered closely to a set of previously published nuclei-sorted methylation profiles from matching cell types (26). 45 of 53 nuclei-sorted neuron samples clustered with our deconvolved neurons and 40 of 42 nucleisorted oligodendrocyte samples clustered with our deconvolved oligodendrocyte/OPCs (Supplementary Table 3).

### Summarizing cell type-specific brain methylation using *regionalpcs*

To better understand the complexity of cell type-specific methylation in the brain, we used our novel *regionalpcs* package to summarize methylation signals across 16,417 protein-coding genes. Our primary analysis focused on the full gene region, extending from 5kb upstream of the transcription start site (TSS) to the end of the 3’ untranslated region (Figure 2d). Additionally, we examined three sub-regions—promoters, 5kb upstream (of TSS), and gene bodies (in Supplement). The count of CpGs per gene varied widely, with a median of 15 CpGs and a range from one to 872 CpGs (Supplementary Table 4). Each gene was represented by one average value and a varying number of rPCs, ranging between one and thirteen. This approach yielded an average of 1.04 rPCs per gene, explaining a median of 15% variance across genes (Supplementary Table 4).

Our analysis confirmed that rPCs primarily reflected regional methylation signals rather than global factors like batch, sex, or age. We observed consistent correlation distributions among rPCs of the same rank using a correlation analysis of rPCs spanning multiple genes (see Methods). Briefly, we calculated the correlation distribution among rPCs of the same rank. If specific rPCs mainly reflected global signals, we would expect different distribution patterns across rPC ranks 1 through 3 (an illustrative example is provided in Supplementary Figure 3a). Following rigorous data refinement, we observed minimal correlation among all rPC ranks and no notable differences between them (Supplementary Figure 3d). These findings support the notion that our rPCs represent region-specific signals.

The number of rPCs positively correlated with gene region complexity, showing that rPCs adeptly capture the multifaceted aspects of methylation signals within gene regions. The region complexity comprised the number of CpGs within a gene, length, and signal variance. Both the count of CpGs and gene length consistently showed positive correlations with the number of rPCs for each gene region across all cell types, achieving median Pearson correlation values of 0.74 and 0.12, respectively, both having *p*-values < 0.0001 (Figure 3d, Supplementary Table 5, additional region types in Supplementary Figure 4). The bulk data yielded the fewest rPCs, suggesting that cell type deconvolution aids in distinguishing methylation signals from noise. Variance in CpG methylation within gene regions, measured by median absolute deviation, also correlated positively with rPC count across all cell types (median Pearson *r* = 0.10, *p <* 0.0001).

By summarizing at the gene level, we substantially reduced the features from 271,223 CpGs to 17,341 rPCs and 16,417 averages for endothelial cells, with similar reductions for other cell types (Supplementary Table 4). This approach efficiently reduced multiple testing challenges and computational demands.

### *regionalpcs* identify more disease-relevant differential methylation in AD

Our comparative differential methylation (DM) analysis utilized *regionalpcs* to identify more disease-relevant methylation changes than traditional averaging or CpG-level assessments. We found notable DM associated with AD, mainly linked to neocortical neuritic plaques (CERAD score) and neurofibrillary degeneration (Braak stage). Interestingly, fewer associations were observed with *APOE* genotype and ADD diagnosis, suggesting that methylation endophenotypes might offer clearer biolog-ical distinctions than these clinical and genotype phenotypes alone (Supplementary Figure 5).

For our comparison of methylation summary methods, we focused on the proportion of significantly differentially methylated features identified by each method. This approach accounts for the variability in the number of CpGs and rPCs representing each gene, in contrast to using a single average value per gene (Figure 2c). Across all cell types, rPCs consistently identified a significantly higher proportion of differentially methylated features associated with neuritic plaque density to both averages and individual CpGs (Supplementary Figure 6). The proportion of features identified by rPCs was a median of four times greater than the proportion identified with averages and 24 times greater than CpGs (Supplementary Table 6), highlighting their greater power to capture methylation changes associated with neuritic plaque load.

At the gene level, our analysis highlighted an improved ability of rPCs to detect differential methylation. rPCs identified a more extensive set of differentially methylated genes than averages across all cell types. In astrocytes, rPCs identified 638 differentially methylated genes versus the 192 detected by averages (Supplementary Figure 5, Supplementary Table 7). Astrocytes were the cell type with the largest number of associated genes for both neuritic plaques and tau, supporting previous studies underlining their critical role in AD (69–73). Differentially methylated genes associated with neuritic plaques were also found in endothelial cells, neurons, and oligodendrocytes/OPCs (Figure 4a). In contrast, analysis of bulk data identified few DM genes—10 using rPCs and none using averages—highlighting the inherent challenge in discerning methylation associations within mixed cell populations. Most genes (62%) associated with neuritic plaque burden were exclusive to a single cell type, with astrocytes having the greatest number of unique genes (125 with averages, 324 with rPCs).

**Fig. 4.**
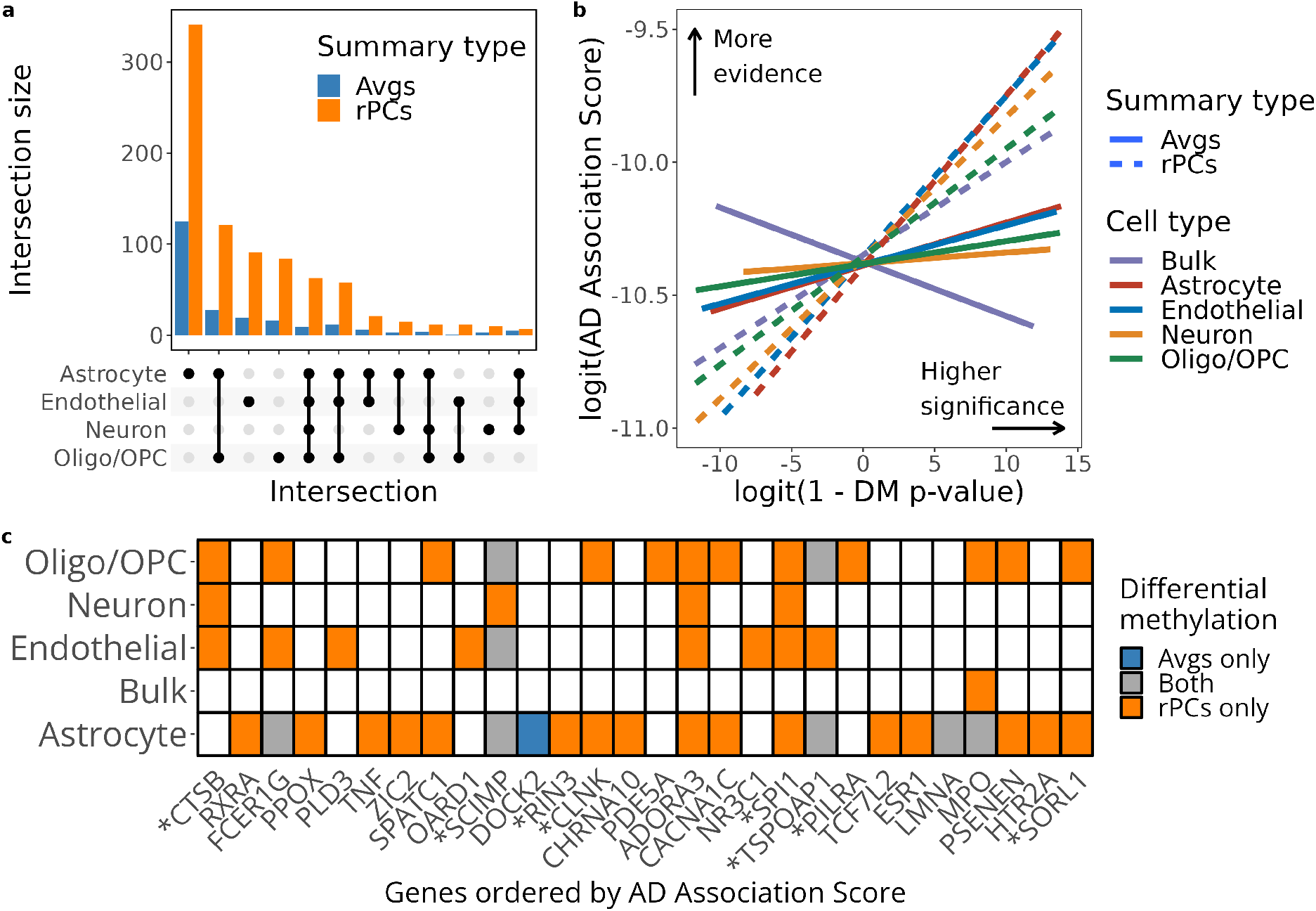
Cell type-specific gene methylation associations with AD risk. a) An UpSet plot visualizing the intersections of neuritic plaque-associated gene methylation across different cell types. The plot emphasizes the cell type-specific gene associations, with a substantial number of associated genes being unique to individual cell types, particularly astrocytes. We can also see the utility of rPCs in unveiling these cell type-specific patterns. b) A line plot illustrating the correlation between differential methylation significance levels and AD association scores, using dashed lines for rPCs and solid lines for averages, color-coded by cell types. The plot emphasizes the significantly stronger correlation established by rPCs, accentuating their enhanced proficiency in pinpointing well-established AD risk genes. c) A tile plot presenting genes with high evidence for AD association, based on the top 5% of AD association scores derived from the Open Targets AD gene list. The color-coded tiles indicate genes identified as having neuritic plaque-associated methylation (Benjamini-Hochberg corrected *p*-value less than 0.05) by each summary type: purple tiles for rPCs only, brown tiles for averages, and grey tiles for genes identified by both methods. Notably, rPCs exclusively identified the majority of high-evidence genes, further validating their utility in identifying disease-relevant differential methylation patterns.

We observed that many differentially methylated genes associated with neuritic plaques were identified by specific regional summaries rather than across all region types (Supplementary Figure 7). While the full gene region is composed of the other region types, several genes were identified only at the level of sub-regions and not in the full gene, while others were exclusive to full genes. This observation demonstrates that each region type may offer unique insights into DNA methylation patterns, emphasizing the importance of exploring DNA methylation at multiple scales to understand its relationship with hallmark neuropathologic features comprehensively.

To assess the relevance of these differentially methylated genes to AD, we used the AD association score from the Open Targets database (74). This score reflects the level of evidence for each gene’s involvement in AD based on previously published research (See Methods). Linear regression models correlated these scores with *p*-values from differential methylation analysis using either rPCs or averages. We assumed that genes strongly linked to AD would show more significant methylation changes, expecting a positive correlation between the AD association scores and differential methylation *p*-values. Indeed, we observed such positive correlations across all cell types, except for one, using both rPCs and averages (Figure 4b). However, when summarized using averages, bulk tissue methylation *p*-values were negatively correlated with AD association scores. A linear interaction model revealed a significantly stronger correlation using rPCs than averages (*p*-value = 4.2 *×* 10^*−*6^), suggesting that rPCs more effectively identify genes known to be associated with AD. Specifically, out of the top 5 percent of genes ranked by AD association score (348 genes), 41 genes were identified exclusively with rPCs, compared to just one by averages alone; seven genes were identified by both methods (Figure 4c). Among these, eight genes recognized by the National Institute on Aging Genetics of Alzheimer’s Disease (NIAGADS) (https://adsp.niagads.org/gvc-top-hits-list/), such as *SORL1, PILRA*, and *PSENEN*, were identified by rPCs. Notably, the *PSENEN* gene (75) showed a significant association with neuritic plaque load only when analyzed with rPCs (Benjamini-Hochberg (BH) *p*-value = 0.007), whereas no significance was found with averages (BH *p*-value = 0.521) (Supplementary Figure 8).

### rPCs detect more gene-level methylation QTLs

Next, we mapped methylation quantitative trait loci (meQTL) using averages and rPCs to study their ability to capture genetic effects on gene-level DNA methylation across 16,417 genes. Focusing on genomic variants within one megabase (Mb) of the transcription start site, we observed an advantage of rPCs over averages, detecting approximately 15% more genes with significant associations with at least one genetic variant (meGenes, Figure 5). This increase translated to a median increase of 1,608 meGenes identified by rPCs across different cell types. A substantial proportion of the genes evaluated were meGenes—75% for rPCs and 66% for averages. There was minimal evidence of cell type specificity of meGenes using averages or rPCs across all region types because most genes were meGenes across all cell types (Supplementary Figure 9).

**Fig. 5.**
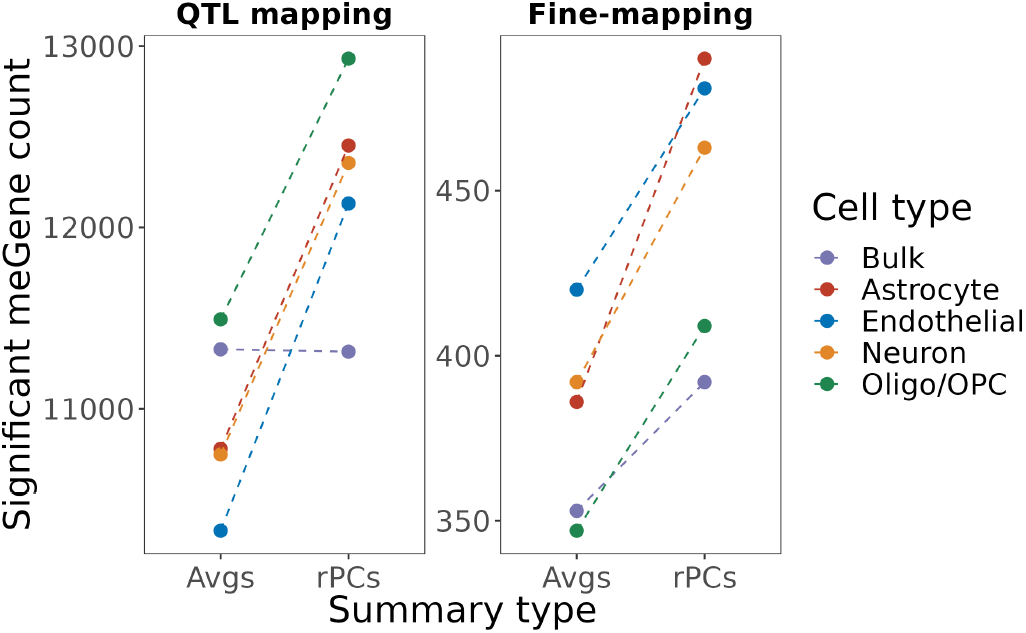
Enhanced gene identification in QTL mapping using rPCs. *regionalpcs* (rPCs) identified more genes with significant associations between methylation levels and genomic variants than averages in QTL mapping. The y-axis shows the number of genes identified as having significant associations with FDR *p*-values below 0.05 (QTL mapping) and genes further refined through fine-mapping with cluster PIP exceeding 0.5 (Fine-mapping). Color variations denote different cell types. Using rPCs consistently results in a higher identification rate for genes of interest across both QTL mapping and fine-mapping stages.

We used statistical fine-mapping of our meQTLs to identify credible sets of causal variants most likely to influence methylation. This analysis highlighted the widespread effects of genetic variation on DNA methylation, identifying both pan-cellular and cell type-specific meQTLs. After fine-mapping, a small fraction (4%) of meGenes harbored strong credible sets with a cluster posterior inclusion probability (CPIP) greater than 0.5 (Figure 5).

A comparison of fine-mapping between rPCs and averages revealed a greater number of meGenes associated with strong credible sets when using rPCs. We identified a median of 463 meGenes when summarizing with rPCs compared to 386 with averages across cell types. A limited overlap of meGenes (median of 95) was observed between the two methods, suggesting that each approach might capture distinct but complementary signals (Supplementary Figure 10). The median credible set size was very similar for rPCs (median n = 69 variants) and averages (median n = 64 variants), indicating that the choice of summary method for methylation does not alter the size of credible sets.

We found that the set of meGenes with strong credible sets was much more cell type-specific than the full set of meGenes identified with QTL mapping. Endothelial cells had the greatest number of unique meGenes with strong credible sets (rPCs 259, averages 189), while oligodendrocyte/OPCs had the fewest (rPCs 94, averages 43) (Supplementary Figure 9). We note that rPCs identified 7% more cell type-specific meGenes with strong credible sets compared to averages, suggesting that rPCs can identify more cell type-specific meQTLs.

### Integration of meQTLs and AD GWAS reveals causal relationships between DNA methylation and AD risk

We integrated our meQTLs with an AD GWAS (76) to identify putative causal associations between gene-level DNA methylation and AD risk. Our GWAS integration pipeline is comprised of three parts: colocalization with fastENLOC (46), instrumental variable analysis with probabilistic transcriptome-wide association study (PTWAS) (47), and the probabilistic integration of these two analyses with INTACT (48).

Using colocalization, we identified individual variants and loci likely to be causal for both a methylation QTL and the GWAS trait, AD. Across all cell types, about 1% of genes with strong meQTL credible sets were colocalized with AD at a gene-level colocalization probability (GLCP) threshold of greater than 0.5. A median of six genes with strong evidence of colocalization were identified using both rPCs and averages (Supplementary Table 8) for a total of seventeen colocalized genes identified across cell types. Four of these genes were found in the AD Open Targets database (AD Association Score greater than 88th percentile). One of these genes, *MS4A6A*, had strong evidence of colocalization in neurons and was found in the NIAGADS list of high-priority AD genes. *MS4A6A* is a four-transmembrane domain protein reported to be differentially expressed in the brains of AD subjects compared to controls (77, 78). Colocalized genes were predominantly cell type-specific, with the greatest number of genes identified in astrocytes and neurons for rPCs and averages, respectively (Supplementary Figure 11). The median credible set size of colocalized genes was 36% smaller for genes identified with rPCs (a decrease of 69 to 44 variants) and about 44% smaller for genes identified with averages (a reduction of 64 to 36) compared to the median credible set size for all other genes with strong credible sets (Supplementary Table 9). This decrease in credible set size may indicate that genes that colocalized with AD risk tend to have smaller meQTL credible sets than genes that are not colocalized. None of these colocalized genes were within one centimorgan (cM) of the *APOE* locus, indicating that they are independent signals from the effect of the *APOE*4 variant and other strong AD risk variants in *APOE*.

We used instrumental variable (IV) analysis with PTWAS to identify causal relationships between meQTLs and AD risk. Consistent with prior findings (79), we found that PTWAS identified a greater number of genes associated with AD (n = 43) compared to colocalization (n = 6) at a genome-wide significance threshold (*p <* 5 *×* 10^*−*8^, Supplementary Table 8). In contrast to the colocalized genes we identified, approximately 32% of the genes identified by PTWAS through rPCs (and 44% using averages) were located within 1 cM of *APOE*, which can likely be attributed partly to “LD hitchhiking” (80). LD hitchhiking can occur when some variants that are strong meQTLs for genes near *APOE* but are not risk variants for AD are in LD with AD risk variants near *APOE* that are not meQTLs for those nearby genes. These discrepancies in the gene sets identified by colocalization and IV analysis motivated our use of INTACT (48) to reconcile them.

We observed strongly divergent results from our colocalization and IV analysis, as anticipated from previous studies (48). Colocalization cannot infer causal relationships between methylation and AD risk but is resistant to LD hitchhiking, while IV analysis can infer causal relationships but is vulnerable to LD hitchhiking (81–83). We used INTACT to integrate gene-level colocalization probabilities from our colocalization analysis and gene-level causal associations identified by PTWAS. This method assumes that genes with joint evidence of both colocalization and a causal association are more likely to represent genuine biological relationships (79) (Supplementary Figure 12). We identified 20 unique genes using INTACT compared to 17 with colocalization and 95 with PTWAS across cell types. We also observed that results could depend on which part of a gene’s methylation was summarized. For example, the AD gene *TOMM40* (84–86), had an INTACT posterior probability of 0.831 when summarized over the promoter in oligodendrocytes/OPCs compared to zero when summarized over the full gene region (Supplementary Figure 13). This example illustrates the importance of a comprehensive analysis of different region types when studying the causal relationship between DNA methylation and disease etiology.

Across four cell types and bulk tissue, a median of eight genes were identified with putative causal relationships between methylation and AD using rPCs to summarize methylation compared to seven genes when using averages to summarize methylation (Supplementary Table 8). A total of 17 high-confidence genes (with posterior probabilities exceeding 0.6) were identified using the rPCs in one or more cell types, 12 of which were identified exclusively by rPCs (Table 1). Among these, *MS4A4A* and *PICALM* were identified by NIAGADS as high-priority AD-related genes. *PICALM* was only identified when using rPCs to summarize methylation, highlighting the limitation of relying on averages for summarization. The gene *RELB*, identified as a causal gene in oligodendrocyte/OPCs, was recently found to be an eGene within the same cell type in a cell type-specific expression QTL (eQTL) analysis of the ROSMAP cohort (87). These results suggest the possibility of a multi-omic network in which genetic variation affects gene expression of *RELB* through methylation, ultimately affecting the risk of AD.

**Table 1.**
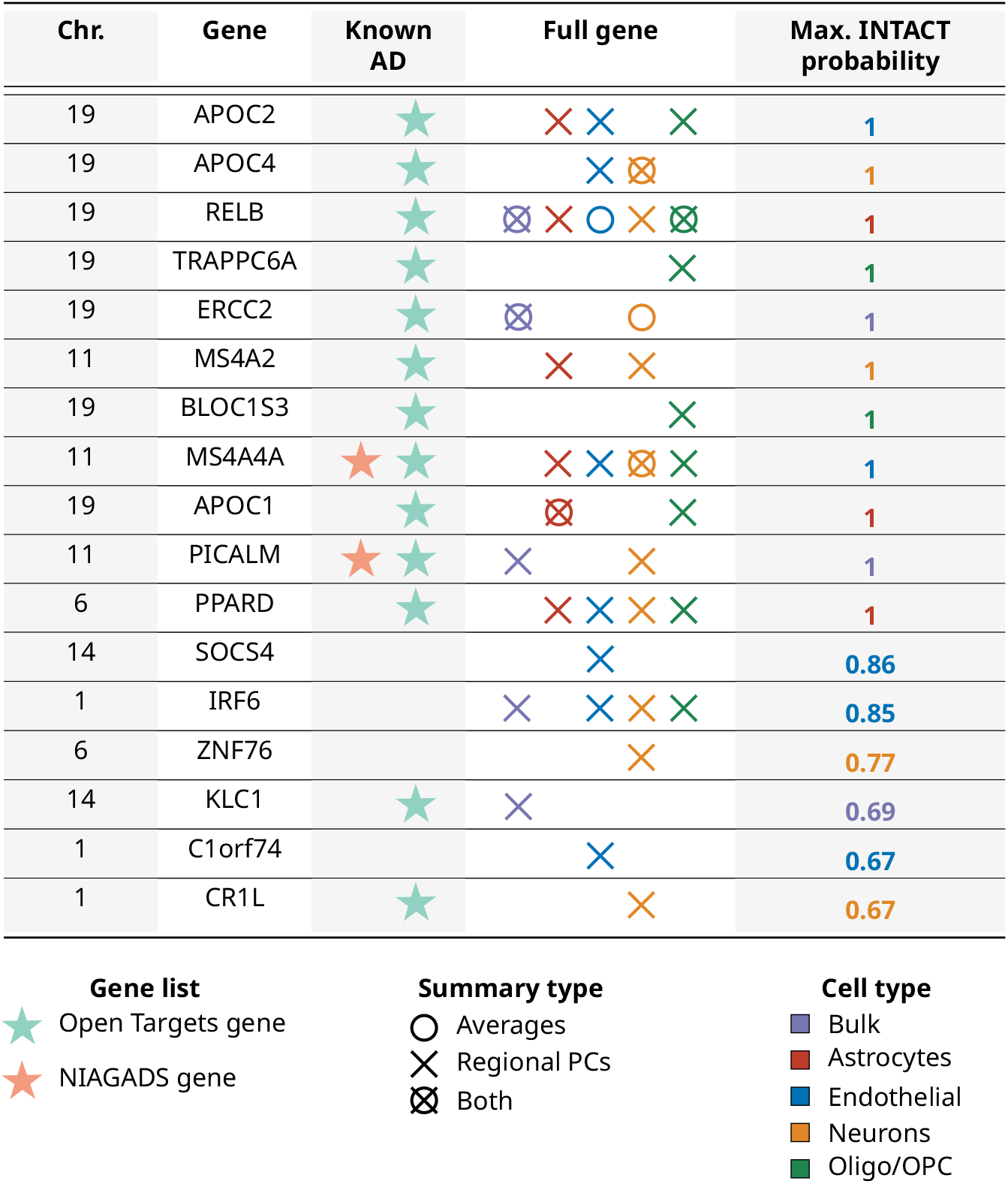
Candidate genes linking methylation and AD risk via INTACT analysis. Table highlighting candidate genes with shared genetic effects on methylation and Alzheimer’s disease susceptibility. These genes were identified with INTACT posterior probabilities exceeding 0.6 using rPCs, integrating evidence from colocalization and PTWAS analyses. Genes were identified using *regionalpcs* (denoted as ‘X’) and those identified by averages marked with ‘O’. Both are color-coded according to the cell type in which they were identified. Maximum INTACT probabilities for each gene, evaluated across cell types, are presented in the ‘Max. INTACT probability’ column and are also color-coded to the respective cell type. The chromosome on which each gene is specified in the far-left column, labeled ‘Chr.’.

Among the genes identified by INTACT, 33% (rPCs) and 25% (averages) were found within 1 cM of the *APOE* variant. Since this analysis incorporates colocalization, we anticipate that these genes are less likely to be false positives driven by LD hitchhiking than those in our IV analysis. These results support previous studies that implicated DNA methylation in the vicinity of *APOE* in AD pathogenesis (88, 89).

## Discussion

In this study, we introduced regional principal components (rPCs) as a novel approach for summarizing and interpreting gene-level methylation. rPCs provide an improved alternative to traditional CpG-level analysis and regional averaging. By focusing on methylation changes across various gene regions, we achieved a more complete understanding of methylation dynamics across the genome. The functional regions analyzed included promoters, 5kb upstream (of TSS) areas, gene bodies, and full genes. Our method’s effectiveness was further enhanced by employing cell type deconvolution, allowing us to summarize imputed cell type-specific methylation profiles derived from bulk tissue data. The accuracy of our deconvolution was supported by clustering our imputed cell type-specific profiles with nuclei-sorted data.

Leveraging the *RRBS-sim* simulation framework, our comparative analysis showcased the improvement of rPCs over traditional averaging in identifying differentially methylated regions. These simulations were designed to mimic reduced representation bisulfite sequencing (RRBS) data. Simulations varied in CpG site counts, sample sizes, proportions of differentially methylated sites, and mean methylation differences between cases and controls. The rPCs consistently outperformed averages in detecting differentially methylated regions, especially in scenarios with smaller sample sizes and fewer differentially methylated sites. This increased sensitivity of rPCs to subtle methylation differences was observed without yielding false positive results. These findings underscore the robustness of rPCs across a diverse range of study designs and types of methylation data.

We applied rPCs to the cell type-specific methylation data from the ROSMAP cohort to demonstrate its effectiveness in identifying AD-relevant methylation changes. We observed a stronger correlation between the degree of differential methylation detected with established gene relevance to AD pathology using rPCs compared to averages. The gene-level associations we discovered were predominantly cell type-specific, with astrocytes showing the most substantial methylation signal associated with CERAD score. Although some disease-relevant genes emerged from bulk methylation data, cell type-specific analyses yielded far richer sets of genes. We found relatively few associations of methylation changes with ADD diagnosis and *APOE* genotype compared to neuritic plaque burden and Braak stage. These results support previous findings (65) that endophenotypes derived from proteinopathy measures are more consistently associated with molecular traits than clinical diagnoses and individual genetic markers. Mapping meQTLs with rPCs further identified more genes with significant associations between DNA methylation and genetic variation than traditional averaging methods. Integration with AD GWAS through colocalization, instrumental variable analysis, and INTACT revealed 17 unique genes potentially mediating AD via DNA methy-lation, 12 of which were exclusively identified by rPCs. The unique genes included *MS4A4A* and *PICALM*, both identified by NIAGADS as high-priority AD-related genes.

While the rPCs method improves upon existing approaches for aggregating methylation by several metrics, interpreting the contribution of individual features remains challenging. Future extensions of this method could leverage approaches such as non-negative matrix factorization, factor analysis, or independent components analysis to generate more interpretable summaries. However, unlike PCA, they do not guarantee orthogonality between factors and there are no analytic approaches to choosing the number of components. We chose to use singular value decomposition (SVD) to estimate our eigenvectors, which can be viewed as a frequentist approach to PCA. SVD does not give us a full generative model that would allow us to quantify uncertainty about the latent space representation of regional methylation, but SVD is more computationally tractable for the larger number of regions we studied. Nevertheless, broadening our framework to include frequentist and probabilistic implementations of dimensionality reduction methods is a promising future direction for our method.

Our initial focus on methylation microarrays sets the stage for expanding rPCs to whole-genome methylation sequencing (WGMS) and other epigenomic datasets. The present scarcity of large-scale WGMS datasets limits comprehensive analyses, but as more extensive datasets emerge, we expect our methodology to significantly enhance our understanding of the role of methylation in complex diseases. Beyond WGMS, we foresee the application of rPCs to a diverse array of epigenomic data, including ATAC-seq, ChIP-seq, and HiC techniques. This expansion will enable a holistic view of the epigenetic landscape and foster the development of models capable of integrating insights across various epigenomic modalities.

The application of our *regionalpcs* methodology to brain data highlights a notable challenge: the absence of detectable microglial signals. This challenge is likely due to the rel-atively low abundance of microglia in the dorsolateral pre-frontal cortex. Despite this limitation, this approach has demonstrated promise in mapping expression QTLs in microglial populations (90), suggesting future research paths. Employing techniques such as nuclei sorting or intact cell isolation is essential for acquiring sufficient samples for methylation and other functional genomic studies, particularly for cell types like microglia. Subsequent validation of our findings using these refined cellular populations will be critical in corroborating the genes and pathways we have implicated in AD. Such validation is the first step in the development of novel therapeutic strategies and enhancing our capacity to predict AD risk, thereby advancing the field’s move from broad observations to targeted interventions.

In conclusion, the rPCs method is a significant advancement in the analysis of methylation data, particularly for complex diseases like AD. Our findings underscore the value of rPCs for uncovering the complex mechanisms underlying disease, providing novel research directions that could lead to targeted therapeutic interventions and refined disease risk assessments. Further, the versatility of rPCs extends its potential impact beyond AD to encompass a broad spectrum of conditions where methylation plays a pivotal role. With the *regionalpcs* Bioconductor package, we offer the scientific community an accessible tool to advance epigenetic research, fostering discoveries that could transform our approach to disease understanding and treatment.

## Methods

### A. Regional principal components

Let 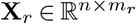 be a matrix of *n* samples and *m*_*r*_ CpG sites for distinct regions *r* ∈ *{*1, 2, …, *R}*, where each entry of the matrix corresponds to a methylation quantification value such as a beta value or an M-value. Note that in contrast to Figure 1, in this section we will work with **X**_*r*_ instead of **X** . Without loss of generality, we assume that **X**_*r*_ has been transformed by the inverse normal transformation

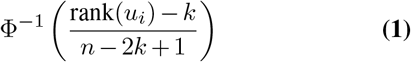

where Φ^*−*1^ is the probit function, and *k* ∈ (0, 1*/*2) is an adjustable offset. The inverse normal transformation guarantees that each column is normally distributed when there are no ties in the data. To compute the regional principal components (rPCs) of region *r*, we first perform singular value decomposition on **X**_*r*_ to decompose the matrix into,

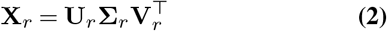

where **U**_*r*_ is an *n × n* orthogonal matrix whose columns are the left singular vectors of **X**_*r*_, **Σ**_*r*_ is an *n × m*_*r*_ diagonal matrix containing singular values of **X**_*r*_ in descending order, and **V**_*r*_ is an *m*_*r*_ *× m*_*r*_ orthogonal matrix whose columns are the right singular vectors of **X**_*r*_. The eigenvalues of the co-variance matrix 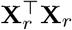 are the squares of the singular values along the diagonal of **Σ**_*r*_.

We use the Gavish-Donoho method (49) to estimate the optimal number *k*^*∗*^ of principal components needed to capture the essential variability in the data. The Gavish-Donoho method estimates the optimal number of principal components purely as a function of the ratio of number of columns to the number of rows of the matrix as well as estimated noise in the matrix. We select the first *k*^*∗*^ right singular vectors (columns of **V**_*r*_) and denote the resulting matrix as

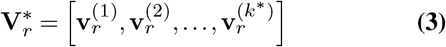

where 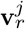 is the *j*-th column of **V**_*r*_ . The columns of 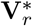 are the first *k*^∗^ principal components of **X**_*r*_. Finally, we project **X**_*r*_ onto the principal components 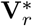 to compute the princi-pal component scores,

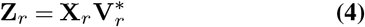

where **Z**_*r*_ is an *n × k*^∗^ matrix, representing the transformed coordinates of the original data **X**_*r*_ in the reduced-dimensional space spanned by the first *k*^∗^ principal components contained in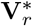. Each row corresponds to a sample in **X**_*r*_. We refer to each column of **Z**_*r*_ as the regional PCs (rPCs), which are used to summarize regional methylation for all downstream comparative analyses.

### B. Simulation Analysis

We employed the *RRBS-sim* tool, developed by ENCODE (51), to simulate reduced representation bisulfite sequencing data. This tool models CpG site locations using a hidden Markov model, sequencing coverage through a two-component gamma distribution mixture, and spatial correlations between sites via a Gaussian variogram.

Our simulations focused on methylation within gene regions. We explored four key parameters: the count of CpG sites, the sample size, the proportion of differentially methylated (DM) CpG sites, and the methylation percentage difference between case and control groups at DM sites. For CpG site counts, we referenced an ENCODE RRBS dataset from a K562 cell line, determining the typical number of CpGs within full gene (mean = 50, median = 29) and promoter regions (mean = 19, median = 12) using the *TxDb*.*Hsapiens*.*UCSC*.*hg19*.*knownGene* Bioconductor package for gene annotations (91). Based on these findings, we conducted simulations with 20 and 50 CpG sites. Sample sizes of 50, 500, and 5,000 were chosen to represent both small and large datasets. We tested DM site proportions at 0.25, 0.50, and 0.75, and differential methylation percentage differences from 1 to 9% in 1% increments. For differentially methylated regions, we set the length at half the number of CpG sites, ensuring a minimum of 2% CpG density.

In total, 162 unique parameter combinations were used to simulate methylation beta values for 1,000 regions per set. Samples were equally divided into cases and controls. Methylation data for each CpG site underwent normalization using the `RankNorm` function from the *RNOmni* R package (92). Methylation was summarized for each region using both averages and regional principal components.

For differential methylation analysis, we utilized the `lmFit` function from the *limma* package (93), fitting linear models to the summarized methylation data of each gene region. Stability in test statistics was achieved using the empirical Bayes approach via *limma*’s `ebayes` function, and *p*-values were Bonferroni-adjusted for multiple testing. Regions with adjusted *p*-values below 0.05 were classified as significantly differentially methylated.

To assess the accuracy in identifying true negative results, we generated a matrix comprising regions with varied differential methylation levels. This included 700 regions with no methylation difference (true negatives) and 300 regions with methylation differences varying from 1 *×* 10^*−*3^ to 9% at intervals of 1% (30 each). Methylation was summarized using both averages and rPCs, followed by differential methylation analysis as previously described.

### C. Study cohort

To explore the relationship between DNA methylation and AD, we analyzed a subset of the ROSMAP cohort with DNA methylation and whole-genome sequencing (WGS) data. DNA methylation and WGS data were obtained from the Synapse AD Knowledge Portal following the submission of a Data Use Agreement (https://adknowledgeportal.org). We applied quality control filtering to the data (detailed below). Specifically, we executed a series of filtering steps on both the DNA methylation and WGS data, opting for individuals who possessed data for both and adhered to our filtering standards. This filtering procedure resulted in a final cohort of 563 individuals, all of whom self-identified as white. The average age of the cohort was 86.4 years, with females comprising 63% of the cohort (N = 356, Supplementary Table 2). Each individual in this cohort provided a single sample for methylation and genotype analysis, forming the basis for all subsequent investigations in this study.

### D. Preprocessing genotype data

Subject genotypes were obtained from ROSMAP WGS VCF files (Synapse ID: syn11707420) generated by The Whole Genome Sequence Harmonization Study, a multi-institutional effort to harmonize genotype data from several AD cohorts (94). WGS libraries were constructed using the KAPA Hyper Library Preparation Kit and sequenced on an Illumina HiSeq X sequencer. Raw sequence reads were aligned to the GRCh37 human reference genome using the Burrows-Wheeler Aligner (BWA-MEM v0.7.08). The aligned data were processed following the GATK best practices workflow, which includes haplotype calling, joint genotyping with GATK v3.5, and variant filtering with Variant Quality Score Recalibration (VQSR).

#### D.1. Variant filtering

Variants were normalized with *BCFtools* (95) and lifted over from GRCh37 to GRCh38 using *LiftoverVcf* from GATK v4.0.10.0 (96). We used PLINK2 (97) to remove multiallelic variants, variants with more than 5% missing genotype calls, variants with minor allele frequency less than 0.01, variants that failed the haplotype-based test for non-random missing genotype data with a *p*-value threshold of 1 *×* 10^*−*9^, variants that deviated from Hardy-Weinberg equilibrium at *p <* 0.001, variants on the sex chromosomes, and variants that are not at CpG sites. Our filtering pipeline yielded 9,470,439 high-quality autosomal variants for subsequent analyses.

#### D.2. Sample filtering

Utilizing PLINK (98), we excluded samples exhibiting missing call rates exceeding 5%. We also used the *SmartPCA* algorithm from EIGENSTRAT (99) to detect population outliers using the default parameters of 10 principal components and six standard deviations; none were discovered in our cohort.

### E. Selection and interpretation of ROSMAP phenotypic traits

We chose four phenotypic traits from the ROSMAP cohort to test for associations with DNA methylation. These traits were a CERAD score-based diagnosis of ADD, Braak staging for neurofibrillary degeneration, clinical diagnosis of ADD, and *APOE* genotype (Supplementary Table 2).

The CERAD score (55), a tool from the Consortium to Establish a Registry for Alzheimer’s Disease, offers a semi-quantitative assessment of neuritic plaque density in neocortical regions from autopsy brain samples. Based on the CERAD score, a score between one through four was assigned to each individual. This score, treated as a continuous value in our study, has integer values ranging from one (indicating definite AD) to four (indicating no AD). The distribution of subjects across CERAD score categories is Definite AD (N = 170), Probable AD (N = 195), Possible AD (N = 59), and No AD (N = 139) (Supplementary Table 2).

Braak staging (54) evaluates the anatomical extent of neurofibrillary degeneration, with integer scores from zero to six reflecting the stereotypical distribution of neurofibrillary tangles across limbic and cerebral cortical regions. Since our focus was on methylation data from the dorsolateral prefrontal cortex (DLPFC), we discretized the Braak stage into a categorical variable indicating the presence (≥ 5, N = 132) or absence (*<* 5, N = 431) of neurofibrillary tangles involving a neocortical region, which includes the DLPFC.

To create a comprehensive diagnosis variable, we combined clinical and pathology scores. Our clinical measure was the final consensus cognitive diagnosis variable (cogdx), determined by a dementia-specialist neurologist based on all clinical data available, excluding post-mortem information. Our pathology scores were the previously described CERAD score and Braak stage. We labeled subjects as “AD” if they were diagnosed with cognitive impairment or dementia (cogdx 2-5), exhibited a high Braak stage (V or VI), implying the spread of neurofibrillary tangles to neocortical regions, and had a low CERAD score (≤ 2) indicating probable or definite AD based on neuritic plaques (N = 217). Subjects were designated as “Control” if they had a clinical diagnosis of no cognitive impairment (cogdx = 1), a low Braak stage (≤ 3), and a high CERAD score (≥ 3) (N = 80). In analyses of our diagnosis variable, we omitted individuals who did not fit into either the “AD” or “Control” categories as we found this indicative of a discrepancy between the clinical and pathological variables (N = 266).

In our analyses of the *APOE* genotype, we focused on individuals with *ϵ*3*/ϵ*3 (N = 340) and *ϵ*3*/ϵ*4 (N = 129) genotypes so that we could focus on exploring the impact of a single *ϵ*4 allele, which is associated with an increased risk of AD. The *ϵ*3 allele is the most common and is used as the reference in estimating relative risk, while the rare *ϵ*2 allele is reported to be protective against AD (84, 100). We excluded individuals carrying the *ϵ*2 allele from our *APOE* analyses because we did not have a sufficient sample to robustly analyze these subjects as a distinct group.

### F. Preprocessing methylation data

IDAT files of DNA methylation microarray from ROSMAP (Synapse ID: syn7357283) were generated as described in De Jager *et al*. (7). DNA was extracted from frozen dorsolateral prefrontal cortex sections of deceased ROSMAP participants. The Illumina Infinium HumanMethylation450 bead chip assay was used to quantify DNA methylation.

We used the *minfi* R package (101) for normal-exponential convolution using out-of-band probes dye bias correction, mapping of probes to the genome, and removal of probes in sex chromosomes or probes overlapping any genomic variant in Illumina and dbSNP references (102). The *ewastools* package (103) was used to compute detection *p*-values, and we used the *wateRmelon* package (104) to remove probes with detection *p*-values*>* 0.01 or bead count *<* 3. We used the Beta Mixture Quantile dilation (BMIQ) normalization method (105) to correct for biases resulting from differences between type 1 and type 2 probes in the Infinium Human-Methylation450k platform. Lastly, we removed probes overlapping any genomic variant called in any subject from the WGS genotype data from the ROSMAP cohort. After all filtering steps, 365,899 probes were used for downstream analysis.

Samples were excluded if their bisulfite conversion efficiency was below 80% or if they failed the *ewastools* quality control metrics estimated following Illumina’s BeadArray Controls Reporter Software Guide. In total, 87 samples were removed because they did not pass these quality control measures.

### G. Cell type deconvolution

We used cell type deconvolution to estimate cell type-specific methylation from the processed and filtered bulk tissue methylation data. We estimated cell type-specific methylation from 365,899 probes and 563 samples for which matched whole-genome sequencing data was available. Our deconvolution pipeline has two steps: estimation of cell type proportions and imputation of cell type-specific methylation profiles. We used the *EpiSCORE* R package (62) to estimate the cell type proportions for six major brain cell types: astrocytes, endothelial cells, neurons, oligodendrocytes, oligodendrocyte progenitor cells, and microglia, using the brain reference panel provided by *EpiSCORE*. As detailed in (62), this reference imputed cell type-specific methylation at specific CpGs from a single-cell RNA-seq data set.

We estimated cell type proportions for each sample using the `wRPC` function from *EpiSCORE*. Our estimated cell type proportion for microglia across all samples was zero, likely due to their low proportions in the DLPFC, so we had to exclude microglia from further analysis. Additionally, the proportions of oligodendrocytes were very low (mean = 0.003%, median = 0%), with only four individuals having a non-zero estimated proportion. This is likely because of the high similarity to oligodendrocyte progenitor cells, for which we estimated proportions consistent with the expected abundance of oligodendrocytes. Under the assumption that it was difficult for *EpiSCORE* to distinguish between these very similar cell types, we combined oligodendrocytes and oligodendrocyte progenitors into a single proportion. This resulted in cell type proportion estimates for four cell types: astrocytes, endothelial cells, neurons, and oligodendrocyte/oligodendrocyte progenitor cells.

To validate the estimated neuron proportions, we used an independent brain reference panel and estimation method published by Houseman *et al*. (66) for estimating cell type proportions from the `estimateCellCounts` function in the *minfi* package. This reference panel only distinguishes neurons from non-neurons but was generated from methylation microarray data from sorted nuclei from brain samples. We found that the neuron proportions estimated by the *EpiSCORE* and *Houseman* methods were highly correlated (Pearson *r* = 0.8, *p <* 0.001). We used the cell type proportion estimated by *EpiSCORE* for downstream analysis because these proportions were more granular for non-neuronal cell types.

We used the *Tensor Composition Analysis (TCA)* (63) package for the imputation of cell type-specific methylation profiles. *TCA* uses a multivariate regression model that decomposes a matrix of bulk tissue methylation beta values into a tensor of cell type-specific beta values given a vector of cell type proportions for each sample. Our inputs to *TCA* were the filtered bulk tissue methylation data and the cell type proportions estimated by *EpiSCORE*.

*TCA* accommodates the inclusion of several types of covariates in the deconvolution model. *TCA* accounts for variables that may affect methylation at the cell-type level (C1) and global level (C2). We tested models with age and sex as C1 covariates, batch, post-mortem interval, and study as C2 covariates, or a combination of C1 and C2 covariates. The model containing only C2 covariates emerged as the best fit since the C1 covariates seemed to exaggerate the effects of age and sex in downstream differential methylation analysis. We used other options for *TCA* to constrain the model’s mean parameter during optimization and refit the estimated cell type proportions iteratively. The resulting refitted cell type proportions were used as covariates to model cell type proportions in downstream analyses. The final output of *TCA* was imputed CpG-level methylation beta values in all subjects for the four cell types whose proportions we estimated with *EpiSCORE*.

### H. Normalization and cleaning of deconvolved methylation data

We removed probes with zero variance for each cell type-specific dataset and the original bulk tissue dataset. We applied the rank-based inverse normal transformation (INT) using the `RankNorm` function from the *RNOmni* package (92). Transforming methylation beta values to M-values or inverse-normal transformed values is a standard approach to reduce heteroscedasticity and give the methylation values an approximately Gaussian distribution for downstream statistical analyses (106). We compared the non-parametric inverse normal transformation to the logit transformation for M-values and found similar results, consistent with previous reports comparing these approaches (107). We chose to use inverse normal transformation because it does not produce large, transformed values for beta values close to 0 or 1, as seen with M-values.

We identified technical variables strongly correlated with our methylation data by estimating the top methylome-wide principal components (PCs) from the full dataset. The number of PCs was chosen using the Gavish-Donoho method (49), described in more detail in the “*Summarizing regional DNA methylation*” methods section. We correlated traits with PCs using Spearman’s rank correlation for numeric variables and the Kruskal-Wallis test for categorical variables. We used a Bonferroni-adjusted *p*-value *<* 0.05 as our significance cutoff. We found that batch, post-mortem interval (PMI), study (ROS or MAP), sex, age at death, and cell type proportions were highly correlated with one or more PCs in datasets (Supplementary Figure 14). We used the `removeBatchEffect` function from the *limma* package (93) to remove the effects of the top four methylome-wide PCs and all of the previously described covariates except for sex, which we instead included as a covariate in downstream analyses. After removing these covariates, some correlations persisted between the technical variables and methylome-wide PCs, and we incorporated them as covariates in all downstream analyses (Supplementary Figure 14).

### I. Visualizing deconvolved cell type methylation profiles using UMAP

The cell type-specific and bulk tissue methylation profiles were visualized with the Uniform Manifold Approximation and Projection (UMAP) method for dimensionality reduction (67) as implemented in the *uwot* package (108). We transformed the deconvolved methylation data into M-values and used principal components analysis on the union of all cell type-specific datasets and the bulk tissue dataset using the default parameters of the `pca` function from the *PCAtools* package (109). To determine the number of principal components to retain, we used the Gavish-Donoho method (49), described in the “Summarizing regional DNA methylation” section. The UMAP projection was estimated with the `umap` function from the *uwot* package, setting the number of neighbors to 500, the spread to 10, and the minimum distance to 5.

### J. Annotating genomic region types

Our investigation focused on DNA methylation patterns across four genomic regions: promoters (spanning from 1kb upstream of the transcription start site (TSS) to the TSS itself), gene bodies (ranging from the TSS to the downstream end of the 3’ untranslated region (UTR)), 5kb upstream region (extending from 5kb upstream of the TSS to the TSS), and the full gene (a union of the 5kb upstream and gene body regions). We delineated these regions using data from the *Annotatr* package (110), Gencode (111), and the UCSC Table Browser (112), all of which used the GRCh38 reference genome build.

We used the Gencode v32 gene annotations for *Homo sapiens* to select genes and transcript isoforms. We filtered this annotation for protein-coding transcripts with a transcript support level less than four. For genes whose canonical isoform passed these filters, we used this isoform, and for genes with no canonical isoform after filtering, we used the longest available isoform instead. We chose the four genomic region types described above because these regions are generally consistent across transcript isoforms, in contrast to more variable features such as exons and introns.

### K. Summarizing regional DNA methylation

We designed an approach to summarize gene-level DNA methylation within the four gene regions with two different methods: averaging and *regionalpcs* (rPCs). We extracted the genomic positions of each probe from the GRCh37 annotations of the HumanMethylation450K array provided by *RnBeads* package (34). We remapped the genomic positions to GRCh38 using the *GenomicRanges* and *liftOver* packages (113, 114), and CpGs were assigned to region types using the `findOverlaps` function from the *GenomicRanges* package.

We summarized methylation with averaging by calculating the mean across all CpGs falling within a region type. We summarized methylation with rPCs by estimating the principal components of CpGs within a region using the `pca` function from the *PCAtools* package. We used the Gavish-Donoho method implemented by the `chooseGavishDonoho` function in *PCAtools* to select the optimal number of principal components to represent methylation in a genomic region. The Gavish-Donoho method (49) estimates the optimal number of principal components purely as a function of the ratio of the number of columns to the number of rows of the matrix as well as estimated noise in the matrix. Because our matrices are always scaled and centered, the noise parameter always had a value very close to 1. We assessed both the Gavish-Donoho and Marchenko-Pasteur methods (50) for selecting the number of principal components and chose the Gavish-Donoho method because it was more conservative.

We applied the averaging and rPCs methods for each gene and region type which included at least one CpG. In cases where a region type for a gene contained only one CpG, we used the original CpG methylation value to represent that gene in summarized datasets for averages and rPCs. In total, we summarized 20 datasets using averages and rPCs, encompassing all four region types and four cell types as well as bulk tissue methylation.

### L. Assessment of rPCs for potential global methylation factors

We performed a correlation analysis between rPCs across multiple genes to test for the possibility that the rPCs may capture global methylation effects rather than regional variation. We estimated the correlation between the first three rPCs (ordered by the proportion of variance explained) for each gene with those of genes located on different chromosomes. Our underlying hypothesis posited that if rPC1 primarily captured global methylation effects, it would exhibit higher inter-gene correlations compared to other rPCs.

We estimated partial correlations that adjusted for several confounding variables: sex, age, PMI, study, batch, and cell type proportions. This analysis revealed marked disparities in the distribution of correlations, with rPC1 displaying the highest inter-gene correlations (Supplementary Figure 3a).

We next tested if incorporating the top global methylation PCs into our partial correlation model would reduce the inter-gene correlations we observed. The number of global PCs was determined using the Gavish-Donoho method. The inclusion of these global PCs attenuated the previously observed correlation discrepancies among the rPCs, demonstrating that these rPCs were capturing global signals that were also reflected in the top global PCs (Supplementary Figure 3b).

We sequentially excluded each global PC from the model, re-estimated partial correlations for all other variables, and determined that the first four global PCs drove the shared signal. When any of these global PCs were omitted, intergene correlation distributions were discordant between rPCs (Supplementary Figure 3c).

We refined our methylation summarization model by removing the first four global PCs with the `removeBatchEffect` function from the *limma* package prior to estimating the final set of rPCs. The adjustment was validated with a partial correlation analysis, which found negligible disparities in inter-gene correlations among the newly derived rPCs (Supplementary Figure 4d). These results suggest that the rPCs estimated after removing the top global PCs were no longer influenced by global methylation patterns and instead captured regional methylation signals.

### M. Differential methylation analysis across cell and region types

We performed a comprehensive differential methylation analysis of the processed methylation data for each region type and cell type, using averages and rPCs to summarize methylation. Our disease phenotypes described in the “*Selection and interpretation of ROSMAP phenotypic traits*” methods section were CERAD score (a measure of neuritic plaques), Braak stage (a measure of Tau protein), Alzheimer’s disease diagnosis, and *APOE* genotype.

In addition to the previously described disease phenotypes, our model included age of death, post-mortem interval (PMI), study (ROS or MAP), batch, sex, and cell type proportions we estimated for all four cell types. We also included the top global methylation principal components (PCs) computed from the full CpG-level methylation data. These global PCs were calculated separately for each of the four cell type datasets across all CpGs and samples, and the number of PCs was determined using the Gavish-Donoho method described in the “*Summarizing regional DNA methylation*” methods section. To prevent the removal of disease-associated signals from the methylation data, we omitted global PCs if they were significantly correlated with the outcome trait and not with batch, PMI, and study (Bonferroni-adjusted *p <* 0.1). We used Spearman rank correlation for numeric variables and the Kruskal-Wallis test for categorical variables.

We identified differentially methylated genes with the `lmFit` function from the *limma* package by fitting linear models to the methylation data for each summarized gene region. The methylation was normalized using rank-based inverse normal transformation (INT) from the `RankNorm` function in the *RNOmni* package. We used the empirical Bayes method implemented by the `ebayes` function in the *limma* package to obtain more stable estimates of the test statistics, and we used the Benjamini-Hochberg method for multiple test adjustments.

### N. Identifying high-confidence Alzheimer’s disease genes

We used the Open Targets database (74) to identify a high-confidence set of genes associated with Alzheimer’s disease. This comprehensive resource compiles information on the relationship between genes and diseases from a range of sources. We used a curated list of 6,595 genes linked to Alzheimer’s disease based on seven distinct evidence categories: genetic associations, somatic mutations, drugs, pathways systems biology, text mining, RNA expression, and animal models. Open Targets assigns an overall association score to each gene on the list according to the available evidence.

Mapping cis-QTLs We used tensorQTL (115) to map cis-methylation QTLs (meQTLs) for each gene in every region type and cell type. tensorQTL is a reimplementation of FastQTL (116) that uses linear models to identify correlations between alternate allele count and molecular traits such as methylation. For every gene, we tested for associations between alternate allele count and gene-level methylation summaries within a cis-window of 1 million base pairs upstream and downstream of the transcription start site (TSS) for both averages and rPCs.

We incorporated several covariates into the QTL mapping model to control for potential confounding factors, including age of death, post-mortem interval (PMI), batch, study (ROS or MAP), sex, and cell type proportions for our four cell types of interest. We also included the top 10 genotype principal components (PCs) and the top methylation PCs, determined using the Gavish-Donoho method (49) as previously described. We used the `get_significant_pairs` function from tensorQTL with default parameters to extract significant meQTLs with a false discovery rate (FDR) threshold of 0.05 that accounted for the number of genes and variants tested. We used the beta-approximated *p*-value distribution estimated by tensorQTL as a faster approximation of permutation testing.

### O. Fine-mapping QTLs

We used fine-mapping to identify credible sets of causal variants in our meQTLs for gene region-level methylation summaries estimated with averages and rPCs. First, we used TORUS (117) to estimate priors for each variant based on their distance to the transcription start site. Next, we used DAP-G (118) to identify credible sets from the priors generated by TORUS and the individual-level genotype and methylation data we also used for meQTL mapping. DAP-G uses a Bayesian framework to estimate the posterior probabilities of each genetic variant associating with a molecular trait such as methylation. We chose an R2 cutoff of 0.25 as the minimal LD correlation needed for two variants to be included within the same credible set. Credible sets represent a group of genetic variants that contain a causal association with a trait independent of any other credible set, and a single gene can contain multiple credible sets. Because the variants within the credible set are in LD with each other, the exact causal variant cannot be determined solely from QTL mapping unless the credible set contains a single variant. Our threshold for defining a strong credible set in downstream analyses was a posterior inclusion probability (PIP) of 0.5 or greater.

### P. Fine-mapping GWAS

In order to integrate our meQTLs with genome-wide association studies (GWAS) of AD risk, we first needed to fine-map the GWAS summary statistics. We used DAP-G (118) for this task similar to how we used it for fine-mapping the QTL data.

We used a published meta-analysis of Alzheimer’s Disease risk GWAS (76). We converted the genomic positions in the GWAS summary statistics from GRCh37 to GRCh38 using the GATK v4.0.10.0 LiftoverVcf tool to match genomic coordinates in our meQTL analysis. We matched 8,114,981 variants in the GWAS summary statistics to the variants tested in our meQTL analysis and removed 6,912 variants with inconsistent reference/alternate alleles.

Since we only had summary statistics rather than individual-level data, we used linkage disequilibrium (LD) blocks for fine-mapping. We used a published set of 1,361 LD block annotations in GRCh38 coordinates (119) and intersected them with the GWAS variants in the summary statistics using the *GenomicRanges* package.

We used PLINK 1.90b6.21 to create LD matrices within each LD block and ran DAP-G with default parameters. Similar to our QTL fine mapping, we identified credible sets across several LD blocks representing independent AD risk signals.

### Q. GWAS integration

We integrated our meQTLs with the Wightman *et al*. AD GWAS (76) to identify putative causal associations between gene-level DNA methylation and AD risk. In this framework, we can view the perturbation of DNA methylation by germline genetic variants as quantified by meQTLs as a “natural experiment” that tests how changes in DNA methylation affect the risk of developing AD. Our GWAS integration pipeline is comprised of three parts: colocalization with fastENLOC (46), instrumental variable analysis with probabilistic transcriptome-wide association study (PTWAS) (47), and the probabilistic integration of these two analyses with INTACT (48).

#### Q.1. Colocalization

We used colocalization with fastENLOC (46) to identify genes with evidence of shared causal variants between meQTLs and AD risk. fastENLOC uses a Bayesian framework to estimate the posterior probability of colocalization between a QTL and a GWAS trait while also estimating the overall GWAS heritability enrichment of QTL variants across all genes tested for QTLs.

The inputs for this analysis were the fine-mapped QTL and AD GWAS summary statistics generated in the previous steps. We limited our colocalization analysis to fine-mapped SNPs assigned to credible sets in the meQTL or GWAS summary statistics with posterior inclusion probabilities (PIPs) < 1 *×* 10^*−*4^. We used the default parameters for fastENLOC and set the “total variants” flag to 12,688,339 variants.

We used a gene-level colocalization probability (GLCP) threshold of 0.50 to decide which genes were colocalized with AD risk. The GLCP is a locus-level measure of colocalization which addresses some of the limitations found in variant-level colocalization analysis, such as the potential for excessive type 2 errors. It quantifies the probability that a particular genomic locus harbors both a molecular QTL with a causal effect and a causal GWAS variant. This approach has more power to detect colocalization between methylation QTLs and AD, improving our understanding of the molecular basis of Alzheimer’s disease.

#### Q.2. Instrumental variable analysis with PTWAS

We used PTWAS (47) to explore potential causal relationships between DNA methylation and AD risk. The premise of IV analysis is that genetic variants identified as risk variants for AD in a GWAS mediate their effects through changes in DNA methylation. PTWAS constructs a composite instrumental variable by integrating weights from multiple independent methylation QTLs for each gene. The weights are derived from their respective posterior effect sizes from our fine-mapping meQTL analysis. A causal association Z-score is computed as a weighted sum of GWAS trait Z-scores for each independent instrument in the tested gene.

We followed the published PTWAS pipeline which first calculates weights for each gene with the `ptwas_builder` function from DAP-G using the individual level data previously used to map meQTLs and results generated via DAP-G. Effect sizes and standard errors for the Wightman *et al*. GWAS were computed based on the reported Z-scores for individual genetic variants. We used GAMBIT (120) to perform the PTWAS scan step with the 1000 Genomes project, phase 3 version 5 LD reference panel for Europeans (121), chosen for its phase and ancestry match to the ROSMAP cohort. The LD panel was converted from GRCh37 to GRCh38 genome build using the *liftOver* tool, and any mismatches between reference and alternative alleles were rectified using PLINK v1.90b6.21. The significance of putative causal genes was defined by the Z-scores generated in the PTWAS scan. We set a genome-wide significance threshold considering genes with an absolute Z-score greater than 5.45 (corresponding to a *p*-value *<* 5 *×* 10^*−*8^) as having a likely causal association with AD.

#### Q.3. Integration of colocalization and TWAS with INTACT

We used the Integration of TWAS and Colocalization (INTACT) approach to integrate the results from our colocalization and PTWAS analyses (47). Colocalization is designed to identify QTLs that share causal variants with a GWAS trait. In contrast, TWAS is designed to uncover molecular features such as methylation mediating causal variants’ effects on AD risk. INTACT is designed to reconcile the discrepancies observed when colocalization and TWAS are used separately by multiplying the posterior probabilities from colocalization and gene-trait associations Z-scores from TWAS. This integration provides a single metric to quantify the probability of a causal association between genetic variations, methylation changes and Alzheimer’s disease traits. As inputs to INTACT, we used the gene-level colocalization probability (GLCP) estimated by fastENLOC and the gene-level Z-score statistics estimated by PTWAS. We used the default settings for INTACT with a linear prior function with a truncation threshold of 0.05 and we filtered the posterior probabilities at a threshold of 0.6. This filtered set of genes was used as our final list with strong evidence of a causal relationship between DNA methylation and AD risk.

## Data Availability

The original whole genome sequencing data from ROSMAP (Synapse ID: syn11707420) and the IDAT files for DNA methylation microarray (Synapse ID: syn7357283) are accessible for download from the Synapse AD Knowledge Portal (https://adknowledgeportal.org) subject to a valid data use agreement. Additionally, processed individual-level data, which include deconvolved cell type-specific methylation profiles (Synapse ID: XXX), and gene region methylation summary data (Synapse ID: XXX) are available for download from Synapse upon agreement to data use terms.

For broader accessibility, we have also made available summarized results that do not contain individual-level data; these can be downloaded from Synapse without the requirement for a data use agreement (Synapse ID: syn57397239). These summarized datasets include differential methylation results (syn57397243), tensorQTL results (syn57398471), DAP-G fine-mapped QTL results (syn57398365), variant-level colocalization results (syn57397535), and PTWAS results (syn57398295). Gene-level scores from GWAS integration steps (colocalization, PTWAS, and INTACT) can be found in Supplementary Table 10.

## Code Availability

The *regionalpcs* R package, is available through Bioconductor at https://bioconductor.org/packages/release/bioc/html/regionalpcs.html or on GitHub at https://github.com/tyeulalio/regionalpcs. The code used to generate the analyses in this paper are available at https://github.com/tyeulalio/rosmap_methylation_regional_pcs.

## ACKNOWLEDGEMENTS

We extend our gratitude to the participants of the ROSMAP study and their families, whose generosity made this research possible. We acknowledge the Rush Alzheimer’s Disease Center, Rush University Medical Center, Chicago, for providing access to data collected from the Religious Orders Study and the Memory and Aging Project. Special thanks to the National Institute on Aging (NIA) for funding the creation and sharing of the ROSMAP dataset (grants P30AG10161, R01AG15819, R01AG17917, R01AG36836). We extend our sincere appreciation to Jake Chang (Department of Biomedical Data Science, Stanford University) and Chansen Hesia for their significant contributions to this study. Jake Chang played a crucial role in the early development of the instrumental variable analysis. Chansen Hesia provided steadfast support throughout the project’s duration and contributed to the review of the final manuscript. We appreciate the technical support provided by Stanford University and the SCG Informatics Cluster for computational resources. Lastly, we’re grateful to the Bioconductor community for their indispensable tools and forums that supported this work. This research was supported by funding from the National Institute of Aging (R01AG066490, U01AG072573), the National Institute of Health (NIH) T15 NLM007033 and T32AG047126 training grants, and additional NIH grants R01AG066490, U01AG072573, R01MH125244. Additional support was provided from the NIH NHGRI IGVF Program (U01HG012069-03).

## Bibliography

1. Jonathan Cedernaes, Milena Schönke, Jakub Orzechowski Westholm, Jia Mi, Alexander Chibalin, Sarah Voisin, Megan Osler, Heike Vogel, Katarina Hörnaeus, Suzanne L. Dickson, Sara Bergström Lind, Jonas Bergquist, Helgi B Schiöth, Juleen R. Zierath, and Christian Benedict. Acute sleep loss results in tissue-specific alterations in genome-wide dna methylation state and metabolic fuel utilization in humans. Science Advances, 4(8): eaar8590, 8 2018. doi: 10.1126/sciadv.aar8590. publisher: American Association for the Advancement of Science.

2. Alexandra Lahtinen, Sampsa Puttonen, Päivi Vanttola, Katriina Viitasalo, Sonja Sulkava, Natalia Pervjakova, Anni Joensuu, Perttu Salo, Auli Toivola, Mikko Härmä, Lili Milani, Markus Perola, and Tiina Paunio. A distinctive dna methylation pattern in insufficient sleep. Scientific Reports, 9(1):1193, 2 2019. ISSN 2045-2322. doi: 10.1038/s41598-018-38009-0. number: 1 publisher: Nature Publishing Group.

3. Ken Lee and Zdenka Pausova. Cigarette smoking and dna methylation. Frontiers in Genetics, 4, 2013. ISSN 1664-8021. [Online; accessed 2023-09-11].

4. Sonja Zeilinger, Brigitte Kühnel, Norman Klopp, Hansjörg Baurecht, Anja Kleinschmidt, Christian Gieger, Stephan Weidinger, Eva Lattka, Jerzy Adamski, Annette Peters, Konstantin Strauch, Melanie Waldenberger, and Thomas Illig. Tobacco smoking leads to extensive genome-wide changes in dna methylation. PLOS ONE, 8(5):e63812, 5 2013. ISSN 1932-6203. doi: 10.1371/journal.pone.0063812. publisher: Public Library of Science.

5. Marie E. Gaine, Snehajyoti Chatterjee, and Ted Abel. Sleep deprivation and the epigenome. Frontiers in Neural Circuits, 12, 2018. ISSN 1662-5110. [Online; accessed 2023-09-11].

6. Natacha Coppieters, Birger V. Dieriks, Claire Lill, Richard L. M. Faull, Maurice A. Curtis, and Mike Dragunow. Global changes in dna methylation and hydroxymethylation in alzheimer’s disease human brain. Neurobiology of Aging, 35(6):1334–1344, 6 2014. ISSN 0197-4580. doi: 10.1016/j.neurobiolaging.2013.11.031.

7. Philip L. De Jager, Gyan Srivastava, Katie Lunnon, Jeremy Burgess, Leonard C. Schalkwyk, Lei Yu, Matthew L. Eaton, Brendan T. Keenan, Jason Ernst, Cristin McCabe, Anna Tang, Towfique Raj, Joseph Replogle, Wendy Brodeur, Stacey Gabriel, High S. Chai, Curtis Younkin, Steven G. Younkin, Fanggeng Zou, Moshe Szyf, Charles B. Epstein, Julie A. Schneider, Bradley E. Bernstein, Alex Meissner, Nilufer Ertekin-Taner, Lori B. Chibnik, Manolis Kellis, Jonathan Mill, and David A. Bennett. Alzheimer’s disease: early alterations in brain dna methylation at ank1, bin1, rhbdf2 and other loci. Nature Neuroscience, 17(9): 1156–1163, 9 2014. ISSN 1546-1726. doi: 10.1038/nn.3786. PMID: 25129075 PMCID: PMC4292795.

8. Diego Mastroeni, Andrew Grover, Elaine Delvaux, Charisse Whiteside, Paul D. Coleman, and Joseph Rogers. Epigenetic changes in alzheimer’s disease: Decrements in dna methylation. Neurobiology of Aging, 31(12):2025–2037, 12 2010. ISSN 0197-4580. doi: 10.1016/j.neurobiolaging.2008.12.005.

9. Hasan A. Irier and Peng Jin. Dynamics of dna methylation in aging and alzheimer’s disease. DNA and Cell Biology, 31(S1):S–42, 10 2012. ISSN 1044-5498. doi: 10.1089/dna.2011.1565. number-of-pages: S-48 publisher: Mary Ann Liebert, Inc., publishers.

10. Hussain I. Saba and Pierre W. Wijermans. Decitabine in myelodysplastic syndromes. Seminars in Hematology, 42(3 Suppl 2):S23–31, 7 2005. ISSN 0037-1963. doi: 10.1053/j.seminhematol.2005.05.009. PMID: 16015501.

11. Yasuhiro Oki, Etsuko Aoki, and Jean-Pierre J. Issa. Decitabine–bedside to bench. Critical Reviews in Oncology/Hematology, 61(2):140–152, 2 2007. ISSN 1040-8428. doi: 10.1016/j.critrevonc.2006.07.010. PMID: 17023173.

12. Gillian M. Keating. Azacitidine: a review of its use in higher-risk myelodysplastic syndromes/acute myeloid leukaemia. Drugs, 69(17):2501–2518, 2009. ISSN 1179-1950. doi: 10.2165/11202840-000000000-00000. PMID: 19911860.

13. Giuseppe Leone, Maria Teresa Voso, Luciana Teofili, and Michael Lübbert. Inhibitors of dna methylation in the treatment of hematological malignancies and mds. Clinical Immunology (Orlando, Fla.), 109(1):89–102, 10 2003. ISSN 1521-6616. doi: 10.1016/s1521-6616(03)00207-9. PMID: 14585280.

14. Dna methylation analysis as a tool for cell typing. DOI: 10.4161/epi.1.1.2643.

15. Andrew E. Teschendorff and Caroline L. Relton. Statistical and integrative system-level analysis of dna methylation data. Nature Reviews. Genetics, 19(3):129–147, 3 2018. ISSN 1471-0064. doi: 10.1038/nrg.2017.86. PMID: 29129922.

16. Stefan U Kass, Nicoletta Landsberger, and Alan P Wolffe. Dna methylation directs a time-dependent repression of transcription initiation. Current Biology, 7(3):157–165, 3 1997. ISSN 0960-9822. doi: 10.1016/S0960-9822(97)70086-1.

17. Lisa D. Moore, Thuc Le, and Guoping Fan. Dna methylation and its basic function. Neuropsychopharmacology, 38(1):23–38, 1 2013. ISSN 1740-634X. doi: 10.1038/npp.2012. 112. number: 1 publisher: Nature Publishing Group.

18. Netanel Loyfer, Judith Magenheim, Ayelet Peretz, Gordon Cann, Joerg Bredno, Agnes Klochendler, Ilana Fox-Fisher, Sapir Shabi-Porat, Merav Hecht, Tsuria Pelet, Joshua Moss, Zeina Drawshy, Hamed Amini, Patriss Moradi, Sudharani Nagaraju, Dvora Bauman, David Shveiky, Shay Porat, Uri Dior, Gurion Rivkin, Omer Or, Nir Hirshoren, Einat Carmon, Alon Pikarsky, Abed Khalaileh, Gideon Zamir, Ronit Grinbaum, Machmud Abu Gazala, Ido Mizrahi, Noam Shussman, Amit Korach, Ori Wald, Uzi Izhar, Eldad Erez, Vladimir Yutkin, Yaacov Samet, Devorah Rotnemer Golinkin, Kirsty L. Spalding, Henrik Druid, Peter Arner, A. M. James Shapiro, Markus Grompe, Alex Aravanis, Oliver Venn, Arash Jamshidi, Ruth Shemer, Yuval Dor, Benjamin Glaser, and Tommy Kaplan. A dna methylation atlas of normal human cell types. Nature, 613(7943):355–364, 1 2023. ISSN 1476-4687. doi: 10.1038/s41586-022-05580-6. number: 7943 publisher: Nature Publishing Group.

19. Matea Nikolac Perkovic, Alja Videtic Paska, Marcela Konjevod, Katarina Kouter, Dubravka Svob Strac, Gordana Nedic Erjavec, and Nela Pivac. Epigenetics of alzheimer’s disease. Biomolecules, 11(2):195, 2 2021. ISSN 2218-273X. doi: 10.3390/biom11020195. number: 2 publisher: Multidisciplinary Digital Publishing Institute.

20. Illumina. Infinium methylationepic v2.0 beadchip data sheet. 2022.

21. Alexander Meissner, Andreas Gnirke, George W. Bell, Bernard Ramsahoye, Eric S. Lander, and Rudolf Jaenisch. Reduced representation bisulfite sequencing for comparative high-resolution dna methylation analysis. Nucleic Acids Research, 33(18):5868–5877, 10 2005. ISSN 0305-1048. doi: 10.1093/nar/gki901.

22. Swarnaseetha Adusumalli, Mohd Feroz Mohd Omar, Richie Soong, and Touati Benoukraf. Methodological aspects of whole-genome bisulfite sequencing analysis. Briefings in Bioinformatics, 16(3):369–379, 5 2015. ISSN 1467-5463. doi: 10.1093/bib/bbu016.

23. Miren Altuna, Amaya Urdánoz-Casado, Javier Sánchez-Ruiz de Gordoa, María V. Zelaya, Alberto Labarga, Julie M. J. Lepesant, Miren Roldán, Idoia Blanco-Luquin, Álvaro Perdones, Rosa Larumbe, Ivonne Jericó, Carmen Echavarri, Iván Méndez-López, Luisa Di Stefano, and Maite Mendioroz. Dna methylation signature of human hippocampus in alzheimer’s disease is linked to neurogenesis. Clinical Epigenetics, 11(1):91, 6 2019. ISSN 1868-7083. doi: 10.1186/s13148-019-0672-7.

24. Peipei Li, Lee Marshall, Gabriel Oh, Jennifer L. Jakubowski, Daniel Groot, Yu He, Ting Wang, Arturas Petronis, and Viviane Labrie. Epigenetic dysregulation of enhancers in neurons is associated with alzheimer’s disease pathology and cognitive symptoms. Nature Communications, 10(1):2246, 5 2019. ISSN 2041-1723. doi: 10.1038/s41467-019-10101-7. number: 1 publisher: Nature Publishing Group.

25. Lanyu Zhang, Tiago C. Silva, Juan I. Young, Lissette Gomez, Michael A. Schmidt, Kara L. Hamilton-Nelson, Brian W. Kunkle, Xi Chen, Eden R. Martin, and Lily Wang. Epigenome-wide meta-analysis of dna methylation differences in prefrontal cortex implicates the immune processes in alzheimer’s disease. Nature Communications, 11:6114, 11 2020. ISSN 2041-1723. doi: 10.1038/s41467-020-19791-w. PMID: 33257653 PMCID: PMC7704686.

26. Isabel Mendizabal, Stefano Berto, Noriyoshi Usui, Kazuya Toriumi, Paramita Chatterjee, Connor Douglas, Iksoo Huh, Hyeonsoo Jeong, Thomas Layman, Carol A. Tamminga, Todd M. Preuss, Genevieve Konopka, and Soojin V. Yi. Cell type-specific epigenetic links to schizophrenia risk in the brain. Genome Biology, 20(1):135, 7 2019. ISSN 1474-760X. doi: 10.1186/s13059-019-1747-7. PMID: 31288836 PMCID: PMC6617737.

27. Gilles Gasparoni, Sebastian Bultmann, Pavlo Lutsik, Theo F. J. Kraus, Sabrina Sordon, Julia Vlcek, Vanessa Dietinger, Martina Steinmaurer, Melanie Haider, Christopher B. Mulholland, Thomas Arzberger, Sigrun Roeber, Matthias Riemenschneider, Hans A. Kretzschmar, Armin Giese, Heinrich Leonhardt, and Jörn Walter. Dna methylation analysis on purified neurons and glia dissects age and alzheimer’s disease-specific changes in the human cortex. Epigenetics Chromatin, 11(1):41, 7 2018. ISSN 1756-8935. doi: 10.1186/s13072-018-0211-3.

28. Gerald S. Wilkinson, Danielle M. Adams, Amin Haghani, Ake T. Lu, Joseph Zoller, Charles E. Breeze, Bryan D. Arnold, Hope C. Ball, Gerald G. Carter, Lisa Noelle Cooper, Dina K. N. Dechmann, Paolo Devanna, Nicolas J. Fasel, Alexander V. Galazyuk, Linus Günther, Edward Hurme, Gareth Jones, Mirjam Knörnschild, Ella Z. Lattenkamp, Caesar Z. Li, Frieder Mayer, Josephine A. Reinhardt, Rodrigo A. Medellin, Martina Nagy, Brian Pope, Megan L. Power, Roger D. Ransome, Emma C. Teeling, Sonja C. Vernes, Daniel Zamora-Mejías, Joshua Zhang, Paul A. Faure, Lucas J. Greville, L. Gerardo Herrera M., José J. Flores-Martínez, and Steve Horvath. Dna methylation predicts age and provides insight into exceptional longevity of bats. Nature Communications, 12(1):1615, 3 2021. ISSN 2041-1723. doi: 10.1038/s41467-021-21900-2. number: 1 publisher: Nature Publishing Group.

29. Mikko Konki, Maia Malonzo, Ida K. Karlsson, Noora Lindgren, Bishwa Ghimire, Johannes Smolander, Noora M. Scheinin, Miina Ollikainen, Asta Laiho, Laura L. Elo, Tapio Lönnberg, Matias Röyttä, Nancy L. Pedersen, Jaakko Kaprio, Harri Lähdesmäki, Juha O. Rinne, and Riikka J. Lund. Peripheral blood dna methylation differences in twin pairs discordant for alzheimer’s disease. Clinical Epigenetics, 11(1):130, 9 2019. ISSN 1868-7083. doi: 10.1186/s13148-019-0729-7.

30. Timothy J. Peters, Michael J. Buckley, Aaron L. Statham, Ruth Pidsley, Katherine Samaras, Reginald V Lord, Susan J. Clark, and Peter L. Molloy. De novo identification of differentially methylated regions in the human genome. Epigenetics Chromatin, 8(1):6, 1 2015. ISSN 1756-8935. doi: 10.1186/1756-8935-8-6.

31. Andrew E. Jaffe, Peter Murakami, Hwajin Lee, Jeffrey T. Leek, M. Daniele Fallin, Andrew P. Feinberg, and Rafael A. Irizarry. Bump hunting to identify differentially methylated regions in epigenetic epidemiology studies. International Journal of Epidemiology, 41(1): 200–209, 2 2012. ISSN 1464-3685. doi: 10.1093/ije/dyr238. PMID: 22422453 PMCID: PMC3304533.

32. Aaron T.L. Lun and Gordon K. Smyth. De novo detection of differentially bound regions for chip-seq data using peaks and windows: controlling error rates correctly. Nucleic Acids Research, 42(11):e95, 6 2014. ISSN 0305-1048. doi: 10.1093/nar/gku351.

33. Jinpu Cai, Yuyang Xu, Wen Zhang, Shiying Ding, Yuewei Sun, Jingyi Lyu, Meiyu Duan, Shuai Liu, Lan Huang, and Fengfeng Zhou. A comprehensive comparison of residue-level methylation levels with the regression-based gene-level methylation estimations by regear. Briefings in Bioinformatics, 22(4):bbaa253, 7 2021. ISSN 1477-4054. doi: 10.1093/bib/bbaa253.

34. Fabian Müller, Michael Scherer, Yassen Assenov, Pavlo Lutsik, Jörn Walter, Thomas Lengauer, and Christoph Bock. Rnbeads 2.0: comprehensive analysis of dna methylation data. Genome Biology, 20(1):55, 3 2019. ISSN 1474-760X. doi: 10.1186/s13059-019-1664-9. PMID: 30871603 PMCID: PMC6419383.

35. Ting Wang, Weihua Guan, Jerome Lin, Nadia Boutaoui, Glorisa Canino, Jianhua Luo, Juan Carlos Celedón, and Wei Chen. A systematic study of normalization methods for infinium 450k methylation data using whole-genome bisulfite sequencing data. Epigenetics, 10(7):662–669, 6 2015. ISSN 1559-2294. doi: 10.1080/15592294.2015.1057384. PMID: 26036609 PMCID: PMC4623491.

36. Nicole Gull, Michelle R. Jones, Pei-Chen Peng, Simon G. Coetzee, Tiago C. Silva, Jasmine T. Plummer, Alberto Luiz P. Reyes, Brian D. Davis, Stephanie S. Chen, Kate Lawrenson, Jenny Lester, Christine Walsh, Bobbie J. Rimel, Andrew J. Li, Ilana Cass, Yonatan Berg, John-Paul B. Govindavari, Joanna K. L. Rutgers, Benjamin P. Berman, Beth Y. Karlan, and Simon A. Gayther. Dna methylation and transcriptomic features are preserved throughout disease recurrence and chemoresistance in high grade serous ovarian cancers. Journal of Experimental Clinical Cancer Research, 41(1):232, 7 2022. ISSN 1756-9966. doi: 10.1186/s13046-022-02440-z.

37. Sherry Bhalla, Harpreet Kaur, Anjali Dhall, and Gajendra P. S. Raghava. Prediction and analysis of skin cancer progression using genomics profiles of patients. Scientific Reports, 9(1):15790, 10 2019. ISSN 2045-2322. doi: 10.1038/s41598-019-52134-4. number: 1 publisher: Nature Publishing Group.

38. Matthew D. Schultz, Robert J. Schmitz, and Joseph R. Ecker. ‘leveling’ the playing field for analyses of single-base resolution dna methylomes. Trends in Genetics, 28(12):583–585, 12 2012. ISSN 0168-9525. doi: 10.1016/j.tig.2012.10.012.

39. Chantriolnt-Andreas Kapourani and Guido Sanguinetti. Higher order methylation features for clustering and prediction in epigenomic studies. Bioinformatics, 32(17):i405–i412, 9 2016. ISSN 1367-4803. doi: 10.1093/bioinformatics/btw432. publisher: Oxford Academic.

40. Yuanning Zheng, John Jun, Kevin Brennan, and Olivier Gevaert. Epimix is an integrative tool for epigenomic subtyping using dna methylation. Cell Reports Methods, 3(7):100515, 6 2023. ISSN 2667-2375. doi: 10.1016/j.crmeth.2023.100515. PMID: 37533639 PMCID: PMC10391348.

41. Tiffany Eulalio. regionalpcs: Summarizing regional methylation with regional principal components analysis, 2023. DOI: 10.18129/B9.bioc.regionalpcs.

42. Talal Jamil Qazi, Zhenzhen Quan, Asif Mir, and Hong Qing. Epigenetics in alzheimer’s disease: Perspective of dna methylation. Molecular Neurobiology, 55(2):1026–1044, 2 2018. ISSN 1559-1182. doi: 10.1007/s12035-016-0357-6.

43. Philip Scheltens, Bart De Strooper, Miia Kivipelto, Henne Holstege, Gael Chételat, Charlotte E. Teunissen, Jeffrey Cummings, and van der Wiesje M. Flier. Alzheimer’s disease. The Lancet, 397(10284):1577–1590, 4 2021. ISSN 0140-6736, 1474-547X. doi: 10.1016/S0140-6736(20)32205-4. publisher: Elsevier PMID: 33667416.

44. Michael A. DeTure and Dennis W. Dickson. The neuropathological diagnosis of alzheimer’s disease. Molecular Neurodegeneration, 14(1):32, 8 2019. ISSN 1750-1326. doi: 10.1186/s13024-019-0333-5. PMID: 31375134 PMCID: PMC6679484.

45. R.G. Smith, E. Pishva, and G. Shireby. A meta-analysis of epigenome-wide association studies in alzheimer’s disease highlights novel differentially methylated loci across cortex. Nat Commun, 12(3517), 2021. doi: 10.1038/s41467-021-23243-4.

46. Xiaoquan Wen, Roger Pique-Regi, and Francesca Luca. Integrating molecular qtl data into genome-wide genetic association analysis: Probabilistic assessment of enrichment and colocalization. PLOS Genetics, 13(3):e1006646, 3 2017. ISSN 1553-7404. doi: 10.1371/journal.pgen.1006646. publisher: Public Library of Science.

47. Yuhua Zhang, Corbin Quick, Ketian Yu, Alvaro Barbeira, GTEx Consortium, Francesca Luca, Roger Pique-Regi, Hae Kyung Im, and Xiaoquan Wen. Ptwas: investigating tissue-relevant causal molecular mechanisms of complex traits using probabilistic twas analysis. Genome Biology, 21(1):232, 9 2020. ISSN 1474-760X. doi: 10.1186/s13059-020-02026-y. PMID: 32912253 PMCID: PMC7488550.

48. Jeffrey Okamoto, Lijia Wang, Xianyong Yin, Francesca Luca, Roger Pique-Regi, Adam Helms, Hae Kyung Im, Jean Morrison, and Xiaoquan Wen. Probabilistic integration of transcriptome-wide association studies and colocalization analysis identifies key molecular pathways of complex traits. The American Journal of Human Genetics, 110(1):44–57, 1 2023. ISSN 0002-9297. doi: 10.1016/j.ajhg.2022.12.002.

49. Matan Gavish and David L. Donoho. The optimal hard threshold for singular values is 4/sqrt(3). 6 2014. doi: 10.48550/arXiv.1305.5870. 1305.5870 [stat].

50. V A Marcenko and L A Pastur. Distribution of eigenvalues for some sets of random matrices. page 28.

51. Michelle R. Lacey, Carl Baribault, and Melanie Ehrlich. Modeling, simulation and analysis of methylation profiles from reduced representation bisulfite sequencing experiments. Statistical Applications in Genetics and Molecular Biology, 12(6):723–742, 12 2013. ISSN 1544-6115. doi: 10.1515/sagmb-2013-0027. publisher: De Gruyter.

52. David A. Bennett, Aron S. Buchman, Patricia A. Boyle, Lisa L. Barnes, Robert S. Wilson, and Julie A Schneider. Religious orders study and rush memory and aging project. Journal of Alzheimer’s disease : JAD, 64(Suppl 1):S161–S189, 2018. ISSN 1387-2877. doi: 10.3233/JAD-179939. PMID: 29865057 PMCID: PMC6380522.

53. L. B. Chibnik, C. C. White, S. Mukherjee, T. Raj, L. Yu, E. B. Larson, T. J. Montine, C. D. Keene, J. Sonnen, J. A. Schneider, P. K. Crane, J. M. Shulman, D. A. Bennett, and P. L. De Jager. Susceptibility to neurofibrillary tangles: role of the ptprd locus and limited pleiotropy with other neuropathologies. Molecular Psychiatry, 23(6):1521–1529, 6 2018. ISSN 1476-5578. doi: 10.1038/mp.2017.20. number: 6 publisher: Nature Publishing Group PMID: 22471860 PMCID: PMC3409291.

54. H. Braak and E. Braak. Neuropathological stageing of alzheimer-related changes. Acta Neuropathologica, 82(4):239–259, 9 1991. ISSN 1432-0533. doi: 10.1007/BF00308809.

55. S. S. Mirra, A. Heyman, D. McKeel, S. M. Sumi, B. J. Crain, L. M. Brownlee, F. S. Vogel, J. P. Hughes, G. van Belle, and L. Berg. The consortium to establish a registry for alzheimer’s disease (cerad). part ii. standardization of the neuropathologic assessment of alzheimer’s disease. Neurology, 41(4):479–486, 4 1991. ISSN 0028-3878. doi: 10.1212/wnl.41.4.479. PMID: 2011243.

56. Meritxell Oliva, Kathryn Demanelis, Yihao Lu, Meytal Chernoff, Farzana Jasmine, Habibul Ahsan, Muhammad G. Kibriya, Lin S. Chen, and Brandon L. Pierce. Dna methylation qtl mapping across diverse human tissues provides molecular links between genetic variation and complex traits. Nature Genetics, 55(1):112–122, 1 2023. ISSN 1546-1718. doi: 10.1038/s41588-022-01248-z. number: 1 publisher: Nature Publishing Group.

57. Diane E. Bild, David A. Bluemke, Gregory L. Burke, Robert Detrano, Ana V. Diez Roux, Aaron R. Folsom, Philip Greenland, David R. Jacobs Jr., Richard Kronmal, Kiang Liu, Jennifer Clark Nelson, Daniel O’Leary, Mohammed F. Saad, Steven Shea, Moyses Szklo, and Russell P. Tracy. Multi-ethnic study of atherosclerosis: Objectives and design. American Journal of Epidemiology, 156(9):871–881, 11 2002. ISSN 0002-9262. doi: 10.1093/aje/kwf113.

58. Daniel Taliun, Daniel N. Harris, Michael D. Kessler, Jedidiah Carlson, Zachary A. Szpiech, Raul Torres, Sarah A. Gagliano Taliun, André Corvelo Stephanie M. Gogarten, Hyun Min Kang, Achilleas N. Pitsillides, Jonathon LeFaive, Seung-been Lee, Xiaowen Tian, Brian L. Browning, Sayantan Das, Anne-Katrin Emde, Wayne E. Clarke, Douglas P. Loesch, Amol C. Shetty, Thomas W. Blackwell, Albert V. Smith, Quenna Wong, Xiaoming Liu, Matthew P. Conomos, Dean M. Bobo, François Aguet, Christine Albert, Alvaro Alonso, Kristin G. Ardlie, Dan E. Arking, Stella Aslibekyan, Paul L. Auer, John Barnard, R. Graham Barr, Lucas Barwick, Lewis C. Becker, Rebecca L. Beer, Emelia J. Benjamin, Lawrence F. Bielak, John Blangero, Michael Boehnke, Donald W. Bowden, Jennifer A. Brody, Esteban G. Burchard, Brian E. Cade, James F. Casella, Brandon Chalazan, Daniel I. Chasman, Yii-Der Ida Chen, Michael H. Cho, Seung Hoan Choi, Mina K. Chung, Clary B. Clish, Adolfo Correa, Joanne E. Curran, Brian Custer, Dawood Darbar, Michelle Daya, Mariza de Andrade, Dawn L. DeMeo, Susan K. Dutcher, Patrick T. Ellinor, Leslie S. Emery, Celeste Eng, Diane Fatkin, Tasha Fingerlin, Lukas Forer, Myriam Fornage, Nora Franceschini, Christian Fuchsberger, Stephanie M. Fullerton, Soren Germer, Mark T. Gladwin, Daniel J. Gottlieb, Xiuqing Guo, Michael E. Hall, Jiang He, Nancy L. Heard-Costa, Susan R. Heckbert, Marguerite R. Irvin, Jill M. Johnsen, Andrew D. Johnson, Robert Kaplan, Sharon L. R. Kardia, Tanika Kelly, Shannon Kelly, Eimear E. Kenny, Douglas P. Kiel, Robert Klemmer, Barbara A. Konkle, Charles Kooperberg, Anna Köttgen, Leslie A. Lange, Jessica Lasky-Su, Daniel Levy, Xihong Lin, Keng-Han Lin, Chunyu Liu, Ruth J. F. Loos, Lori Garman, Robert Gerszten, Steven A. Lubitz, Kathryn L. Lunetta, Angel C. Y. Mak, Ani Manichaikul, Alisa K. Manning, Rasika A. Mathias, David D. McManus, Stephen T. McGarvey, James B. Meigs, Deborah A. Meyers, Julie L. Mikulla, Mollie A. Minear, Braxton D. Mitchell, Sanghamitra Mohanty, May E. Montasser, Courtney Montgomery, Alanna C. Morrison, Joanne M. Murabito, Andrea Natale, Pradeep Natarajan, Sarah C. Nelson, Kari E. North, Jeffrey R. O’Connell, Nicholette D. Palmer, Nathan Pankratz, Gina M. Peloso, Patricia A. Peyser, Jacob Pleiness, Wendy S. Post, Bruce M. Psaty, D. C. Rao, Susan Redline, Alexander P. Reiner, Dan Roden, Jerome I. Rotter, Ingo Ruczinski, Chloé Sarnowski Sebastian Schoenherr, David A. Schwartz, Jeong-Sun Seo, Sudha Seshadri, Vivien A. Sheehan, Wayne H. Sheu, M. Benjamin Shoemaker, Nicholas L. Smith, Jennifer A. Smith, Nona Sotoodehnia, Adrienne M. Stilp, Weihong Tang, Kent D. Taylor, Marilyn Telen, Timothy A. Thornton, Russell P. Tracy, David J. Van Den Berg, Ramachandran S. Vasan, Karine A. Viaud-Martinez, Scott Vrieze, Daniel E. Weeks, Bruce S. Weir, Scott T. Weiss, Lu-Chen Weng, Cristen J. Willer, Yingze Zhang, Xutong Zhao, Donna K. Arnett, Allison E. Ashley-Koch, Kathleen C. Barnes, Eric Boerwinkle, Stacey Gabriel, Richard Gibbs, Kenneth M. Rice, Stephen S. Rich, Edwin K. Silverman, Pankaj Qasba, Weiniu Gan, George J. Papanicolaou, Deborah A. Nickerson, Sharon R. Browning, Michael C. Zody, Sebastian Zöllner, James G. Wilson, L. Adrienne Cupples, Cathy C. Laurie, Cashell E. Jaquish, Ryan D. Hernandez, Timothy D. O’Connor, and Gonçalo R. Abecasis. Sequencing of 53,831 diverse genomes from the nhlbi topmed program. Nature, 590(7845):290–299, 2 2021. ISSN 1476-4687. doi: 10.1038/s41586-021-03205-y. number: 7845 publisher: Nature Publishing Group.

59. Elior Rahmani, Regev Schweiger, Brooke Rhead, Lindsey A. Criswell, Lisa F. Barcellos, Eleazar Eskin, Saharon Rosset, Sriram Sankararaman, and Eran Halperin. Cell-type-specific resolution epigenetics without the need for cell sorting or single-cell biology. Nature Communications, 10(1):3417, 7 2019. ISSN 2041-1723. doi: 10.1038/s41467-019-11052-9. number: 1 publisher: Nature Publishing Group.

60. Andrew E. Jaffe and Rafael A. Irizarry. Accounting for cellular heterogeneity is critical in epigenome-wide association studies. Genome Biology, 15(2):R31, 2 2014. ISSN 1474-760X. doi: 10.1186/gb-2014-15-2-r31. PMID: 24495553 PMCID: PMC4053810.

61. Yun Liu, Martin J. Aryee, Leonid Padyukov, M. Daniele Fallin, Espen Hesselberg, Arni Runarsson, Lovisa Reinius, Nathalie Acevedo, Margaret Taub, Marcus Ronninger, Klementy Shchetynsky, Annika Scheynius, Juha Kere, Lars Alfredsson, Lars Klareskog, Tomas J. Ekström, and Andrew P. Feinberg. Epigenome-wide association data implicate dna methylation as an intermediary of genetic risk in rheumatoid arthritis. Nature Biotechnology, 31(2):142–147, 2 2013. ISSN 1546-1696. doi: 10.1038/nbt.2487. PMID: 23334450 PMCID: PMC3598632.

62. Andrew E. Teschendorff, Tianyu Zhu, Charles E. Breeze, and Stephan Beck. Episcore: cell type deconvolution of bulk tissue dna methylomes from single-cell rna-seq data. Genome Biology, 21(1):221, 9 2020. ISSN 1474-760X. doi: 10.1186/s13059-020-02126-9. PMID: 32883324 PMCID: PMC7650528.

63. Brandon Jew, Marcus Alvarez, Elior Rahmani, Zong Miao, Arthur Ko, Kristina M. Garske, Jae Hoon Sul, Kirsi H. Pietiläinen, Päivi Pajukanta, and Eran Halperin. Accurate estimation of cell composition in bulk expression through robust integration of single-cell information. Nature Communications, 11(1):1971, 4 2020. ISSN 2041-1723. doi: 10.1038/s41467-020-15816-6. PMID: 32332754 PMCID: PMC7181686.

64. Ellis Patrick, Mariko Taga, Ayla Ergun, Bernard Ng, William Casazza, Maria Cimpean, Christina Yung, Julie A. Schneider, David A. Bennett, Chris Gaiteri, Philip L. De Jager, Elizabeth M. Bradshaw, and Sara Mostafavi. Deconvolving the contributions of cell-type heterogeneity on cortical gene expression. PLoS Computational Biology, 16(8):e1008120, 8 2020. ISSN 1553-734X. doi: 10.1371/journal.pcbi.1008120. PMID: 32804935 PMCID: PMC7451979.

65. Anna-Lena Lang, Tiffany Eulalio, Eddie Fox, Koya Yakabi, Syed A. Bukhari, Claudia H. Kawas, Maria M. Corrada, Stephen B. Montgomery, Frank L. Heppner, David Capper, Daniel Nachun, and Thomas J. Montine. Methylation differences in alzheimer’s disease neuropathologic change in the aged human brain. Acta Neuropathologica Communications, 10(1):174, 11 2022. ISSN 2051-5960. doi: 10.1186/s40478-022-01470-0. PMCID: PMC9710143 PMID: 36447297.

66. Eugene Andres Houseman, William P. Accomando, Devin C. Koestler, Brock C. Christensen, Carmen J. Marsit, Heather H. Nelson, John K. Wiencke, and Karl T. Kelsey. Dna methylation arrays as surrogate measures of cell mixture distribution. BMC bioinformatics, 13:86, 5 2012. ISSN 1471-2105. doi: 10.1186/1471-2105-13-86. PMID: 22568884 PMCID: PMC3532182.

67. Leland McInnes, John Healy, and James Melville. Umap: Uniform manifold approximation and projection for dimension reduction. 9 2020. doi: 10.48550/arXiv.1802.03426. 1802.03426 [cs, stat]x.

68. S. Lloyd. Least squares quantization in pcm. IEEE Transactions on Information Theory, 28 (2):129–137, 3 1982. ISSN 1557-9654. doi: 10.1109/TIT.1982.1056489. event-title: IEEE Transactions on Information Theory.

69. Marie Orre, Willem Kamphuis, Lana M. Osborn, Anne H. P. Jansen, Lieneke Kooijman, Koen Bossers, and Elly M. Hol. Isolation of glia from alzheimer’s mice reveals inflammation and dysfunction. Neurobiology of Aging, 35(12):2746–2760, 12 2014. ISSN 1558-1497. doi: 10.1016/j.neurobiolaging.2014.06.004. PMID: 25002035.

70. Jie Zhao, Tracy O’Connor, and Robert Vassar. The contribution of activated astrocytes to aβ production: implications for alzheimer’s disease pathogenesis. Journal of Neuroinflammation, 8:150, 11 2011. ISSN 1742-2094. doi: 10.1186/1742-2094-8-150. PMID: 22047170 PMCID: PMC3216000.

71. Seonmi Jo, Oleg Yarishkin, Yu Jin Hwang, Ye Eun Chun, Mijeong Park, Dong Ho Woo, Jin Young Bae, Taekeun Kim, Jaekwang Lee, Heejung Chun, Hyun Jung Park, Da Yong Lee, Jinpyo Hong, Hye Yun Kim, Soo-Jin Oh, Seung Ju Park, Hyo Lee, Bo-Eun Yoon, YoungSoo Kim, Yong Jeong, Insop Shim, Yong Chul Bae, Jeiwon Cho, Neil W. Kowall, Hoon Ryu, Eunmi Hwang, Daesoo Kim, and C. Justin Lee. Gaba from reactive astrocytes impairs memory in mouse models of alzheimer’s disease. Nature Medicine, 20(8):886–896, 8 2014. ISSN 1546-170X. doi: 10.1038/nm.3639. number: 8 publisher: Nature Publishing Group PMCID: PMC8385452 PMID: 24973918.

72. Kelly Ceyzériat, Lucile Ben Haim, Audrey Denizot, Dylan Pommier, Marco Matos, Océane Guillemaud, Marie-Ange Palomares, Laurene Abjean, Fanny Petit, Pauline Gipchtein, Marie-Claude Gaillard, Martine Guillermier, Sueva Bernier, Mylène Gaudin, Gwenaëlle Aurégan, Charlène Joséphine, Nathalie Déchamps, Julien Veran, Valentin Langlais, Karine Cambon, Alexis P. Bemelmans, Jan Baijer, Gilles Bonvento, Marc Dhenain, Jean-François Deleuze, Stéphane H. R. Oliet, Emmanuel Brouillet, Philippe Hantraye, Maria-Angeles Carrillo-de Sauvage, Robert Olaso, Aude Panatier, and Carole Escartin. Modulation of astrocyte reactivity improves functional deficits in mouse models of alzheimer’s disease. Acta Neuropathologica Communications, 6(1):104, 10 2018. ISSN 2051-5960. doi: 10.1186/s40478-018-0606-1.

73. Michael D. Monterey, Haichao Wei, Xizi Wu, and Jia Qian Wu. The many faces of astrocytes in alzheimer’s disease. Frontiers in Neurology, 12:619626, 2021. ISSN 1664-2295. doi: 10.3389/fneur.2021.619626. PMID: 34531807 PMCID: PMC8438135.

74. Gautier Koscielny, Peter An, Denise Carvalho-Silva, Jennifer A. Cham, Luca Fumis, Rippa Gasparyan, Samiul Hasan, Nikiforos Karamanis, Michael Maguire, Eliseo Papa, Andrea Pierleoni, Miguel Pignatelli, Theo Platt, Francis Rowland, Priyanka Wankar, A. Patrícia Bento, Tony Burdett, Antonio Fabregat, Simon Forbes, Anna Gaulton, Cristina Yenyxe Gonzalez, Henning Hermjakob, Anne Hersey, Steven Jupe, Şenay Kafkas, Maria Keays, Catherine Leroy, Francisco-Javier Lopez, Maria Paula Magarinos, James Malone, Johanna McEntyre, Alfonso Munoz-Pomer Fuentes, Claire O’Donovan, Irene Papatheodorou, Helen Parkinson, Barbara Palka, Justin Paschall, Robert Petryszak, Naruemon Pratanwanich, Sirarat Sarntivijal, Gary Saunders, Konstantinos Sidiropoulos, Thomas Smith, Zbyslaw Sondka, Oliver Stegle, Y. Amy Tang, Edward Turner, Brendan Vaughan, Olga Vrousgou, Xavier Watkins, Maria-Jesus Martin, Philippe Sanseau, Jessica Vamathevan, Ewan Birney, Jeffrey Barrett, and Ian Dunham. Open targets: a platform for therapeutic target identification and validation. Nucleic Acids Research, 45(Database issue):D985–D994, 1 2017. ISSN 0305-1048. doi: 10.1093/nar/gkw1055. PMID: 27899665 PMCID: PMC5210543.

75. Miguel Tábuas-Pereira, Isabel Santana, Rita Guerreiro, and José Brás. Alzheimer’s disease genetics: Review of novel loci associated with disease. Current Genetic Medicine Reports, 8(1):1–16, 3 2020. ISSN 2167-4876. doi: 10.1007/s40142-020-00182-y.

76. Douglas P. Wightman, Iris E. Jansen, Jeanne E. Savage, Alexey A. Shadrin, Shahram Bahrami, Dominic Holland, Arvid Rongve, Sigrid Børte, Bendik S. Winsvold, Ole Kristian Drange, Amy E. Martinsen, Anne Heidi Skogholt, Cristen Willer, Geir Bråthen, Ingunn Bosnes, Jonas Bille Nielsen, Lars G. Fritsche, Laurent F. Thomas, Linda M. Pedersen, Maiken E. Gabrielsen, Marianne Bakke Johnsen, Tore Wergeland Meisingset, Wei Zhou, Petroula Proitsi, Angela Hodges, Richard Dobson, Latha Velayudhan, Karl Heilbron, Adam Auton, 23 andMe Research Team, Julia M. Sealock, Lea K. Davis, Nancy L. Pedersen, Chandra A. Reynolds, Ida K. Karlsson, Sigurdur Magnusson, Hreinn Stefansson, Steinunn Thordardottir, Palmi V. Jonsson, Jon Snaedal, Anna Zettergren, Ingmar Skoog, Silke Kern, Margda Waern, Henrik Zetterberg, Kaj Blennow, Eystein Stordal, Kristian Hveem, John-Anker Zwart, Lavinia Athanasiu, Per Selnes, Ingvild Saltvedt, Sigrid B. Sando, Ingun Ulstein, Srdjan Djurovic, Tormod Fladby, Dag Aarsland, Geir Selbæk, Stephan Ripke, Kari Stefansson, Ole A. Andreassen, and Danielle Posthuma. A genome-wide association study with 1,126,563 individuals identifies new risk loci for alzheimer’s disease. Nature Genetics, 53(9):1276–1282, 9 2021. ISSN 1546-1718. doi: 10.1038/s41588-021-00921-z. PMID: 34493870 PMCID: PMC10243600.

77. Sarah E. Lacher, Adnan Alazizi, Xuting Wang, Douglas A. Bell, Roger Pique-Regi, Francesca Luca, and Matthew Slattery. A hypermorphic antioxidant response element is associated with increased ms4a6a expression and alzheimer’s disease. Redox Biology, 14:686–693, 10 2017. ISSN 2213-2317. doi: 10.1016/j.redox.2017.10.018. PMID: 29179108 PMCID: PMC5705802.

78. Paul Hollingworth, Denise Harold, Rebecca Sims, Amy Gerrish, Jean-Charles Lambert, Minerva M. Carrasquillo, Richard Abraham, Marian L. Hamshere, Jaspreet Singh Pahwa, Valentina Moskvina, Kimberley Dowzell, Nicola Jones, Alexandra Stretton, Charlene Thomas, Alex Richards, Dobril Ivanov, Caroline Widdowson, Jade Chapman, Simon Lovestone, John Powell, Petroula Proitsi, Michelle K. Lupton, Carol Brayne, David C. Rubinsztein, Michael Gill, Brian Lawlor, Aoibhinn Lynch, Kristelle S. Brown, Peter A. Passmore, David Craig, Bernadette McGuinness, Stephen Todd, Clive Holmes, David Mann, A. David Smith, Helen Beaumont, Donald Warden, Gordon Wilcock, Seth Love, Patrick G. Kehoe, Nigel M. Hooper, Emma R. L. C. Vardy, John Hardy, Simon Mead, Nick C. Fox, Martin Rossor, John Collinge, Wolfgang Maier, Frank Jessen, Eckart Rüther, Britta Schürmann, Reiner Heun, Heike Kölsch, Hendrik van den Bussche, Isabella Heuser, Johannes Kornhuber, Jens Wiltfang, Martin Dichgans, Lutz Frölich, Harald Hampel, John Gallacher, Michael Hüll, Dan Rujescu, Ina Giegling, Alison M. Goate, John S. K. Kauwe, Carlos Cruchaga, Petra Nowotny, John C. Morris, Kevin Mayo, Kristel Sleegers, Karolien Bettens, Sebastiaan Engelborghs, Peter P. De Deyn, Christine Van Broeckhoven, Gill Livingston, Nicholas J. Bass, Hugh Gurling, Andrew McQuillin, Rhian Gwilliam, Panagiotis Deloukas, Ammar Al-Chalabi, Christopher E. Shaw, Magda Tsolaki, Andrew B. Singleton, Rita Guerreiro, Thomas W. Mühleisen, Markus M. Nöthen, Susanne Moebus, Karl-Heinz Jöckel, Norman Klopp, H.-Erich Wichmann, V. Shane Pankratz, Sigrid B. Sando, Jan O. Aasly, Maria Barcikowska, Zbigniew K. Wszolek, Dennis W. Dickson, Neill R. Graff-Radford, Ronald C. Petersen, Alzheimer’s Disease Neuroimaging Initiative, Cornelia M. van Duijn, Monique M. B. Breteler, M. Arfan Ikram, Anita L. DeStefano, Annette L. Fitzpatrick, Oscar Lopez, Lenore J. Launer, Sudha Seshadri, CHARGE consortium, Claudine Berr, Dominique Campion, Jacques Epelbaum, Jean-François Dartigues, Christophe Tzourio, Annick Alpérovitch, Mark Lathrop, EADI1 consortium, Thomas M. Feulner, Patricia Friedrich, Caterina Riehle, Michael Krawczak, Stefan Schreiber, Manuel Mayhaus, S. Nicolhaus, Stefan Wagenpfeil, Stacy Steinberg, Hreinn Stefansson, Kari Stefansson, Jon Snaedal, Sigurbjörn Björnsson, Palmi V. Jonsson, Vincent Chouraki, Benjamin Genier-Boley, Mikko Hiltunen, Hilkka Soininen, Onofre Combarros, Diana Zelenika, Marc Delepine, Maria J. Bullido, Florence Pasquier, Ignacio Mateo, Ana Frank-Garcia, Elisa Porcellini, Olivier Hanon, Eliecer Coto, Victoria Alvarez, Paolo Bosco, Gabriele Siciliano, Michelangelo Mancuso, Francesco Panza, Vincenzo Solfrizzi, Benedetta Nacmias, Sandro Sorbi, Paola Bossù, Paola Piccardi, Beatrice Arosio, Giorgio Annoni, Davide Seripa, Alberto Pilotto, Elio Scarpini, Daniela Galimberti, Alexis Brice, Didier Hannequin, Federico Licastro, Lesley Jones, Peter A. Holmans, Thorlakur Jonsson, Matthias Riemenschneider, Kevin Morgan, Steven G. Younkin, Michael J. Owen, Michael O’Donovan, Philippe Amouyel, and Julie Williams. Common variants at abca7, ms4a6a/ms4a4e, epha1, cd33 and cd2ap are associated with alzheimer’s disease. Nature Genetics, 43(5):429–435, 5 2011. ISSN 1546-1718. doi: 10.1038/ng.803. PMID: 21460840 PMCID: PMC3084173.

79. Abhay Hukku, Matthew G. Sampson, Francesca Luca, Roger Pique-Regi, and Xiaoquan Wen. Analyzing and reconciling colocalization and transcriptome-wide association studies from the perspective of inferential reproducibility. The American Journal of Human Genetics, 109(5):825–837, 5 2022. ISSN 0002-9297. doi: 10.1016/j.ajhg.2022.04.005. PMID: 35523146 PMCID: PMC9118134.

80. Siming Zhao, Wesley Crouse, Sheng Qian, Kaixuan Luo, Matthew Stephens, and Xin He. Adjusting for genetic confounders in transcriptome-wide association studies leads to reliable detection of causal genes. 9 2022. doi: 10.1101/2022.09.27.509700. page: 2022.09.27.509700 section: New Results.

81. Abhay Hukku, Milton Pividori, Francesca Luca, Roger Pique-Regi, Hae Kyung Im, and Xiaoquan Wen. Probabilistic colocalization of genetic variants from complex and molecular traits: promise and limitations. American Journal of Human Genetics, 108(1):25–35, 1 2021. ISSN 1537-6605. doi: 10.1016/j.ajhg.2020.11.012. PMID: 33308443 PMCID: PMC7820626.

82. Nicholas Mancuso, Malika K. Freund, Ruth Johnson, Huwenbo Shi, Gleb Kichaev, Alexander Gusev, and Bogdan Pasaniuc. Probabilistic fine-mapping of transcriptome-wide association studies. Nature Genetics, 51(4):675–682, 4 2019. ISSN 1546-1718. doi: 10.1038/s41588-019-0367-1. PMID: 30926970 PMCID: PMC6619422.

83. Richard Barfield, Helian Feng, Alexander Gusev, Lang Wu, Wei Zheng, Bogdan Pasaniuc, and Peter Kraft. Transcriptome-wide association studies accounting for colocalization using egger regression. Genetic epidemiology, 42(5):418–433, 7 2018. ISSN 0741-0395. doi: 10.1002/gepi.22131. PMID: 29808603 PMCID: PMC6342197.

84. Warren J. Strittmatter and Allen D. Roses. Apolipoprotein e and alzheimer’s disease. Annual Review of Neuroscience, 19(1):53–77, 1996. doi: 10.1146/annurev.ne.19.030196.000413. e print: https://doi.org/10.1146/annurev.ne.19.030196.000413P MID: 8833436.

85. Ornit Chiba-Falek, William K. Gottschalk, and Michael W. Lutz. The effects of the tomm40 poly-t alleles on alzheimer’s disease phenotypes. Alzheimer’s Dementia, 14(5):692–698, 5 2018. ISSN 1552-5260. doi: 10.1016/j.jalz.2018.01.015.

86. Yvonne Shao, McKenzie Shaw, Kaitlin Todd, Maria Khrestian, Giana D’Aleo, P. John Barnard, Jeff Zahratka, Jagan Pillai, Chang-En Yu, C. Dirk Keene, James B. Leverenz, and Lynn M. Bekris. Dna methylation of tomm40-apoe-apoc2 in alzheimer’s disease. Journal of Human Genetics, 63 (4):459–471, 4 2018. ISSN 1435-232X. doi: 10.1038/s10038-017-0393-8. number: 4 publisher: Nature Publishing Group.

87. Devanshi Patel, Xiaoling Zhang, John J. Farrell, Jaeyoon Chung, Thor D. Stein, Kathryn L. Lunetta, and Lindsay A. Farrer. Cell-type-specific expression quantitative trait loci associated with alzheimer disease in blood and brain tissue. Translational Psychiatry, 11(1):250, 4 2021. ISSN 2158-3188. doi: 10.1038/s41398-021-01373-z. PMID: 33907181 PMCID: PMC8079392.

88. Riccardo E. Marioni, Sarah E. Harris, Qian Zhang, Allan F. McRae, Saskia P. Hagenaars, W. David Hill, Gail Davies, Craig W. Ritchie, Catharine R. Gale, John M. Starr, Alison M. Goate, David J. Porteous, Jian Yang, Kathryn L. Evans, Ian J. Deary, Naomi R. Wray, and Peter M. Visscher. Gwas on family history of alzheimer’s disease. Translational Psychiatry, 8(1):1–7, 5 2018. ISSN 2158-3188. doi: 10.1038/s41398-018-0150-6. number: 1 publisher: Nature Publishing Group.

89. Jessica Tulloch, Lesley Leong, Zachary Thomson, Sunny Chen, Eun-Gyung Lee, C. Dirk Keene, Steven P. Millard, and Chang-En Yu. Glia-specific apoe epigenetic changes in the alzheimer’s disease brain. Brain Research, 1698:179–186, 11 2018. ISSN 0006-8993. doi: 10.1016/j.brainres.2018.08.006.

90. Adam M. H. Young, Natsuhiko Kumasaka, Fiona Calvert, Timothy R. Hammond, Andrew Knights, Nikolaos Panousis, Jun Sung Park, Jeremy Schwartzentruber, Jimmy Liu, Kousik Kundu, Michael Segel, Natalia A. Murphy, Christopher E. McMurran, Harry Bulstrode, Jason Correia, Karol P. Budohoski, Alexis Joannides, Mathew R. Guilfoyle, Rikin Trivedi, Ramez Kirollos, Robert Morris, Matthew R. Garnett, Ivan Timofeev, Ibrahim Jalloh, Katherine Holland, Richard Mannion, Richard Mair, Colin Watts, Stephen J. Price, Peter J. Kirkpatrick, Thomas Santarius, Edward Mountjoy, Maya Ghoussaini, Nicole Soranzo, Omer A. Bayraktar, Beth Stevens, Peter J. Hutchinson, Robin J. M. Franklin, and Daniel J. Gaffney. A map of transcriptional heterogeneity and regulatory variation in human microglia. Nature Genetics, 53(6):861–868, 6 2021. ISSN 1546-1718. doi: 10.1038/s41588-021-00875-2. number: 6 publisher: Nature Publishing Group.

91. Txdb.hsapiens.ucsc.hg38.knowngene. [Online; accessed 2023-11-08].

92. Zachary R. McCaw, Jacqueline M. Lane, Richa Saxena, Susan Redline, and Xihong Lin. Operating characteristics of the rank-based inverse normal transformation for quantitative trait analysis in genome-wide association studies. Biometrics, 76(4):1262–1272, 12 2020. ISSN 1541-0420. doi: 10.1111/biom.13214. PMID: 31883270 PMCID: PMC8643141.

93. Matthew E. Ritchie, Belinda Phipson, D. Wu, Yifang Hu, Charity W. Law, Wei Shi, and Gordon K. Smyth. limma powers differential expression analyses for rna-sequencing and microarray studies. Nucleic Acids Research, 43(7):e47, 4 2015. ISSN 1362-4962. doi: 10.1093/nar/gkv007. PMID: 25605792 PMCID: PMC4402510.

94. The whole genome sequence harmonization study; ad knowledge portal. [Online; accessed 2023-09-01].

95. Petr Danecek, James K. Bonfield, Jennifer Liddle, John Marshall, Valeriu Ohan, Martin O. Pollard, Andrew Whitwham, Thomas Keane, Shane A. McCarthy, Robert M. Davies, and Heng Li. Twelve years of samtools and bcftools. GigaScience, 10(2):giab008, 2 2021. ISSN 2047-217X. doi: 10.1093/gigascience/giab008. PMID: 33590861 PMCID: PMC7931819.

96. Picard tools - by broad institute. [Online; accessed 2023-09-01].

97. Christopher C. Chang, Carson C. Chow, Laurent Cam Tellier, Shashaank Vattikuti, Shaun M. Purcell, and James J. Lee. Second-generation plink: rising to the challenge of larger and richer datasets. GigaScience, 4:7, 2015. ISSN 2047-217X. doi: 10.1186/s13742-015-0047-8. PMID: 25722852 PMCID: PMC4342193.

98. Shaun Purcell, Benjamin Neale, Kathe Todd-Brown, Lori Thomas, Manuel A. R. Ferreira, David Bender, Julian Maller, Pamela Sklar, Paul I. W. de Bakker, Mark J. Daly, and Pak C. Sham. Plink: A tool set for whole-genome association and population-based linkage analyses. American Journal of Human Genetics, 81(3):559–575, 9 2007. ISSN 0002-9297. PMID: 17701901 PMCID: PMC1950838.

99. Alkes L. Price, Nick J. Patterson, Robert M. Plenge, Michael E. Weinblatt, Nancy A. Shadick, and David Reich. Principal components analysis corrects for stratification in genome-wide association studies. Nature Genetics, 38(8):904–909, 8 2006. ISSN 1061-4036. doi: 10.1038/ng1847. PMID: 16862161.

100. Mirna Safieh, Amos D. Korczyn, and Daniel M. Michaelson. Apoe4: an emerging therapeutic target for alzheimer’s disease. BMC Medicine, 17(1):64, 3 2019. ISSN 1741-7015. doi: 10.1186/s12916-019-1299-4.

101. Martin J. Aryee, Andrew E. Jaffe, Hector Corrada-Bravo, Christine Ladd-Acosta, Andrew P. Feinberg, Kasper D. Hansen, and Rafael A. Irizarry. Minfi: a flexible and comprehensive bioconductor package for the analysis of infinium dna methylation microarrays. Bioinformatics, 30(10):1363– 1369, 5 2014. ISSN 1367-4803. doi: 10.1093/bioinformatics/btu049.

102. S. T. Sherry, M.-H. Ward, M. Kholodov, J. Baker, L. Phan, E. M. Smigielski, and K. Sirotkin. dbsnp: the ncbi database of genetic variation. Nucleic Acids Research, 29(1):308–311, 1 2001. ISSN 0305-1048. PMID: 11125122 PMCID: PMC29783.

103. Katarzyna Murat, Björn Grüning, Paulina Wiktoria Poterlowicz, Gillian Westgate, Desmond J. Tobin, and Krzysztof Poterlowicz. Ewastools: Infinium human methylation beadchip pipeline for population epigenetics integrated into galaxy. GigaScience, 9(5):giaa049, 5 2020. ISSN 2047-217X. doi: 10.1093/gigascience/giaa049. PMID: 32401319 PMCID: PMC7219210.

104. Ruth Pidsley, Chloe C. Y Wong, Manuela Volta, Katie Lunnon, Jonathan Mill, and Leonard C. Schalkwyk. A data-driven approach to preprocessing illumina 450k methylation array data. BMC genomics, 14:293, 5 2013. ISSN 1471-2164. doi: 10.1186/1471-2164-14-293. PMID: 23631413 PMCID: PMC3769145.

105. Andrew E. Teschendorff, Francesco Marabita, Matthias Lechner, Thomas Bartlett, Jesper Tegner, David Gomez-Cabrero, and Stephan Beck. A beta-mixture quantile normalization method for correcting probe design bias in illumina infinium 450 k dna methylation data. Bioinformatics (Oxford, England), 29(2):189–196, 1 2013. ISSN 1367-4811. doi: 10.1093/bioinformatics/bts680. PMID: 23175756 PMCID: PMC3546795.

106. Pan Du, Xiao Zhang, Chiang-Ching Huang, Nadereh Jafari, Warren A. Kibbe, Lifang Hou, and Simon M. Lin. Comparison of beta-value and m-value methods for quantifying methylation levels by microarray analysis. BMC Bioinformatics, 11(1):587, 11 2010. ISSN 1471-2105. doi: 10.1186/1471-2105-11-587.

107. Jeroen van Rooij, Pooja R. Mandaviya, Annique Claringbould, Janine F. Felix, Jenny van Dongen, Rick Jansen, Lude Franke, Peter A. C. ‘t Hoen, Bas Heijmans, and Joyce B. J. van Meurs. Evaluation of commonly used analysis strategies for epigenome- and transcriptome-wide association studies through replication of large-scale population studies. Genome Biology, 20:235, 11 2019. ISSN 1474-7596. doi: 10.1186/s13059-019-1878-x. PMID: 31727104 PMCID: PMC6857161.

108. Jian Tang, Jingzhou Liu, Ming Zhang, and Qiaozhu Mei. Visualizing large-scale and high-dimensional data. pages 287–297, 4 2016. doi: 10.1145/2872427.2883041. 1602.00370 [cs].

109. Kevin Blighe. Pcatools: everything principal component analysis, 9 2023. original-date: 2018-11-06T15:37:13Z.

110. Raymond G. Cavalcante and Maureen A. Sartor. annotatr: genomic regions in context. Bioinformatics (Oxford, England), 33(15):2381–2383, 8 2017. ISSN 1367-4811. doi: 10.1093/bioinformatics/btx183. PMID: 28369316 PMCID: PMC5860117.

111. Adam Frankish, Mark Diekhans, Anne-Maud Ferreira, Rory Johnson, Irwin Jungreis, Jane Loveland, Jonathan M. Mudge, Cristina Sisu, James Wright, Joel Armstrong, If Barnes, Andrew Berry, Alexandra Bignell, Silvia Carbonell Sala, Jacqueline Chrast, Fiona Cunningham, Tomás Di Domenico, Sarah Donaldson, Ian T. Fiddes, Carlos García Girón, Jose Manuel Gonzalez, Tiago Grego, Matthew Hardy, Thibaut Hourlier, Toby Hunt, Osagie G. Izuogu, Julien Lagarde, Fergal J. Martin, Laura Martínez, Shamika Mohanan, Paul Muir, Fabio C. P. Navarro, Anne Parker, Baikang Pei, Fernando Pozo, Magali Ruffier, Bianca M. Schmitt, Eloise Stapleton, Marie-Marthe Suner, Irina Sycheva, Barbara Uszczynska-Ratajczak, Jinuri Xu, Andrew Yates, Daniel Zerbino, Yan Zhang, Bronwen Aken, Jyoti S. Choudhary, Mark Gerstein, Roderic Guigó, Tim J. P. Hub-bard, Manolis Kellis, Benedict Paten, Alexandre Reymond, Michael L. Tress, and Paul Flicek. Gencode reference annotation for the human and mouse genomes. Nucleic Acids Research, 47 (D1):D766–D773, 1 2019. ISSN 1362-4962. doi: 10.1093/nar/gky955. PMID: 30357393 PMCID: PMC6323946.

112. Donna Karolchik, Angela S. Hinrichs, Terrence S. Furey, Krishna M. Roskin, Charles W. Sugnet, David Haussler, and W. James Kent. The ucsc table browser data retrieval tool. Nucleic Acids Research, 32(Database issue):D493–496, 1 2004. ISSN 1362-4962. doi: 10.1093/nar/gkh103. PMID: 14681465 PMCID: PMC308837.

113. Michael Lawrence, Wolfgang Huber, Hervé Pagès, Patrick Aboyoun, Marc Carlson, Robert Gentleman, Martin T. Morgan, and Vincent J. Carey. Software for computing and annotating genomic ranges. PLoS computational biology, 9(8):e1003118, 2013. ISSN 1553-7358. doi: 10.1371/journal.pcbi.1003118. PMID: 23950696 PMCID: PMC3738458.

114. liftover: Changing genomic coordinate systems with rtracklayer::liftover. [Online; accessed 2023-09-06].

115. Amaro Taylor-Weiner, François Aguet, Nicholas J. Haradhvala, Sager Gosai, Shankara Anand, Jaegil Kim, Kristin Ardlie, Eliezer M. Van Allen, and Gad Getz. Scaling computational genomics to millions of individuals with gpus. Genome Biology, 20(1):228, 11 2019. ISSN 1474-760X. doi: 10.1186/s13059-019-1836-7.

116. Halit Ongen, Alfonso Buil, Andrew Anand Brown, Emmanouil T. Dermitzakis, and Olivier Delaneau. Fast and efficient qtl mapper for thousands of molecular phenotypes. Bioinformatics, 32 (10):1479–1485, 5 2016. ISSN 1367-4803. doi: 10.1093/bioinformatics/btv722.

117. Molecular qtl discovery incorporating genomic annotations using bayesian false discovery rate control. The Annals of Applied Statistics, 10(3):1619–1638, 9 2016. ISSN 1932-6157, 1941-7330. doi: 10.1214/16-AOAS952.

118. Yeji Lee, Francesca Luca, Roger Pique-Regi, and Xiaoquan Wen. Bayesian multi-snp genetic association analysis: Control of fdr and use of summary statistics. 5 2018. doi: 10.1101/316471. page: 316471 section: New Results.

119. James W. MacDonald, Tabitha Harrison, Theo K. Bammler, Nicholas Mancuso, and Sara Lindström. An updated map of grch38 linkage disequilibrium blocks based on european ancestry data. 3 2022. doi: 10.1101/2022.03.04.483057. page: 2022.03.04.483057 section: New Results.

120. Corbin Quick, Xiaoquan Wen, Gonçalo Abecasis, Michael Boehnke, and Hyun Min Kang. Integrating comprehensive functional annotations to boost power and accuracy in gene-based association analysis. PLOS Genetics, 16(12):e1009060, 12 2020. ISSN 1553-7404. doi: 10.1371/journal.pgen.1009060. publisher: Public Library of Science.

121. 1000 Genomes Project Consortium, Adam Auton, Lisa D. Brooks, Richard M. Durbin, Erik P. Garrison, Hyun Min Kang, Jan O. Korbel, Jonathan L. Marchini, Shane McCarthy, Gil A. McVean, and Gonçalo R. Abecasis. A global reference for human genetic variation. Nature, 526(7571):68–74, 10 2015. ISSN 1476-4687. doi: 10.1038/nature15393. PMID: 26432245 PMCID: PMC4750478.

