## Supplementary Figures and Table Captions for "*regionalpcs*: improved discovery of DNA methylation associations with complex traits"

### This PDF file includes:

Supplementary Figures 1 to 14
Legends for Supplementary Tables 1 to 9

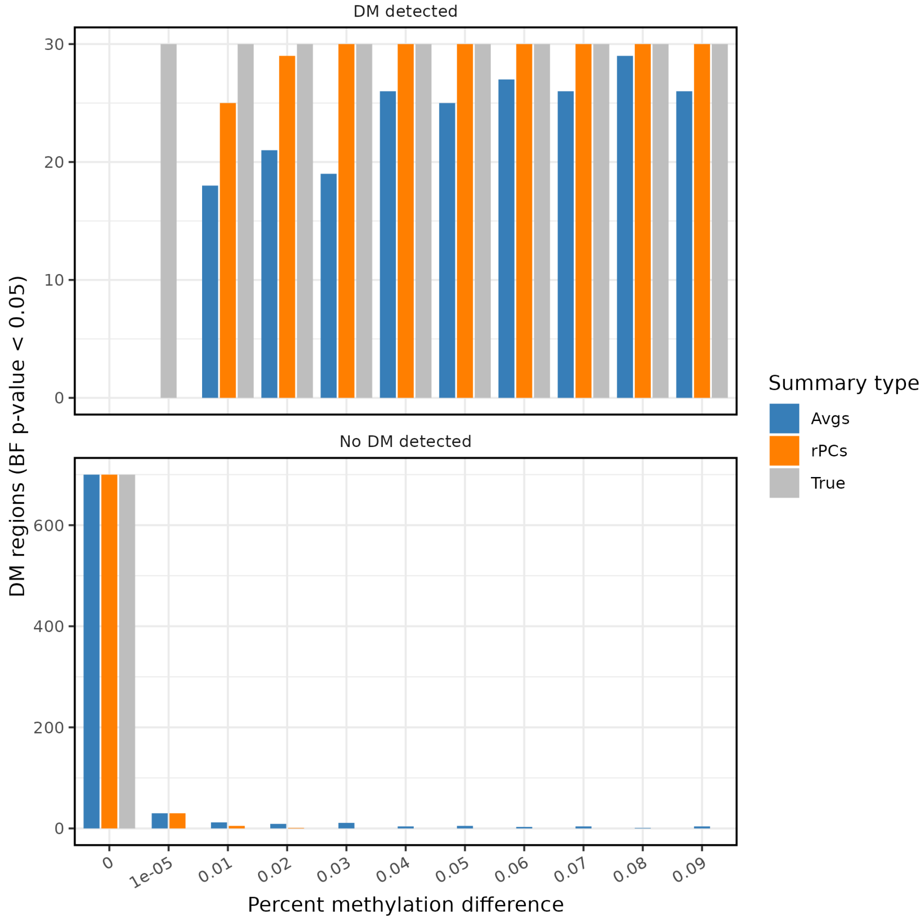


**Supplementary Figure 1. Differential methylation detection in simulated regions.** Bar plots comparing differential methylation detected in simulated regions using averages (blue) and regionalpcs (orange), with true differential methylation counts indicated by grey bars. The x-axis details the extent of methylation differences between cases and controls. The upper plot displays counts of regions identified as differentially methylated, while the lower plot enumerates regions not detected as differentially methylated.

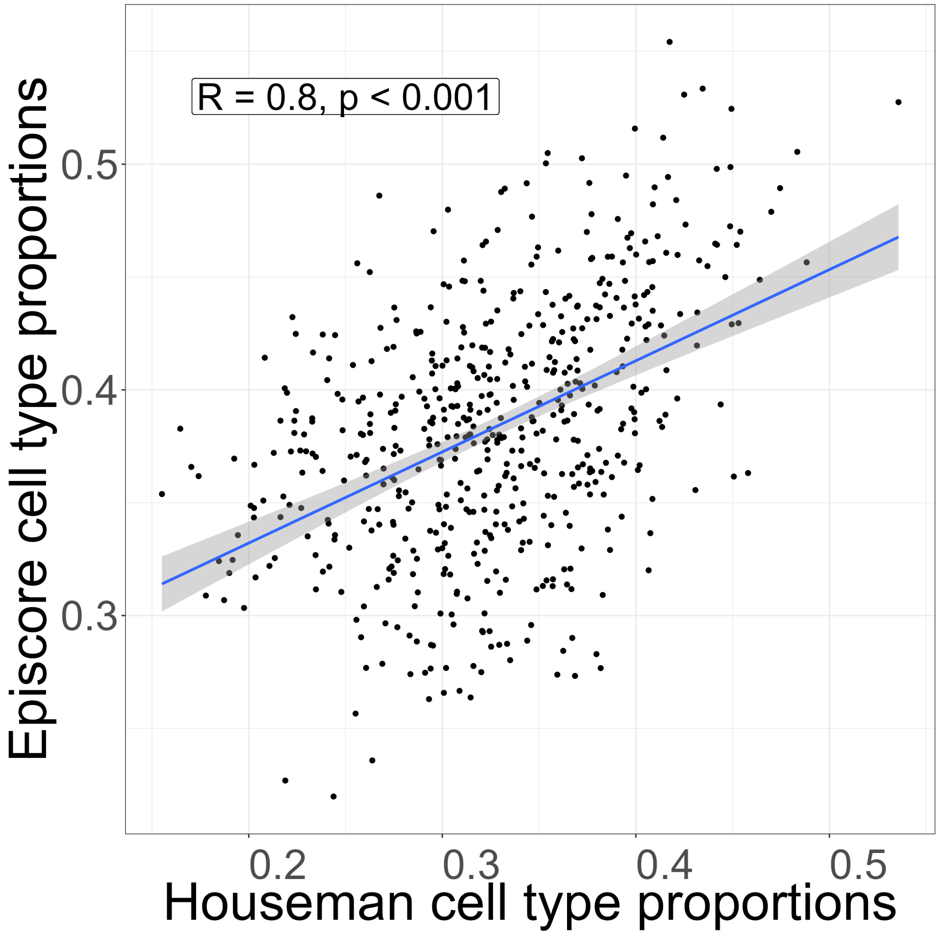


**Supplementary Figure 2. Correlation analysis of Houseman vs. EpiSCORE methods.** The plot displays neuron proportion estimates with the Houseman method on the x-axis and the EpiSCORE method on the y-axis, demonstrating strong agreement (Pearson $r = 0.8, p = 1.2 \times 10^{-128}$)

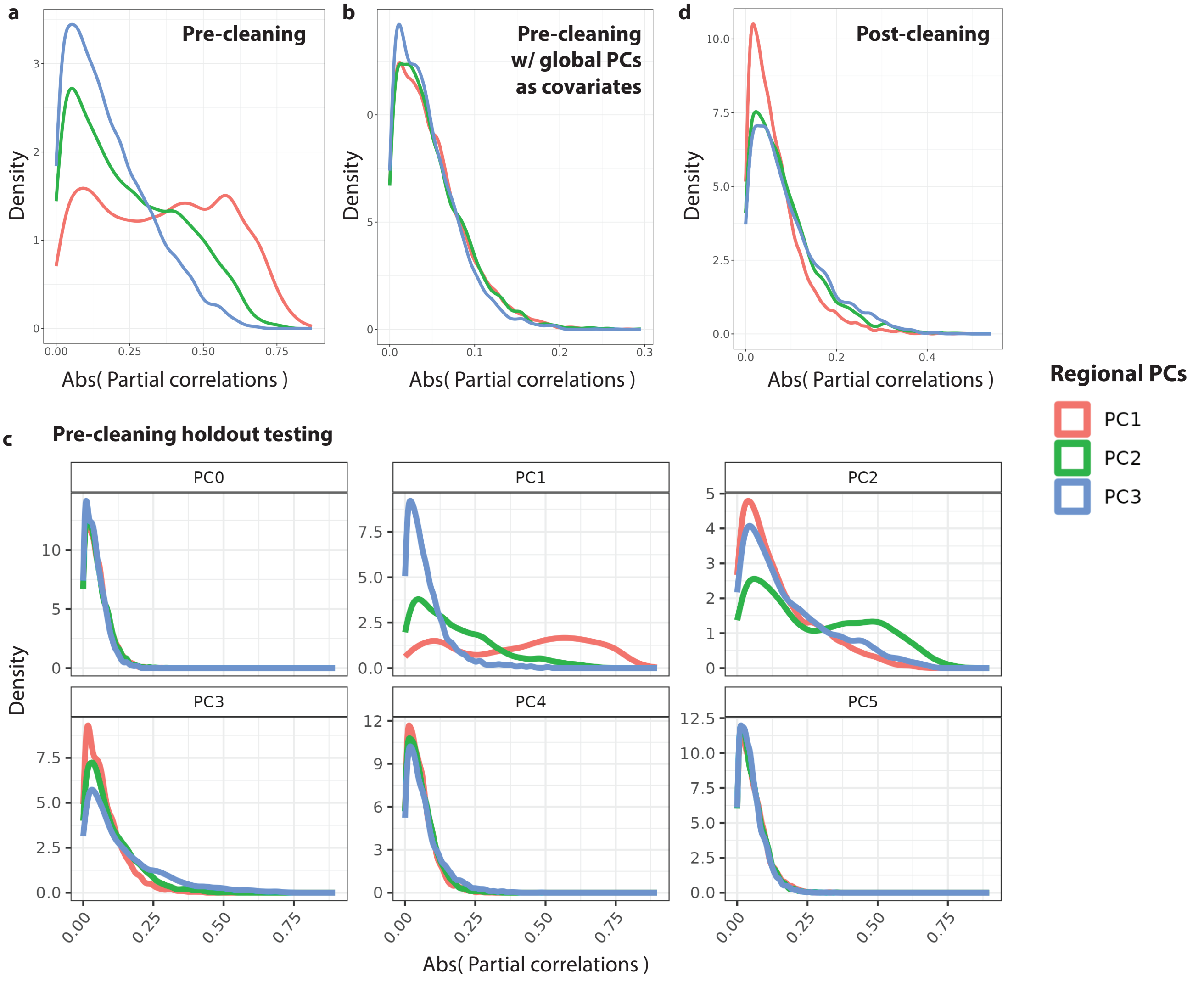


**Supplementary Figure 3. Analysis of rPCs' independence from global signals.** Showing representative results using the full gene region and astrocytes cell type. (a) absolute partial correlations for rPC1, rPC2, and rPC3, controlling for confounders like sex, age, and cell type proportions. Higher correlations in rPC1 and rPC2 suggest potential capture of global signals. (b) Partial correlations incorporating global methylation PCs and confounders, showing consistent results across rPCs, indicative of regional signal specificity. (c) Holdout method results, iterating the exclusion of each global PC (N = 27) from the model. Facets represent the excluded global PC, with variations in distributions for PCs 1-4, implying their global signals overlap with rPCs. Minimal effects observed for global PCs numbered five and above, similar to the baseline (PC 0). (d) Partial correlations calculated on rPCs after removing global PCs 1-4 from the methylation data. Only confounders were included in this model, displaying uniform distributions across rPCs and confirming the effective removal of global influences, thus validating the regional specificity of rPCs.

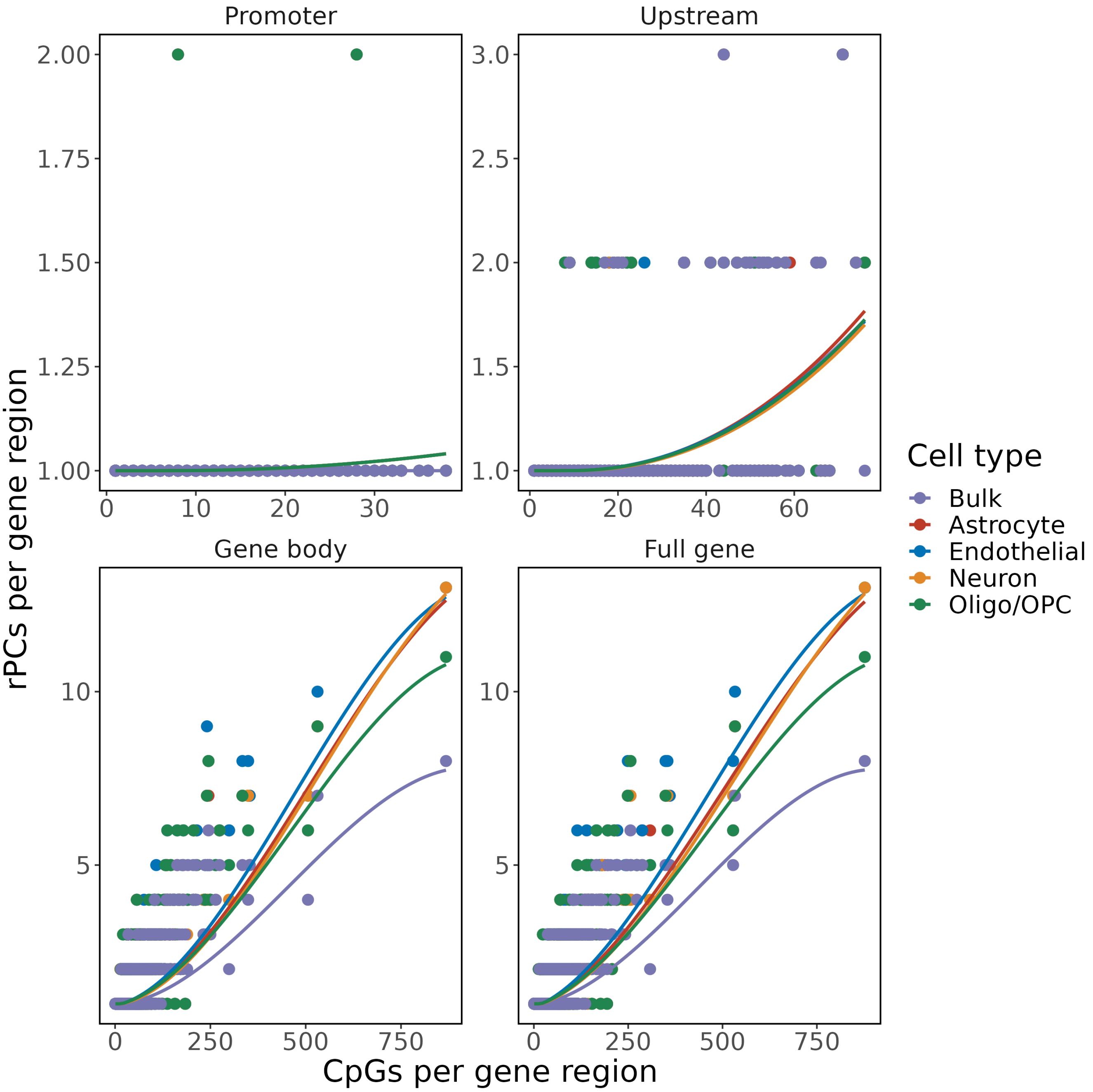


**Supplementary Figure 4. Correlation between CpGs and rPCs in different genome regions.** Scatter plots show the relationship between the count of CpGs and the required number of rPCs for gene representation across promoter, 5kb upstream, and gene body regions. Color-coded by cell type, with LOESS lines tracing the trend for each group. Generally, promoters and 5kb upstream regions require fewer than 3 rPCs, while gene bodies and full genes need more.

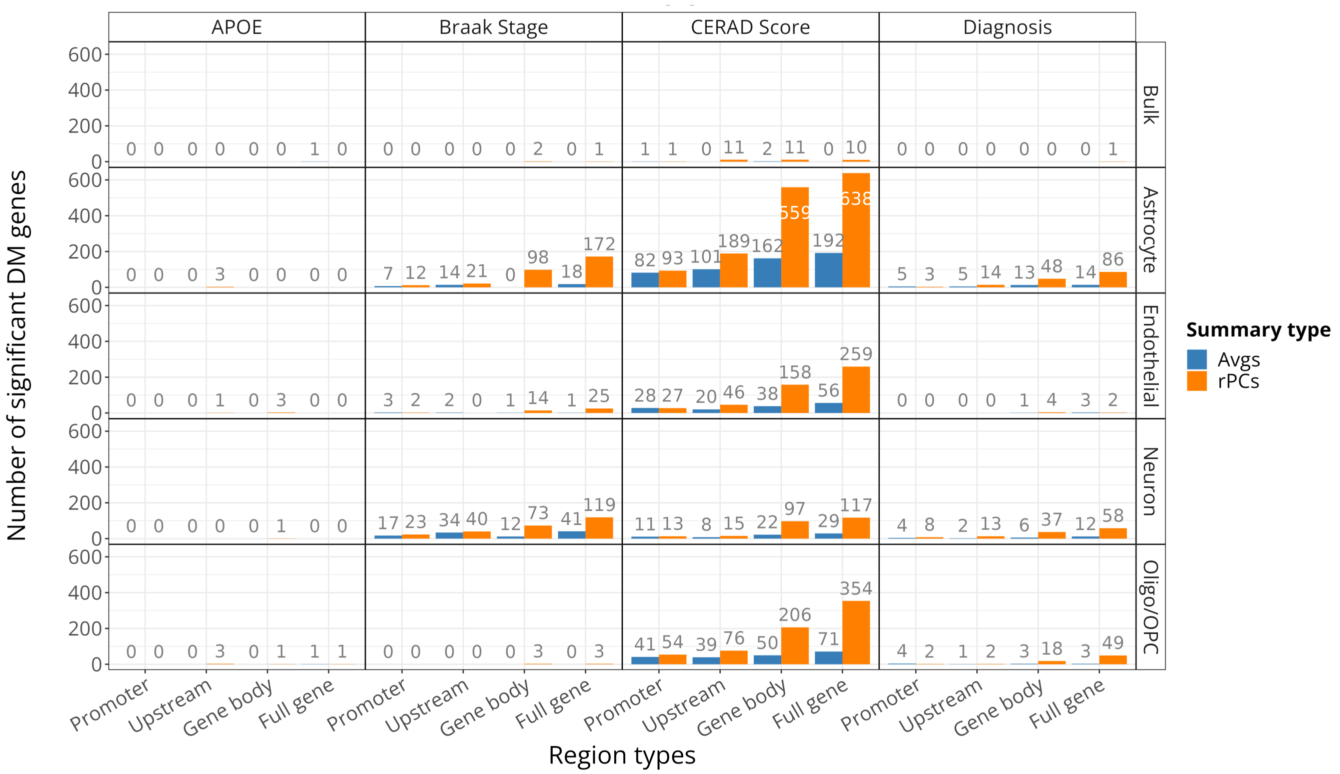


**Supplementary Figure 5. Comparing differential methylation in disease phenotypes using averages and rPCs.** This figure demonstrates a comparative analysis across cell types and genomic regions for four disease outcome phenotypes, using averages (blue) and regional principal components (rPCs, orange). Key observations include: (1) limited identification of disease-associated genes in bulk cells, with astrocytes presenting the most significant findings; (2) fewer genes linked to *APOE* genotype and Alzheimer's disease diagnosis, but notable number associated with CERAD score; (3) a pronounced increase in disease-associated genes detected within full gene and gene body regions, compared to promoters and 5kb upstream areas; (4) a consistent pattern of rPCs identifying a higher number of genes with disease-associated methylation changes than averages, underscoring the effectiveness of rPCs in methylation analysis.

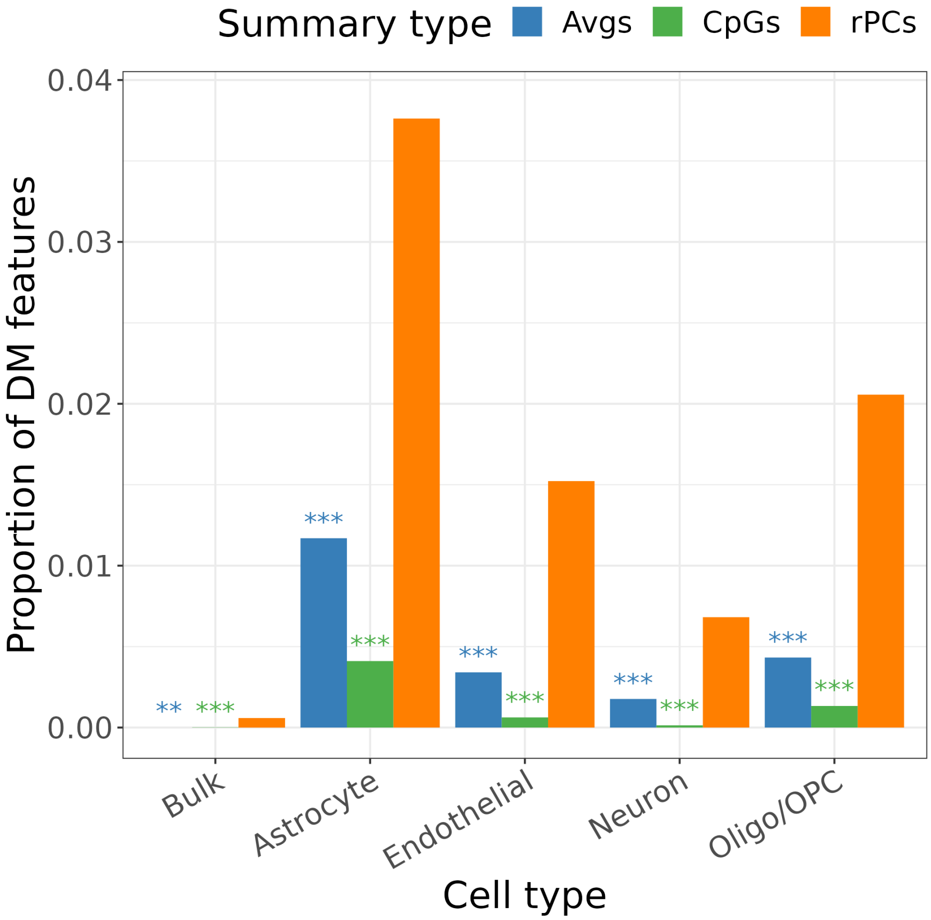


**Supplementary Figure 6. Efficiency of methylation feature detection in neuritic plaques.** This bar graph contrasts the effectiveness of regional principal components (rPCs), averages, and CpG-level feature analysis in identifying neuritic plaque-associated methylation features across different cell types. rPCs notably excel, detecting a significantly larger proportion of methylation features relevant to neuritic plaques compared to other methods. This demonstrates the enhanced ability of rPCs in capturing methylation changes associated with neuritic plaques. Statistical significance is indicated by Bonferroni-adjusted $p$-values from Fisher's test comparing the proportion of differentially methylated features detected against rPCs. (***$p < 0.001$, **$p < 0.01$).

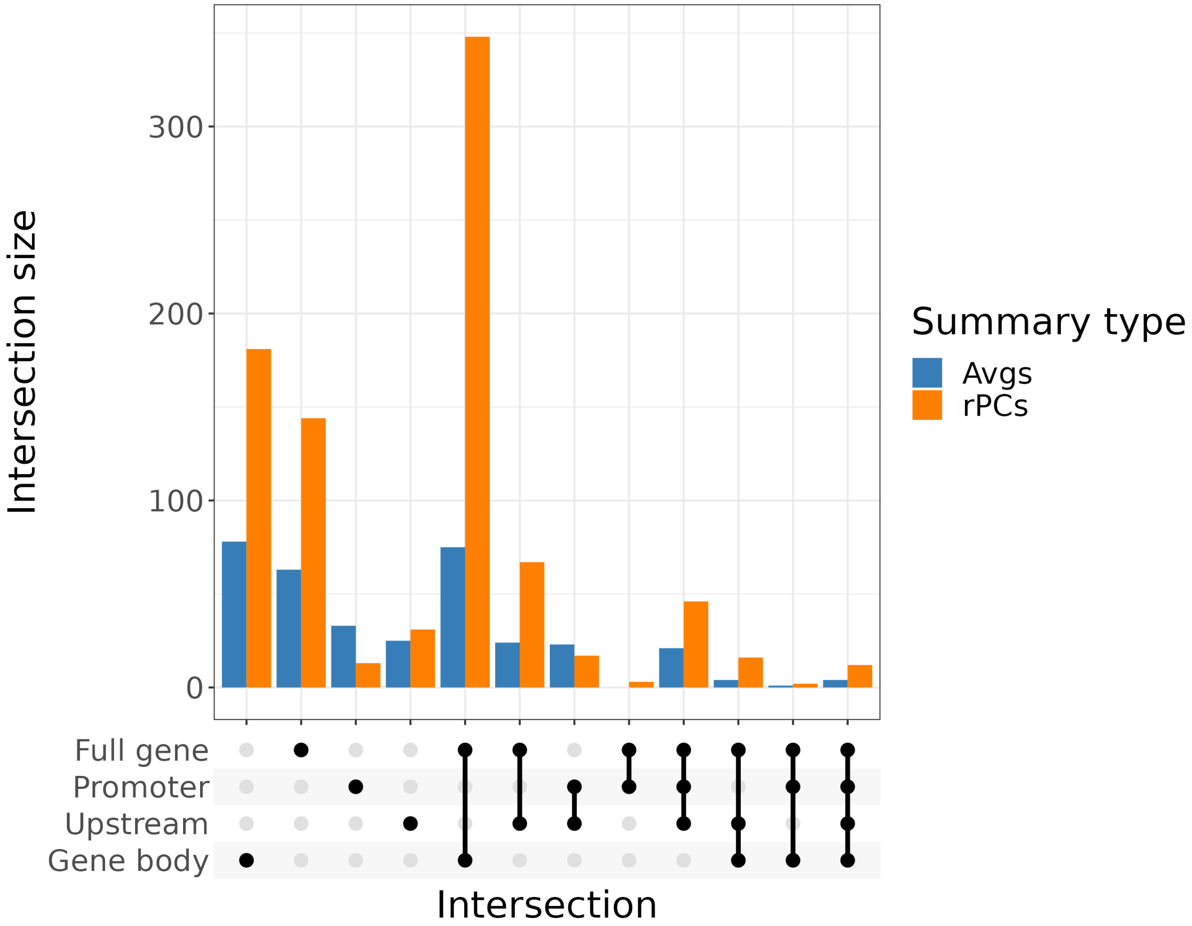


**Supplementary Figure 7. Intersection of neuritic-plaque associated genes across region types.** UpSet plot illustrating the intersections of genes with neuritic-plaque associated methylation levels across various region types in astrocytes, identified through differential methylation analysis (BH $p$-value < 0.05). The x-axis displays the selected intersecting region type groups, chosen based on their size and arranged according to the degree of group intersections. Genes pinpointed by degree 1 and 2 intersections constitute a large portion when either averages or rPCs are used for methylation summarization.

 

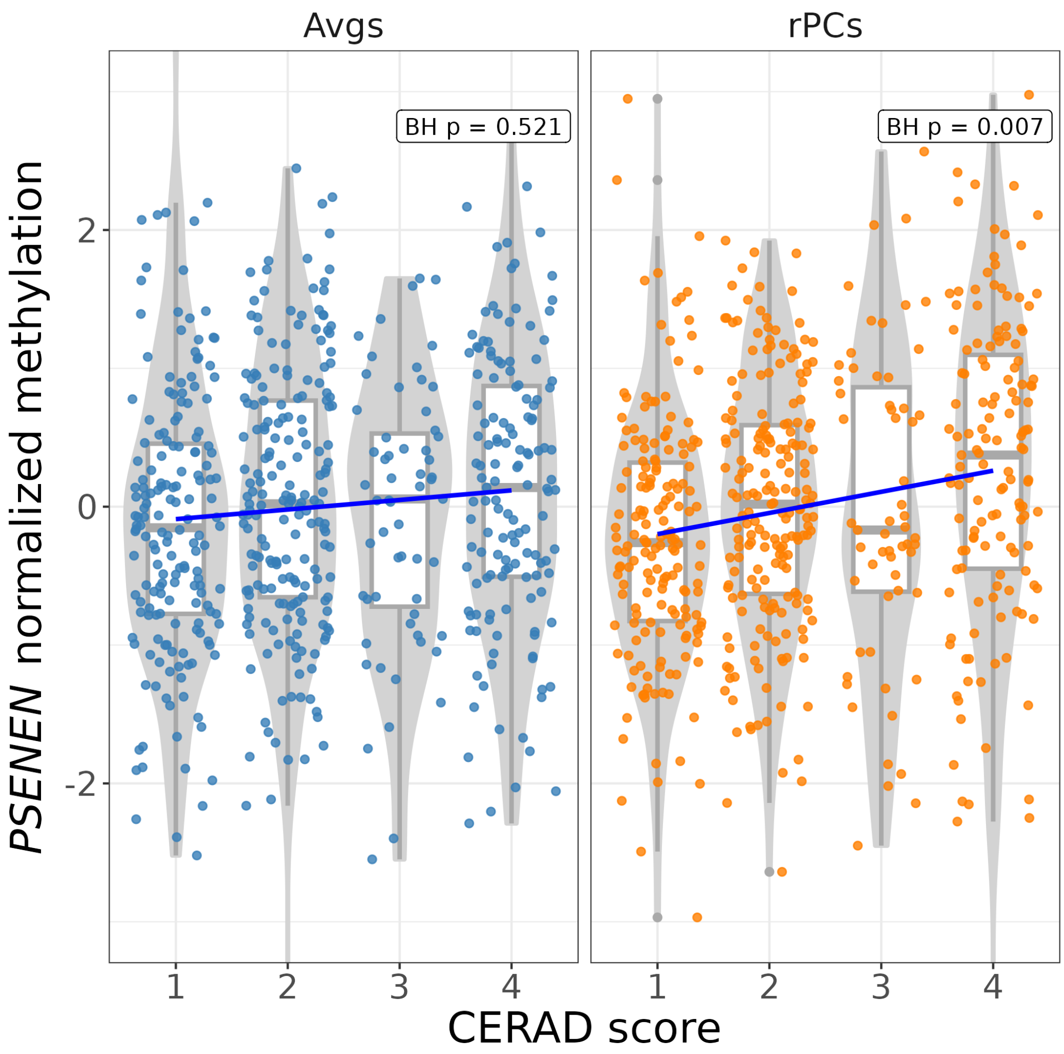


**Supplementary Figure 8.** ***PSENEN* methylation across CERAD scores in astrocytes.** This violin plot presents the gene-level methylation distribution of *PSENEN* in astrocytes, summarized using averages and regional principal components (rPCs), across different CERAD score groups. The distinct pattern observed with rPCs led to the identification of *PSENEN* as differentially methylated, a finding not captured with averages, underscoring the sensitivity of the rPCs method. (BH $p$ = Benjamini-Hochberg adjusted $p$-value).

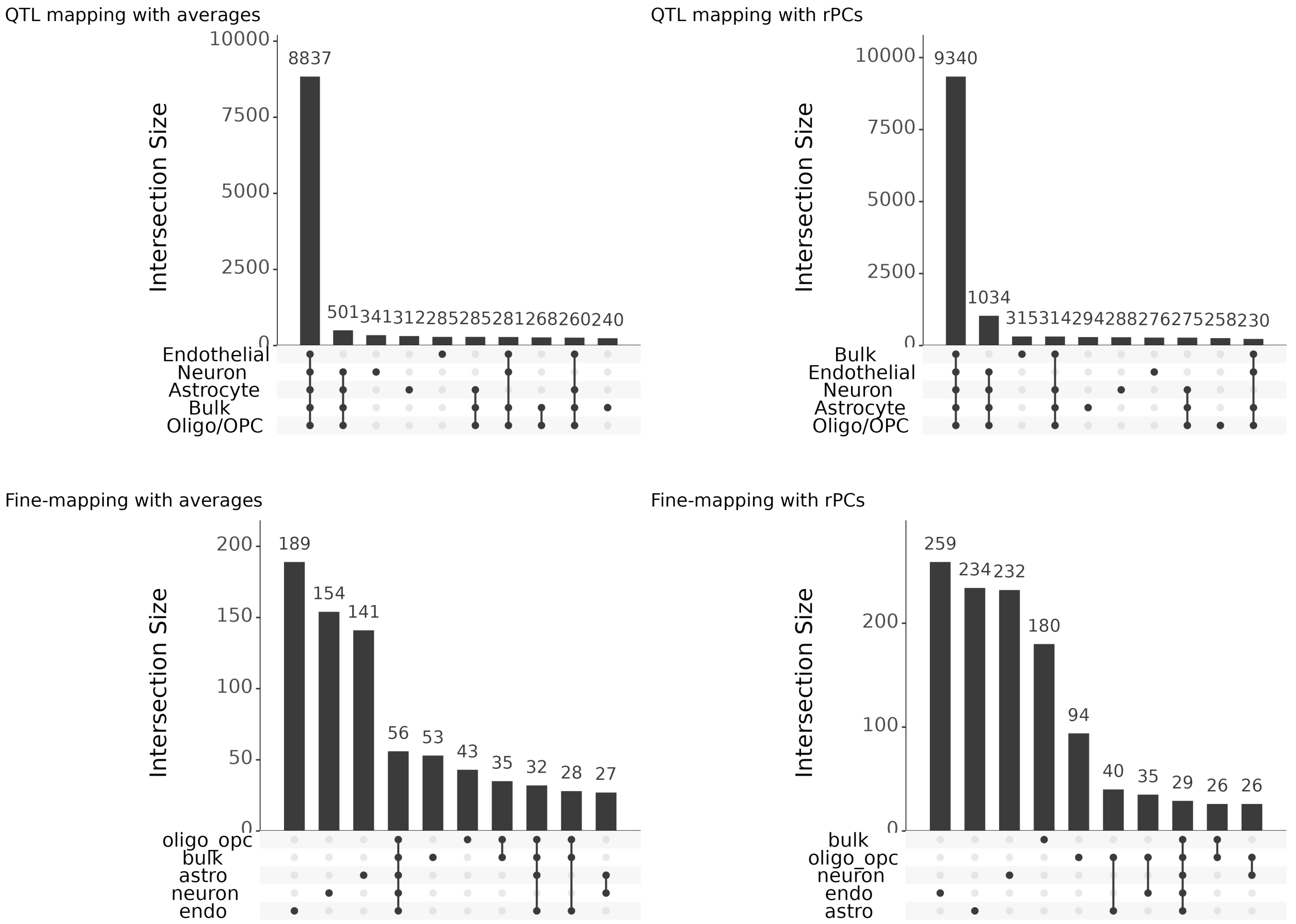


**Supplementary Figure 9. Overlap of meGenes in QTL mapping and fine-mapping.** This figure illustrates the comparative overlap of meGenes identified in QTL mapping before and after fine-mapping. The top row displayed meGenes discovered through QTL mapping using averages (left) and rPCs (right), demonstrating the general lack of specificity in this step. In contrast, the bottom row, with averages (left) and rPCs (right) after fine-mapping, reveals a clear emergence of cell type-specificity achieved through fine-mapping.

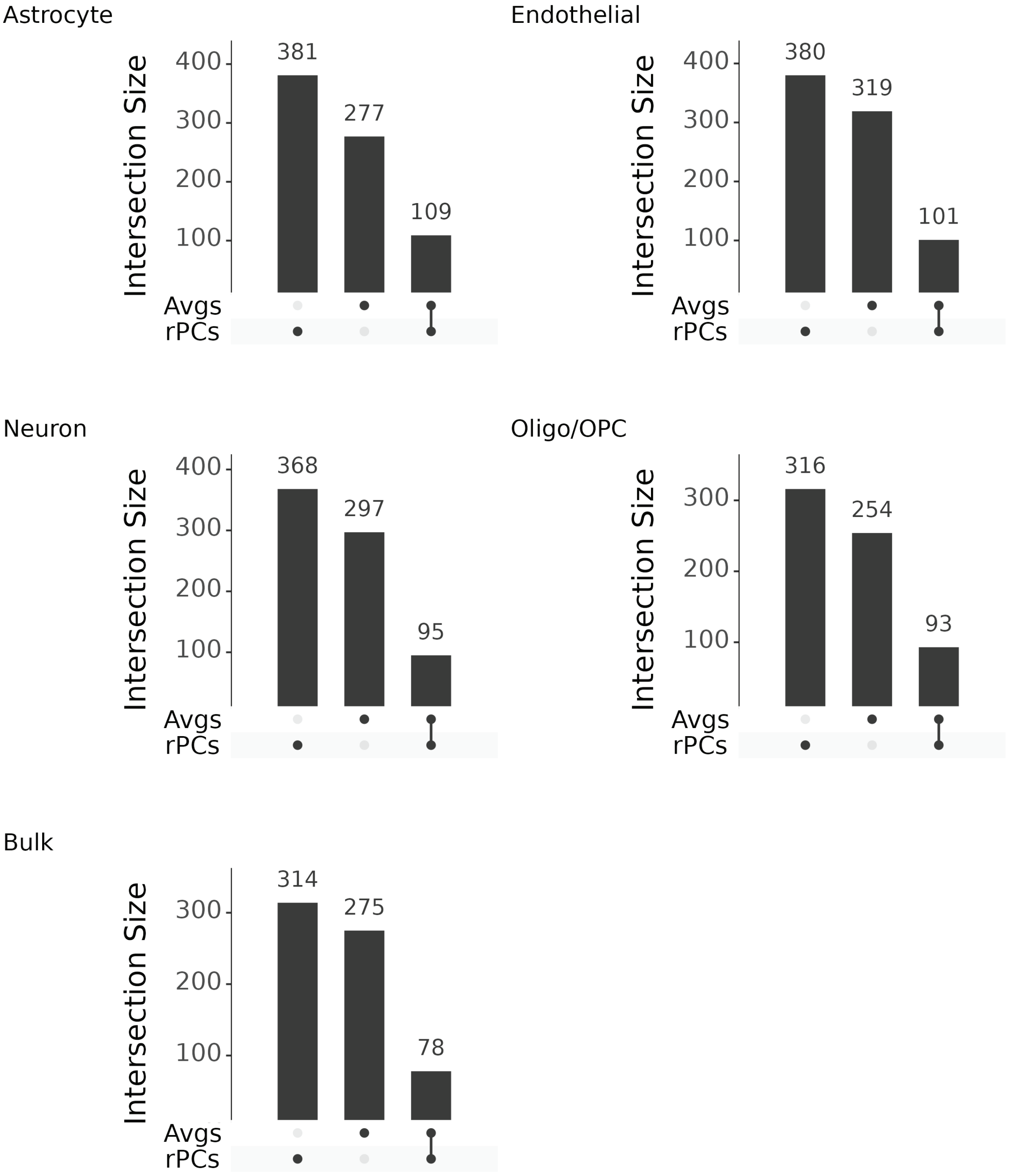


**Supplementary Figure 10. Intersection of meGenes identified by fine-mapping meQTLs.** UpSet plots depict the intersection of fine-mapped meGenes using averages and regional principal components (rPCs) for summarizing full gene methylation. The limited overlap in genes identified by both methods highlights the importance of incorporating diverse summary approaches in meQTL analysis, emphasizing the unique insights offered by each method.

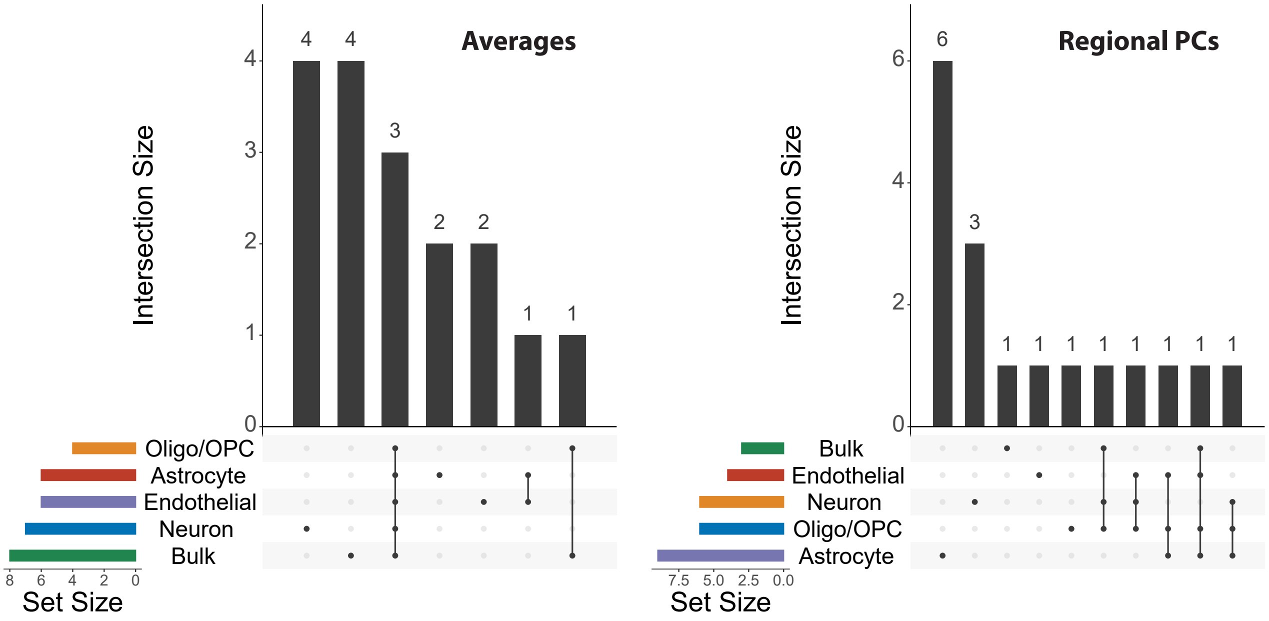


**Supplementary Figure 11. Distinct cell type intersection in colocalization analyses.** UpSet plots demonstrate the unique results across various cell types colocalization analyses as part of GWAS integration. The plots highlight the full gene region, with the majority of the colocalized genes identified being specific to individual cell types.

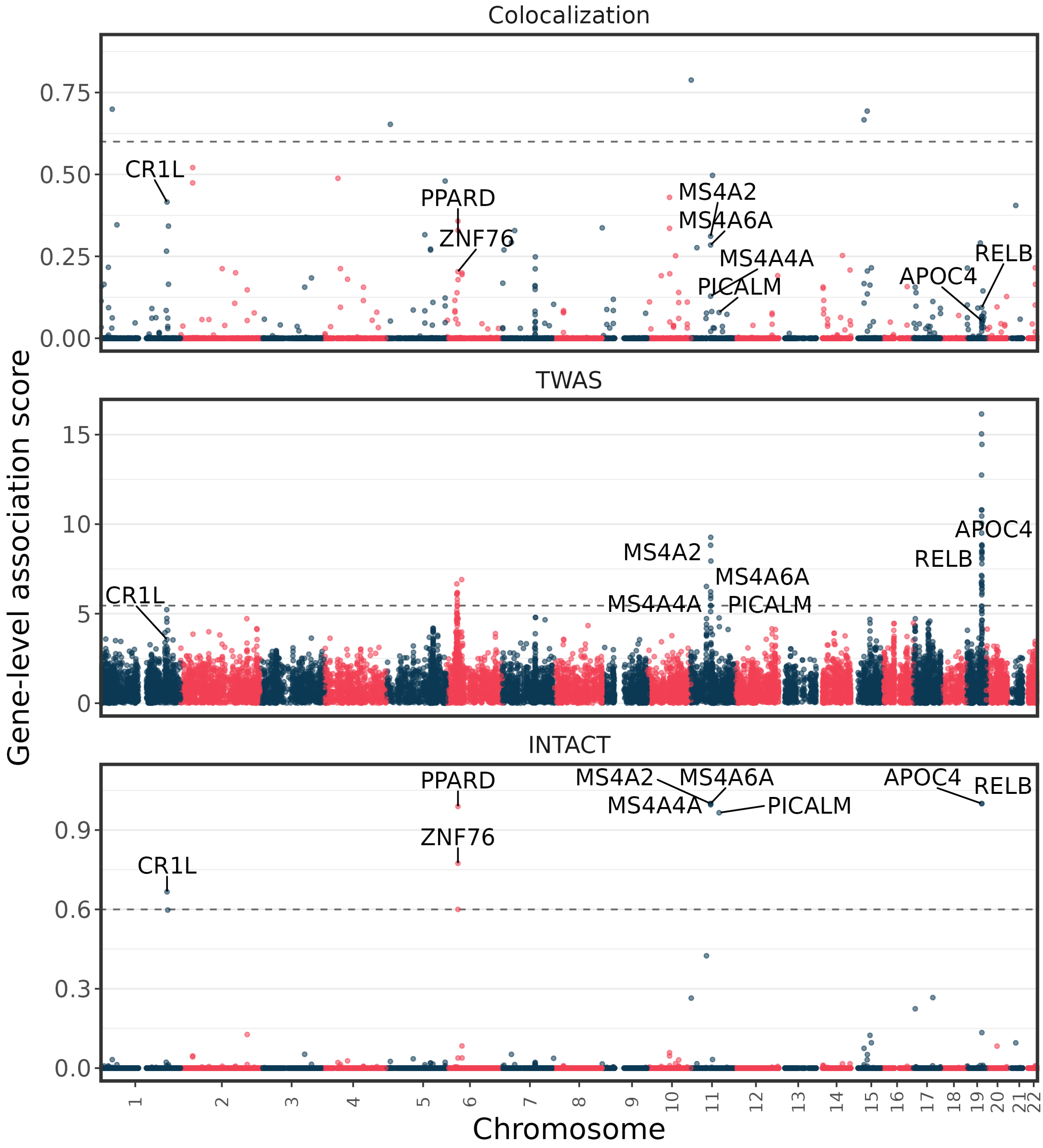


**Supplementary Figure 12. Gene associations in neuronal regions via GWAS integration.** Manhatton plots display gene-level associations in neuronal full gene regions, analyzed through colocalization, instrumental variable analysis using TWAS, and INTACT methods. Association strength is measured on the y-axis, using gene-level colocalization probability (GLCP) for colocalization, z-score for TWAS, and posterior probability for INTACT. Significance thresholds are marked by horizontal dashed lines at 0.6 for colocalization and INTACT, and a z-score of 5.45 for genome-wide significance in TWAS. Genes exceeding INTACT significance threshold are identified, highlighting those with genetic variants potentially influencing Alzheimer's disease risk through methylation changes.

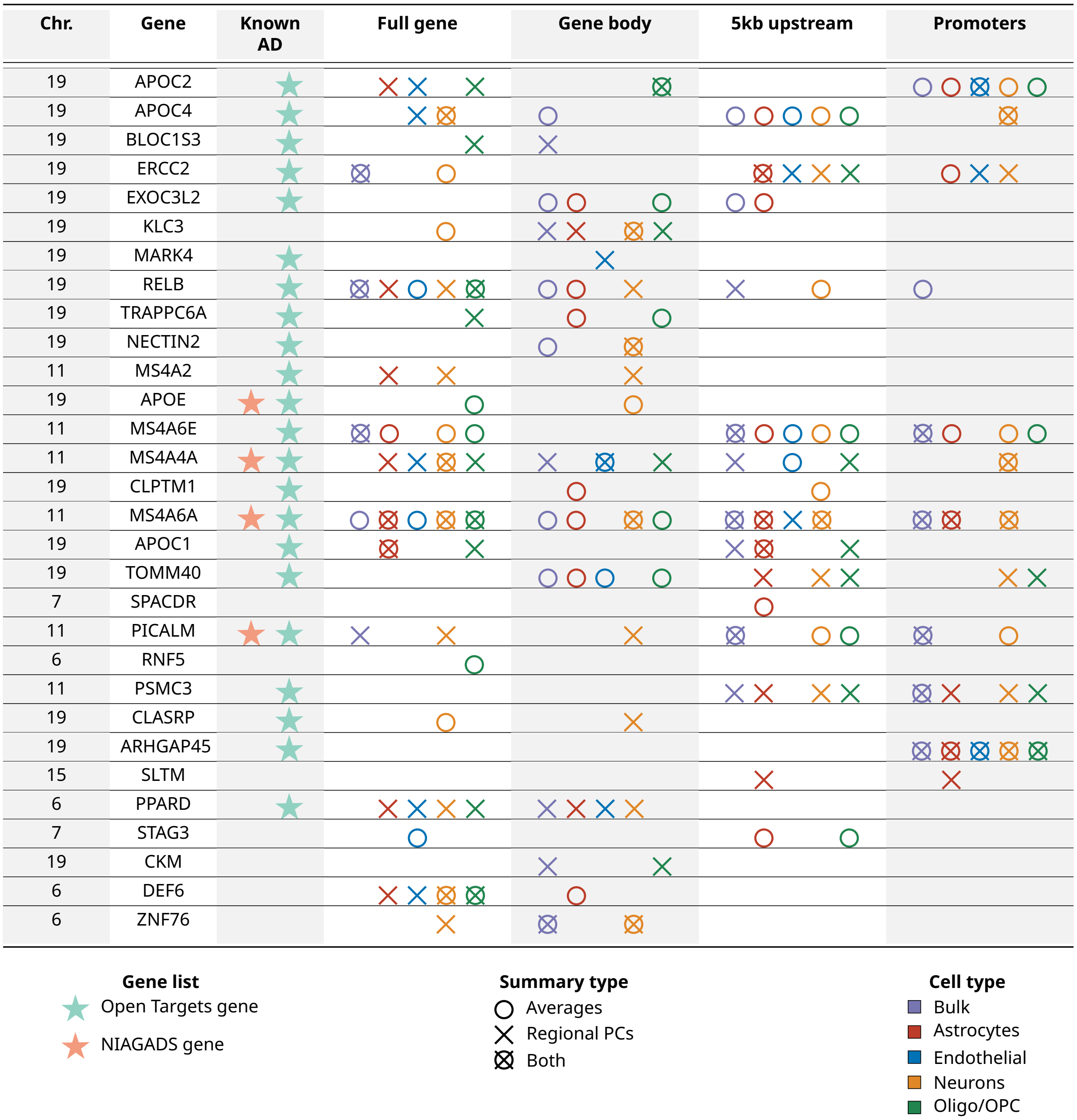


**Supplementary Figure 13. Candidate genes linking methylation and Alzheimer's disease risk across gene region types.** The table highlights genes with shared genetic effects on methylation and Alzheimer's disease susceptibility, which are identified across various region types. Using rPCs, these genes were identified with INTACT posterior probabilities over 0.6, integrating findings from colocalization and PTWAS. Genes detected using regionalpcs are marked with 'X', while those by averages are indicated by 'O', each color-coded by cell type. The chromosome on which each gene is specified in the far-left column, labeled 'Chr.'.

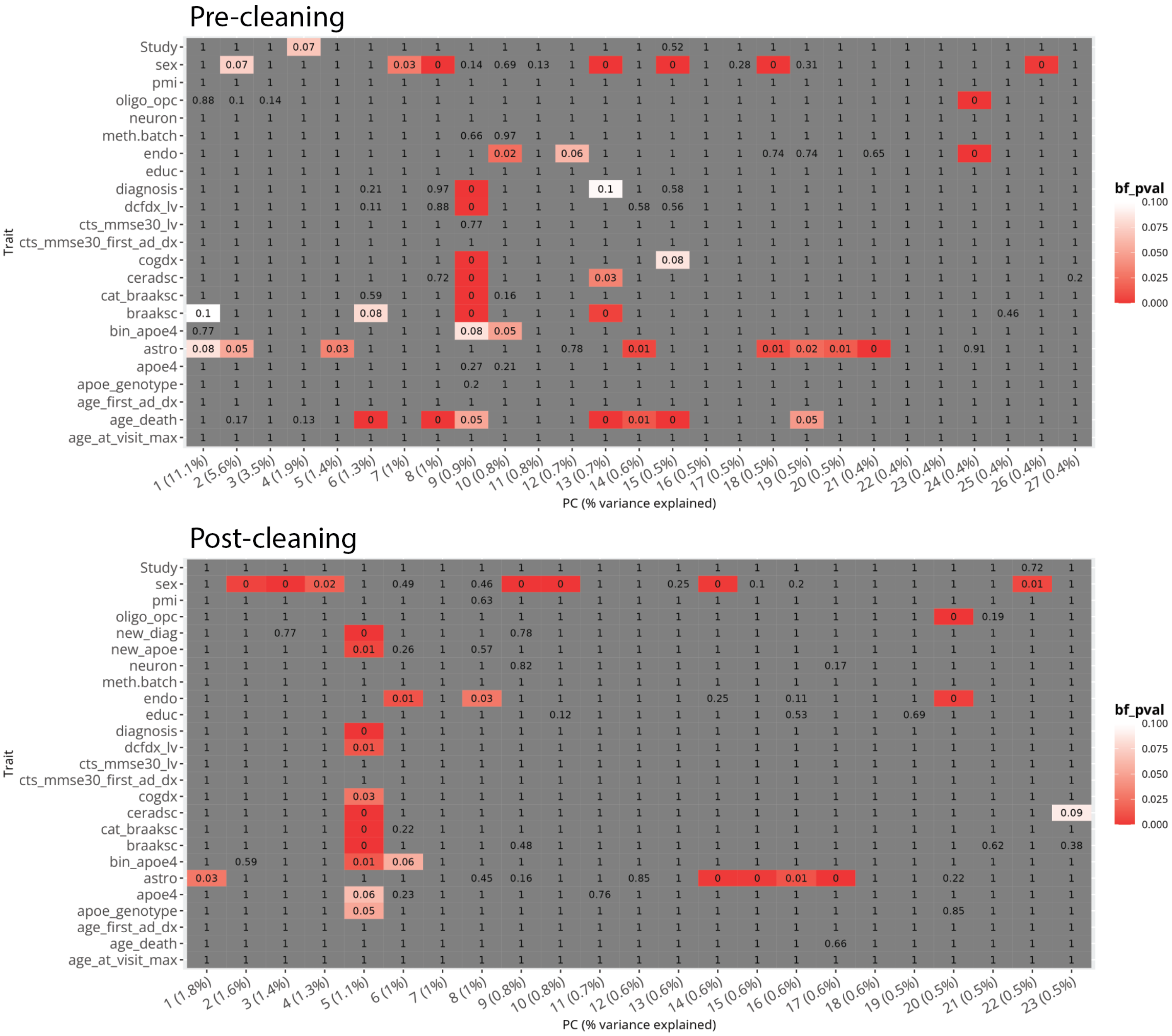


**Supplementary Figure 14. Correlations between traits and global methylation PCs in astrocytes.** Heatmaps display the correlation between various traits and global methylation principal components (PCs) in astrocytes, represented by Bonferroni-adjusted p-values for multiple PC comparisons. The top heatmap illustrates correlations before data cleaning, identifying variables for removal to reduce potential confounders. The bottom heatmap shows correlations post-removal of these confounding variables, providing a clearer view of the intrinsic relationships between traits and global methylation PCs.

**Supplementary Table 1. Summary Statistics for Differential Methylation Performance in Simulation Analysis.** This table outlines a structured comparison of simulation parameters and their corresponding values, alongside detailed summary statistics for differential methylation performance. 'parameter_name' denotes the variable under scrutiny, while 'parameter_value' gives the assigned value for statistical summary. 'diff_*' conveys the discrepancies between regions flagged as differentially methylated by regional principal components (rPCs) versus averages (avgs), with the calculation derived from their differences. 'avgs_*' and 'rpcs_*' columns provide a detailed account of the proportion of regions identified as differentially methylated by averages and rPCs, respectively. The encompassing summary statistics span minimum (min), median (med), mean, and maximum (max) for each considered parameter, offering an exhaustive quantification of the method's analytical scope.

**Supplementary Table 2. Cohort trait distribution in the ROSMAP study.** Traits across 563 selected samples from the ROSMAP cohort are distributed with individual diagnoses derived from clinical assessments and neuropathological evaluations, including CERAD scores and Braak stages. Alzheimer's disease (AD) classification is ascribed to individuals showing concordant dementia diagnosis and neuropathology. Those devoid of both conditions are classified as Controls, while cases presenting discrepant findings are noted as Mixed Pathology*

**Supplementary Table 3. K-means clustering results of deconvolved and sorted cell types.** This representation conveys the cluster assignments from K-means clustering, comparing four deconvolved cell types against two validation cell types sorted by experimental measures, specifically NeuN and Olig2. Inconsistencies are highlighted in red, flagging samples that diverged from their projected cell type clusters. Alignments between predicted and actual cell types are marked in green. Each column enumerates the cluster label designated to the individual samples.

**Supplementary Table 4. Descriptive Analysis of Methylation Features and rPCs Variance.** Presented are the comprehensive statistics for the counts of methylation features per gene and the percentage of variance accounted for by rPCs within distinct genomic regions and cellular classifications. The parameter 'total_genes' denotes the total gene count summarized for each specified region and cell type. 'Total_cpgs' indicates the complete count of CpGs encapsulated in the respective region and cell type. Summary statistics denoted by 'X_cpg', 'X_pc', and 'X_var_expl' correspond to the number of CpGs per gene, number of rPCs per gene, and the variance explained by rPCs for each gene, respectively. Note: Average values are not presented as they consistently represent a single value per gene.

**Supplementary Table 5. Correlation Analysis of rPCs and Gene Complexity Metrics.** This table quantifies the correlation between regional principal components (rPCs) and various indicators of gene complexity. For each metric, 'X_corr' represents the Pearson correlation coefficient, 'X_pval' details the uncorrected p-value from the Pearson correlation test, 'mads' specifies the median absolute deviation of methylation across CpGs within the gene, 'length' indicates the gene length in base pairs, and 'cpg' denotes the count of CpGs present in the gene region. The assessment provides insights into the relationship between rPCs and the intricate structural and functional features of genes in different cell types and genomic contexts.

**Supplementary Table 6. Evaluation of Differential Methylation via Proportion Testing Across Summary Measures.** Comparative analysis of features classified as differentially methylated when summarized by regional principal components (rPCs), averages, and CpG sites. It defines the 'region_type' where methylation is summarized and ties differential methylation to specific 'traits' and 'cell_types'. For each pair of summary types compared, 'summary_type1' and 'summary_type2', the table lists the unadjusted 'pval' derived from Fisher's Tests alongside the counts of features not showing differential methylation ('not_differentially_methylatedX') and those that do ('differentially_methylatedX') for each summary type X. It further details the Bonferroni-adjusted 'adjusted_pval', the 'proportionX' of differentially methylated features to non-differentially methylated ones, the significance level of the adjusted p-value, and the corresponding percentage ('percentX').

**Supplementary Table 7. Differential Methylation Analysis Summary**. This table encapsulates the aggregate count of genes and features deemed significantly differentially methylated across varying cell and region types, utilizing different the summarization approaches, rPCs, averages, and CpGs. 'sig_genes' column tallies the unique genes marked as differentially methylated, counting each gene only once even if multiple associated features are identified. 'sig_features' quantifies all features flagged for differential methylation, which may include several per gene. The table also reports the 'total_genes' and 'total_features' tested, setting a significance threshold with a Benjamini-Hochberg corrected $p$-value of less than 0.05. This stringent criterion ensures a focus on the most substantively altered methylation patterns for subsequent biological interpretation.

**Supplementary Table 8. Integration of QTL, Fine-Mapping, and GWAS Data in Identifying meGenes**. This table provides a summary of the identified meGenes (genes with methylation levels association with at least one genomic variant) through successive layers of genomic analysis. The columns are as follows: 'X_0tested' represents the total number of features subjected to QTL mapping and GWAS integration. 'X_1qtl' indicates the count of features deemed significant from QTL mapping with an FDR threshold of $p < 0.05$. 'X_2finemapped' tallies features identified in fine-mapping with a Cluster PIP exceeding 0.5. 'X_3colocalized' shows the number of features with significant evidence of colocalization, where GLCP exceeds 0.5, noting that this is applicable only at the gene level. 'X_4ptwas' details the features significant in colocalization with stringent p-value criteria. 'X_5intact' lists features significant in the INTACT analysis with a posterior probability greater than 0.5. The table further breaks down the data by feature types, feature-variant pairs, gene-level significance, and linkage disequilibrium (LD) blocks fine-mapped from GWAS results.

**Supplementary Table 9. Distribution of Variant Cluster Sizes in GWAS Integration Steps**. This table enumerates the distribution of variant cluster sizes that were deemed significant in the integrative genomic analysis steps, specifically in colocalization and probabilistic TWAS (PTWAS). The statistics provided include mean, median, maximum, and minimum cluster sizes, offering a comprehensive view of the extent of variant clustering associated with significant results.
